# Methylome-Wide Association Study of Central Adiposity Implicate Genes Involved in Immune and Endocrine Systems

**DOI:** 10.1101/766832

**Authors:** Anne E Justice, Geetha Chittoor, Rahul Gondalia, Phillip E Melton, Elise Lim, Megan L. Grove, Eric A. Whitsel, Ching-Ti Liu, L. Adrienne Cupples, Lindsay Fernandez-Rhodes, Weihua Guan, Jan Bressler, Myriam Fornage, Eric Boerwinkle, Yun Li, Ellen Demerath, Nancy Heard-Costa, Dan Levy, James D Stewart, Andrea Baccarelli, Lifang Hou, Karen Conneely, Trevor Mori, Lawrence J. Beilin, Rae-Chi Huang, Penny Gordon-Larsen, Annie Green Howard, Kari E North

## Abstract

We conducted a methylome-wide association study to examine associations between DNA methylation in whole blood and central adiposity and body fat distribution, measured as waist circumference, waist-to-hip ratio, and waist-to-height ratio adjusted for body mass index, in 2684 African American adults in the Atherosclerosis Risk in Communities study. We validated significantly associated Cytosine-phosphate-Guanine methylation sites (CpGs) among adults using the Women’s Health Initiative and Framingham Heart Study participants (combined N=5743) and generalized associations in adolescents from The Raine Study (N=820). We identified 11 CpGs that were robustly associated with one or more central adiposity trait in adults and 2 in adolescents, including CpG site associations near *TXNIP, ADCY7, SREBF1*, and *RAP1GAP2* that had not previously been associated with obesity-related traits.

## INTRODUCTION

Elevated central adiposity is a recognized risk factor for cardiometabolic disease (CMD) [1-3]. Rates of obesity and particularly central obesity have nonetheless doubled in the U.S. over the past three decades [4-6]. Further, there are stark differences in obesity risk among minorities [4-6]. While the genetic impact on central adiposity is well-established [7], answers to several key questions could lead to important discoveries about potentially preventable contributors to obesity. One such question regards the importance of epigenetic factors in the pathogenesis of central obesity, which may point to dysregulated genomic pathways that underlie manifestation of disease [8], thus enhancing understanding of obesity-related phenotypes.

DNA methylation is an important epigenetic mechanism that links genotypes, the environment, and obesity; yet, epigenetic studies of central adiposity have been lacking [9-11]. Similarly, most of the research on DNA methylation has focused on homogeneous populations, with very few genetic studies on ancestrally diverse, admixed populations [12]. In the U.S., individuals of African American ancestry have elevated burden of hypertension, obesity, insulin resistance, impaired glucose metabolism, and ensuing CMD, compared to other U.S. populations [3, 5, 6]. Studies that interrogate obesity epigenetics utilizing multi-ethnic groups are therefore critical to gain a comprehensive understanding of the genomic architecture of these CMD related traits.

In general, studies have shown significant associations between obesity-related traits and DNA methylation [9-11]. However, some of these studies lack replication in independent samples [13], and/or use small, underpowered samples (N<200) [14]. A paucity of studies have examined the epigenetic influence on central adiposity, with only one study that considered waist circumference to height ratio, and none that adjusted for body mass index (BMI) in their associations to account for increased central adiposity exclusive of overall body size [9-13, 15, 16]. We are only beginning to understand the connections between DNA methylation and adiposity-related traits, but large samples, replication, generalization to other populations and life course stages, and consideration of potential mediators are needed to establish and further clarify the relationship between methylation and central adiposity.

We therefore conducted a methylome-wide association study (MWAS) to identify associations between DNA methylation at cytosine-phosphate-guanine sites (CpGs) in whole blood and measures of central adiposity and body fat distribution including waist circumference (WCadjBMI), WC-to-hip ratio (WHRadjBMI), and WC-to-height ratio (WCHTadjBMI) adjusted for BMI in 2,684 adult African American participants from the Atherosclerosis Risk in Communities (ARIC) study. Further, we attempted replication and generalization of significant associations across ancestrally diverse populations (N up to=5,743, 87% European American, 8% African American, and 5% Hispanic/Latino) from the Women’s Health Initiative Study and Framingham Heart Study, and European descent adolescents (N=820) from a younger generation of Australians from The Raine Study.

## MATERIALS & METHODS

### Discovery Sample

The ARIC study is a population-based prospective cohort study of cardiovascular disease risk in European American and African American individuals from four U.S. communities (Forsyth County, NC, Jackson, MS, Minneapolis, MN, and Washington county, MD) [17]. Recruitment occurred between 1987 and 1989, and participants have been followed for up to six visits across 30 years. Our discovery stage focuses on African American participants (62% women) aged 45–64 years at baseline recruited from two of the four centers (Forsyth County, NC and Jackson, MS). Anthropometrics were measured while everyone wore a scrub suit and no shoes with height recorded to the nearest centimeter (cm) and weight to the nearest pound (lb). Using a flexible tape, WC was measured at the level of the umbilicus and hip circumference (HIP) was measured at the maximum protrusion of the gluteal muscles to the nearest cm. WC and HIP are used to calculate WHR, while WC and height were used to calculate WCHT. Outliers (+/-5 SD) were removed and only WC was log transformed due to non-normality.

DNA methylation was measured in 1,702 female and 982 male African American individuals from the ARIC study from whole blood collected at visit two (1990-1992) or visit three (1993-1995). Genomic DNA was extracted from peripheral whole blood samples and bisulphite conversion of 1 µg genomic DNA was performed following standardized procedures. The Illumina HumanMethlation450 BeacChip (HM450) array (Illumina Inc.; San Diego, CA, USA) was used to quantify DNA methylation in 485,577 CpG sites. The methylation score for each CpG is reported as a beta (β) value, ranging from 0 (non-methylated) to 1 (completely methylated), according to the intensity ratio of detected methylation. Beta MIxture Quantile dilation (BMIQ) [18] was used to adjust the β values of type II design probes into a statistical distribution characteristic of type I probes. BMIQ has been shown to more effectively reduce probe set bias and technical error across replicates compared to some other peak based and quantitative normalization procedures for the HM450 array [19]. Quality control procedures used for the HM450 were published previously in Demerath et al., 2015 [11]; briefly, these procedures involved excluding individuals that were missing >1% CpGs, and excluding CpGs sites with known cross-reactive and polymorphic probes [20] and CpG sites with <1% call rate, yielding a total of 389,477 CpGs used here in discovery analysis. For those with white blood cell (WBC) count available, WBC count was assessed by automated particle counters within 24 hours of venipuncture in a local hospital hematology laboratory. To adjust methylation values for cell type proportions, we used the measured WBC type proportions for 175 participants (lymphocytes, monocytes, neutrophils, eosinophils, and basophils) to impute missing cell type proportions for the remainder of the ARIC African American participants using the Houseman method [21].

### Replication Cohorts

We attempted to replicate our significant associations from ARIC in two adult cohorts, the Framingham Heart Study (FHS) and the Women’s Health Initiative (WHI) for a combined sample size of up to 5,743 European American (N=4,977), African American (N=490), and Hispanic/Latino (N=276) adults. As our primary replication analysis, we conducted a meta-analysis combined across ancestries and both FHS and WHI. Additionally, in the interest of identifying potential ancestry-specific methylation associations, we also conducted meta-analyses stratified by ancestry. We used concordant direction of effect and trait-specific Bonferroni correction in cross-ancestry meta-analysis of FHS and WHI to determine statistically significant replication.

The Framingham Heart Study (FHS) is a prospective, longitudinal cohort study of European Americans that began enrolling participants in 1948, to identify risk factors that contribute to cardiovascular disease [22]. DNA was extracted from peripheral whole blood and DNA methylation quantified by the HM450 array at two different locations: Johns Hopkins University and the University of Minnesota. The cell type proportions we used are CD8+T cells, CD4+T cell, natural killer cells, BCell, and monocytes and they were imputed using the Houseman method and reference data [21]. Due to the batch effects, methylation beta values were adjusted for cell type proportions within each center separately, then combined for a total sample size of 3,987 individuals across both centers, including 2,469 from the offspring cohort and 1,518 from the generation 3 cohort.

WHI is a longitudinal, prospective study of post-menopausal women that began recruiting in 1993. It includes participants in the CT (Clinical Trials) and OS (Observational Study) cohorts [23]. In the present analyses, we include WHI participants from the Epigenetic Mechanisms of Particulate Matter-Mediated Cardiovascular Disease Risk (EMPC) ancillary study with available peripheral blood leukocyte DNA for methylation assay. EMPC was based on an exam site- and race/ethnicity-stratified, minority oversample (∼6%) of WHI CT participants selected from the screening, third annual, and sixth annual follow-up visits from across the 40 WHI clinical centers in the contiguous United States (N=1,755, 28% African American, 16% Hispanic/Latino, 56% European American). Baseline methylation and anthropometric measurements were considered in the current study. Weight, height, WC, and HIP were measured during a physical examination conducted at the clinical centers. Weight was measured to the nearest kg, height to the nearest cm, and WC and HIP to the nearest half cm while participants wore non-binding undergarments without shoes. WHI samples were typed using the HM450 array, BMIQ normalized and ComBat [24] adjusted. Cell type proportions (CD8+ T cell, CD4+ T cell, B cell, natural killer cell, monocyte, granulocyte) proportions were imputed using the Houseman method and reference data [21].

### Statistical Analysis

To control for potential confounding and correlated residuals due to batch effects and technical measurement error, we used linear mixed models (LMM) in R (lmer package) to adjust all CpG β values for methylation chip row as a fixed effect, chip number as a random effect, and WBC counts as fixed effects, and adjusting for kinship matrix to take into account familial correlation in FHS. Resulting methylation residuals were used in subsequent analyses. Additionally, the following variables were evaluated for inclusion as potential fixed effects in the ARIC study: 10 principal components scores (PCs) calculated from the Illumina Infinium HumanExome BeadChip genotyping to account for potential confounding by genetic ancestry, BMI, sex, age, age^2^, education, household income, current smoking status, current alcohol consumption, and physical activity. Both alcohol and physical activity were dropped from the model due to >10% missing data. The final choice of covariates was based on Bayesian model averaging (BMA) which estimates model fit for all possible combinations of covariates operating on central adiposity without methylation included, and then constructs an average weighted by the posterior model probabilities from the Bayesian information criteria (BIC) across all the possible models. Final model selection was based on each variable having a posterior inclusion probability (PIP)>40% and included in at least two of the three models with the highest observed weighted posterior probability for total model fit for any of the three traits, except for PCs which did not meet these criteria, but were included nonetheless. BMA was implemented using the R package BMS v0.3.0.

To determine if CpG site-specific β values were associated with central adiposity, measured by WCadjBMI, WHRadjBMI, and WCHTadjBMI, we implemented linear regression in R. Methylation β values were the independent variable, central adiposity was the dependent variable. All models adjusted for BMI, the PCs, study center, sex, age, highest level of education as an ordinal value, and current smoking status (nominal values: current=1, former=2, and never=3).

CpG sites with association P <1.03×10^-7^ (chip-wide significance [CWS] corrected for number of CpG variants tested, 0.05/∼458,000) for each trait in ARIC were carried forward for replication in a combined meta-analysis of WHI and FHS cohorts. All study-specific analyses were conducted stratified by self-identified race/ethnicity. Given the different analytic strategies and cell types used among the replication studies, we conducted a z-score based, sample size-weighted meta-analysis implemented in the R package EasyStrata [25] both across and within race/ethnic groups. Significant replication was asserted when regression coefficients were directionally consistent, and the meta-analyzed p value was Bonferroni-corrected significant (P<0.05/# variants tested in replication). For replication cohorts, all analytical procedures from discovery analyses were used; however, CpG β values were adjusted for estimated WBC (CD8+ T cell, CD4+ T cell, B cell, natural killer cell, monocyte, granulocyte) proportions imputed using the Houseman method and reference sample data as noted previously. We used concordant direction of effect and trait-specific Bonferroni correction to determine statistically significant replication.

### Generalization to an Adolescent Population

We were also interested in determining if adult-identified CpG-central adiposity associations were already present in a sample of European–ancestry adolescents from the West Australian Pregnancy Cohort (Raine) Study. The Raine study enrolled pregnant women ≤18 weeks gestation (1989-1991) through the antenatal clinic at King Edward Memorial Hospital and nearby private clinics in Perth, Western Australia [26, 27]. Detailed clinical assessments were performed at birth and the children were assessed longitudinally, including at 17 years of age when a blood sample, waist/hip circumference, and other anthropometric measurements were collected [28]. DNA methylation was quantified on the HM450 array. DNA methylation β values were normalized using BMIQ [18]. Analyses were carried out following the methods outlined for the adult cohorts. However, as all Raine participants were of the same age, education was not included as a covariate and smoking was indicated as only current or not current smoker.

### Nearby Genetic Associations

To determine if any of our significant CpG sites might explain previously observed genetic associations with any of the three traits, we conducted a lookup of known GWAS associations within 100 kilobases (kb) of CpGs identified in the discovery phase using the NHGRI-EBI Catalog of published genome-wide association studies (GWAS Catalog) [29]. The full list of associations in the GWAS Catalog was downloaded (v1.0, release 2019-3-22), and, using chromosome and position, all SNP associations in the catalog within the specified windows that met the genome-wide significance threshold (P<5E-8) were retained for further investigation.

### Lookup of Known Obesity-associated CpGs

To assess generalizability of previously reported obesity-related methylation sites to our study findings, we identified a total of 500 CpGs significantly associated with one or more obesity-related traits (BMI, childhood obesity, adult obesity, central obesity, BMI percentile in children, BMI change, and WC and WCHT unadjusted for BMI) in prior MWAS studies [11-13, 16, 30-37], and conducted lookups in our ARIC study results for MWAS of WCadjBMI, WHRadjBMI, and WCHTadjBMI. Of the 500 CpGs previously associated with an obesity-related trait, 408 were available in our discovery analyses for lookup. We consider statistical significance for generalizability based on Bonferroni-corrected significance for the number of sites examined (P<0.05/408=1.2×10^-4^).

## RESULTS

### Discovery

Our discovery analysis included 389,477 CpG sites and up to 2,684 ARIC participants (1,702 women and 982 men) following quality control (**Supplementary Table 1**). We identified 23 CpGs associated (P<1×10^-7^, Bonferroni corrected) with one or more waist traits (4 for WCadjBMI, 21 for WHRadjBMI, and 7 for WCHTadjBMI) (**Figure 1, Supplementary Table 2, Table 1**). As in previous MWAS studies of obesity-related traits, our results show evidence of inflation [30, 32] (**Supplementary Figure 1)**; however, since genomic control correction may be overly conservative [38], we rely on replication to protect against Type 1 error and use strict Bonferroni-corrected significance for selection of variants for replication. While the current study examines central adiposity or body fat distribution associations with methylation after adjustment for overall body size (BMI), five of the significant CpGs have been associated with other obesity-related traits, including BMI. All five have been previously associated with a central adiposity trait, while 18 are newly identified associations for any obesity-related anthropometric measure. The novel central adiposity associations include cg19693031 near *TXNIP* and cg16778018 in *MGRN1*, which were significantly associated with all three traits (WCadjBMI, WHRadjBMI, and WCHTadjBMI) (**Supplementary Table 2**).

**Figure 1.**
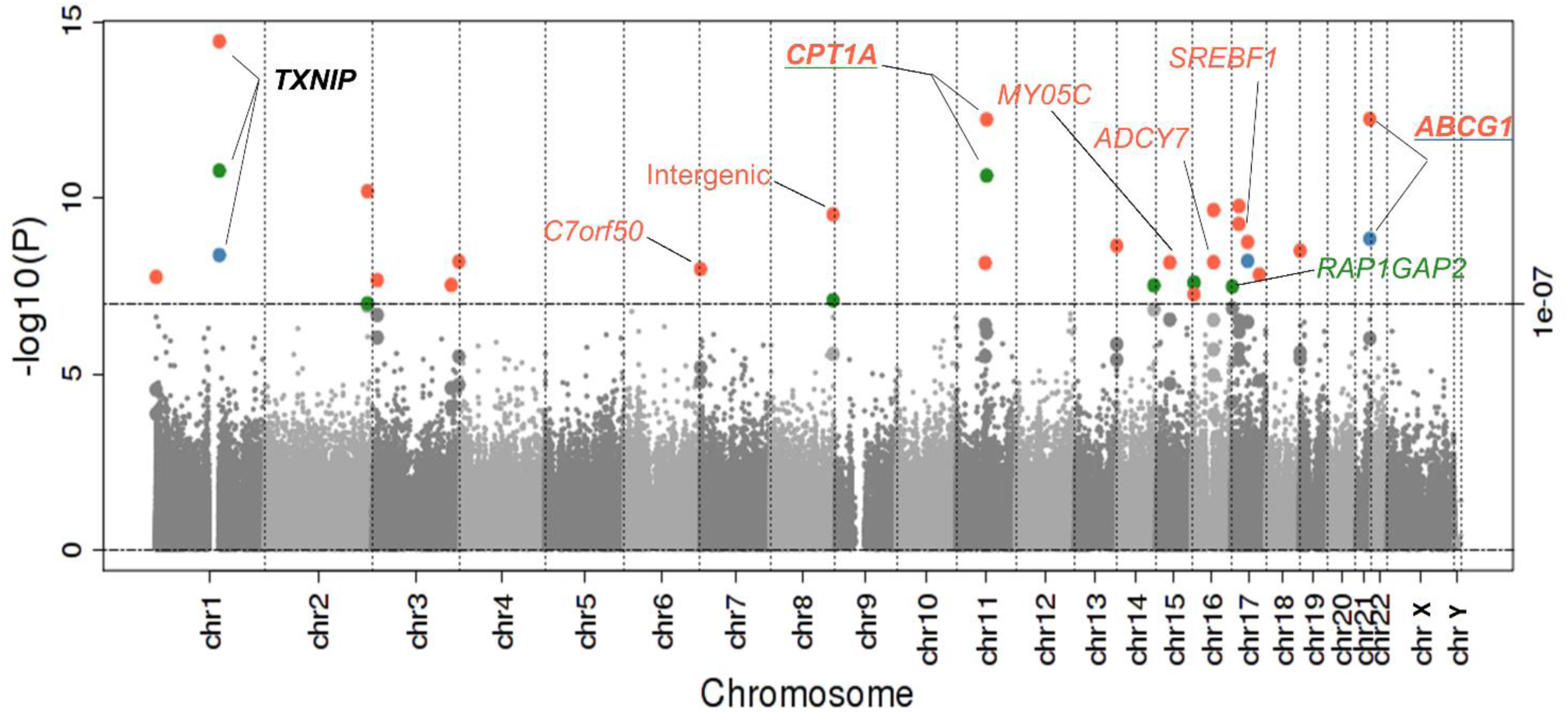
Manhattan plot of association results in the ARIC study discovery analyses. CpGs associated with WCadjBMI are in blue, WHRadjBMI in orange, and WCHTadjBMI in green. All replicated CpGs are annotated with their nearest gene. Gene names are bolded if representing a CpG associated with more than one central adiposity trait. *TXNIP* was associated with all three traits and is highlighted in black.

### Multiethnic Replication Analysis

Eleven CpGs in nine gene regions met significance criteria (consistent direction of effects and Bonferroni significance) for replication in the multiethnic meta-analysis of WHI and FHS (**Table 1, Supplementary Table 2, Figure 2**). Of the four CpGs associated with WCadjBMI in the discovery analysis, two CpGs replicated, cg19693031 near *TXNIP* a novel CpG, and cg06500161 within *ABCG1* that was previously associated with BMI and WC unadjusted for BMI [11, 30-34]. Of the 21 CpGs associated with WHRadjBMI brought forward for replication, 10 replicated, including both CpGs that replicated for WCadjBMI; as well as CpGs associated with other obesity-related traits cg00574958 near *CPT1A* [11, 12, 16, 30-32, 34, 35], cg04816311 in *C7orf50* [11, 31], cg06192883 in *MY05C* [11, 30, 31]; and novel obesity-related associations for cg23580000 and cg06897661 in *ADCY7*, cg15863539 and cg20544516 in *SREBF1*, and an intergenic CpG, cg26610247, on chromosome 8. For WCHTadjBMI, three of the seven CpGs brought forward from the discovery stage replicated, including cg19693031 (*TXNIP*), associated with both WCadjBMI and WHRadjBMI, cg00574958 (*CPT1A*), also significant for WHRadjBMI, and cg12537003 in *RAP1GAP2*, which has not been previously associated with any adiposity trait.

**Table 1.**
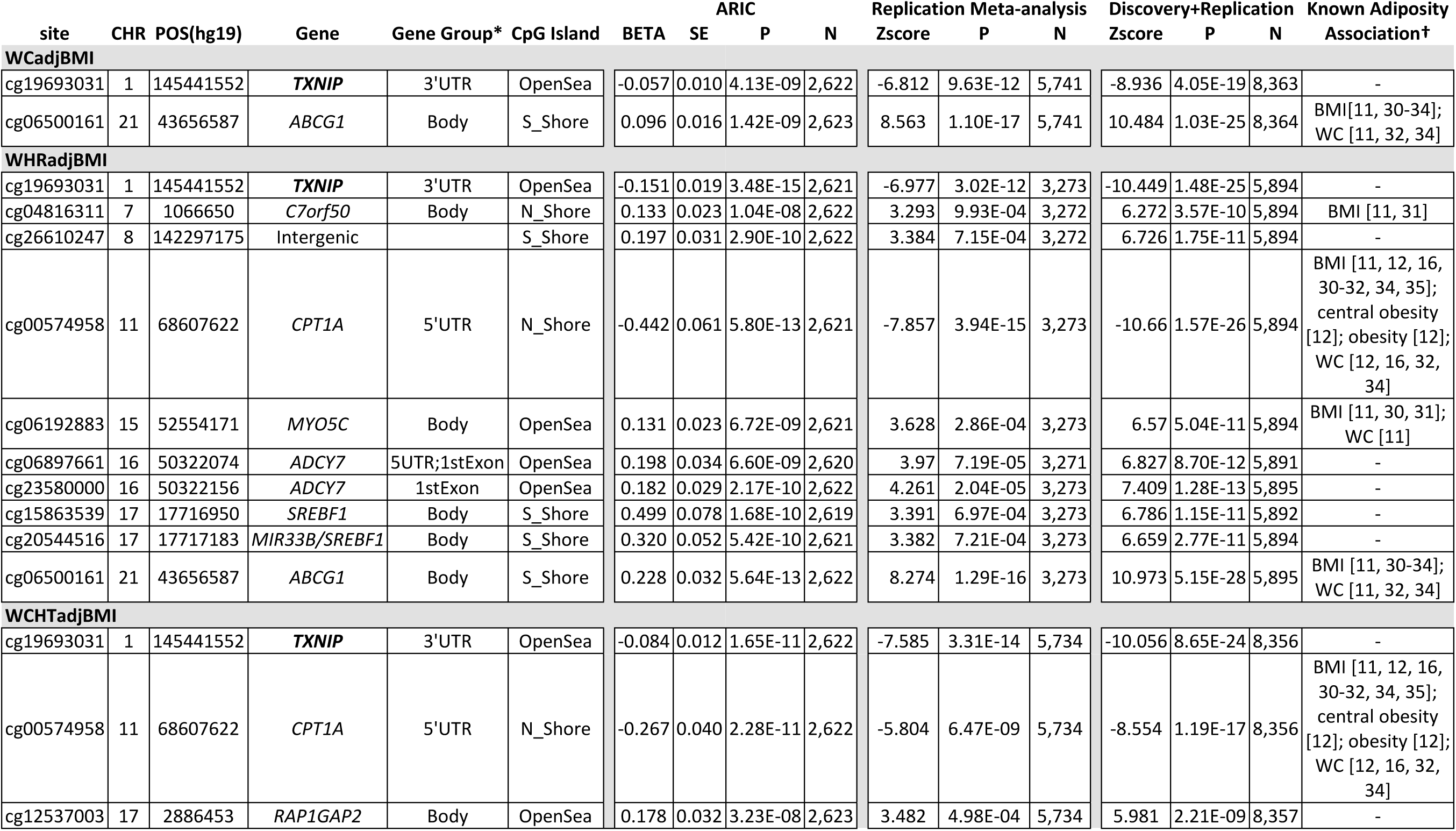
Association results for CpGs significantly associated with central adiposity and body fat distribution in the discovery and replication analyses. *Note that multiple GeneRef Groups denote location respective to CpG sites. New annotation only provided when ref location or ref gene changes. †For any CpG with a known association with an obesity trait, we list the trait and reference. Gene names are bolded if they are significantly associated with all three traits.

| site | CHR | POS(hg19) | Gene | Gene Group* | CpG Island | BETA | SE | ARIC<br>P | N | Replication<br>Zscore | Meta-analysis<br>P | N | Discovery+Replication<br>Zscore | P | N | Known Adiposity<br>Association† |
| --- | --- | --- | --- | --- | --- | --- | --- | --- | --- | --- | --- | --- | --- | --- | --- | --- |
| **WCadjBMI** |  |  |  |  |  |  |  |  |  |  |  |  |  |  |  |  |
| cg19693031 | 1 | 145441552 | **TXNIP** | 3'UTR | OpenSea | -0.057 | 0.010 | 4.13E-09 | 2,622 | -6.812 | 9.63E-12 | 5,741 | -8.936 | 4.05E-19 | 8,363 | - |
| cg06500161 | 21 | 43656587 | **ABCG1** | Body | S_Shore | 0.096 | 0.016 | 1.42E-09 | 2,623 | 8.563 | 1.10E-17 | 5,741 | 10.484 | 1.03E-25 | 8,364 | BMI[11, 30-34];<br>WC [11, 32, 34] |
| **WHRadjBMI** |  |  |  |  |  |  |  |  |  |  |  |  |  |  |  |  |
| cg19693031 | 1 | 145441552 | **TXNIP** | 3'UTR | OpenSea | -0.151 | 0.019 | 3.48E-15 | 2,621 | -6.977 | 3.02E-12 | 3,273 | -10.449 | 1.48E-25 | 5,894 | - |
| cg04816311 | 7 | 1066650 | **C7orf50** | Body | N_Shore | 0.133 | 0.023 | 1.04E-08 | 2,622 | 3.293 | 9.93E-04 | 3,272 | 6.272 | 3.57E-10 | 5,894 | BMI [11, 31] |
| cg26610247 | 8 | 142297175 | Intergenic |  | S_Shore | 0.197 | 0.031 | 2.90E-10 | 2,622 | 3.384 | 7.15E-04 | 3,272 | 6.726 | 1.75E-11 | 5,894 | - |
| cg00574958 | 11 | 68607622 | **CPT1A** | 5'UTR | N_Shore | -0.442 | 0.061 | 5.80E-13 | 2,621 | -7.857 | 3.94E-15 | 3,273 | -10.66 | 1.57E-26 | 5,894 | BMI [11, 12, 16,<br>30-32, 34, 35];<br>central obesity<br>[12]; obesity [12];<br>WC [12, 16, 32,<br>34] |
| cg06192883 | 15 | 52554171 | **MYO5C** | Body | OpenSea | 0.131 | 0.023 | 6.72E-09 | 2,621 | 3.628 | 2.86E-04 | 3,273 | 6.57 | 5.04E-11 | 5,894 | BMI [11, 30, 31];<br>WC [11] |
| cg06897661 | 16 | 50322074 | **ADCY7** | 5UTR;1stExon | OpenSea | 0.198 | 0.034 | 6.60E-09 | 2,620 | 3.97 | 7.19E-05 | 3,271 | 6.827 | 8.70E-12 | 5,891 | - |
| cg23580000 | 16 | 50322156 | **ADCY7** | 1stExon | OpenSea | 0.182 | 0.029 | 2.17E-10 | 2,622 | 4.261 | 2.04E-05 | 3,273 | 7.409 | 1.28E-13 | 5,895 | - |
| cg15863539 | 17 | 17716950 | **SREBF1** | Body | S_Shore | 0.499 | 0.078 | 1.68E-10 | 2,619 | 3.391 | 6.97E-04 | 3,273 | 6.786 | 1.15E-11 | 5,892 | - |
| cg20544516 | 17 | 17717183 | **MIR33B/SREBF1** | Body | S_Shore | 0.320 | 0.052 | 5.42E-10 | 2,621 | 3.382 | 7.21E-04 | 3,273 | 6.659 | 2.77E-11 | 5,894 | - |
| cg06500161 | 21 | 43656587 | **ABCG1** | Body | S_Shore | 0.228 | 0.032 | 5.64E-13 | 2,622 | 8.274 | 1.29E-16 | 3,273 | 10.973 | 5.15E-28 | 5,895 | BMI [11, 30-34];<br>WC [11, 32, 34] |
| **WCHTadjBMI** |  |  |  |  |  |  |  |  |  |  |  |  |  |  |  |  |
| cg19693031 | 1 | 145441552 | **TXNIP** | 3'UTR | OpenSea | -0.084 | 0.012 | 1.65E-11 | 2,622 | -7.585 | 3.31E-14 | 5,734 | -10.056 | 8.65E-24 | 8,356 | - |
| cg00574958 | 11 | 68607622 | **CPT1A** | 5'UTR | N_Shore | -0.267 | 0.040 | 2.28E-11 | 2,622 | -5.804 | 6.47E-09 | 5,734 | -8.554 | 1.19E-17 | 8,356 | BMI [11, 12, 16,<br>30-32, 34, 35];<br>central obesity<br>[12]; obesity [12];<br>WC [12, 16, 32,<br>34] |
| cg12537003 | 17 | 2886453 | **RAP1GAP2** | Body | OpenSea | 0.178 | 0.032 | 3.23E-08 | 2,623 | 3.482 | 4.98E-04 | 5,734 | 5.981 | 2.21E-09 | 8,357 | - |

### Ancestry Stratified Analyses

We conducted ancestry-specific analyses to identify any potential population-specific associations (**Supplementary Figure 2, Supplementary Table 3**). Among all traits considered, 72% of all trait-CpG associations were directionally consistent across all ancestries. Three CpG-trait associations met Bonferroni-corrected significance in the WHI-only African Americans (cg06500161 with WCadjBMI and WHRadjBMI; and cg00574958 with WHRadjBMI), all of which were significant in the multiethnic replication analysis. For meta-analyses of African-Americans from WHI and ARIC together, 24 of the trait-CpG associations remained chip-wide significant (P<1.03×10^-7^). Of these 24 trait-CpG associations, nine did not meet the required significance threshold for replication in the combined meta-analysis, including cg16778018 in *MGRN1* and cg00994936 in *DAZAP1*, which were nominally associated (P<0.05) with WHRadjBMI only in the WHI African Americans. For European-descent meta-analyses (WHI + FHS), 11 trait-CpG associations were significant after multiple test correction (two for WCadjBMI, six for WHRadjBMI, and three for WCHTadjBMI); all of which were significant in the multiethnic replication analysis. For the WHI Hispanic/Latino-specific analysis, two associations met significance criteria, including cg06500161 for WCadjBMI, and cg26610247 for WCHTadjBMI, which did not reach significance in the multiethnic replication analysis. A lack of generalization for many of the sites may be the result of reduced power due to sample size, especially in Hispanic/Latinos (N=276).

### Generalization to an Adolescent Population

We examined CpG-trait associations from our adult discovery analysis in The Raine Study’s Generation 2 cohort at the 17 year follow up to test for generalization (**Table 2, Supplementary Table 2**). Two CpGs met our criteria for statistical significance, cg00994936 in *DAZAP1* associated with WHRadjBMI, and cg19693031 downstream of *TXNIP*, associated with WCadjBMI and WHRadjBMI. While cg19693031 replicated in our adult meta-analyses, cg00994936 was only nominally significant and directionally significant in our adult meta-analysis (P=4.23E-3). Overall, 21 (66%) of the 32 CpG-trait associations were directionally consistent with the discovery analyses, of which 10 (48%) reached nominal significance (P<0.05). For WCadjBMI, two CpGs displayed nominally significant associations, including cg19693031, which remained significant following multiple-test correction. For WHRadjBMI, 11 CpGs were nominally associated, including the two that remained significant after multiple test correction. Only one CpG site, cg26610247, was nominally significant for WCHTadjBMI, but did not meet the Bonferroni significance threshold.

**Table 2.**
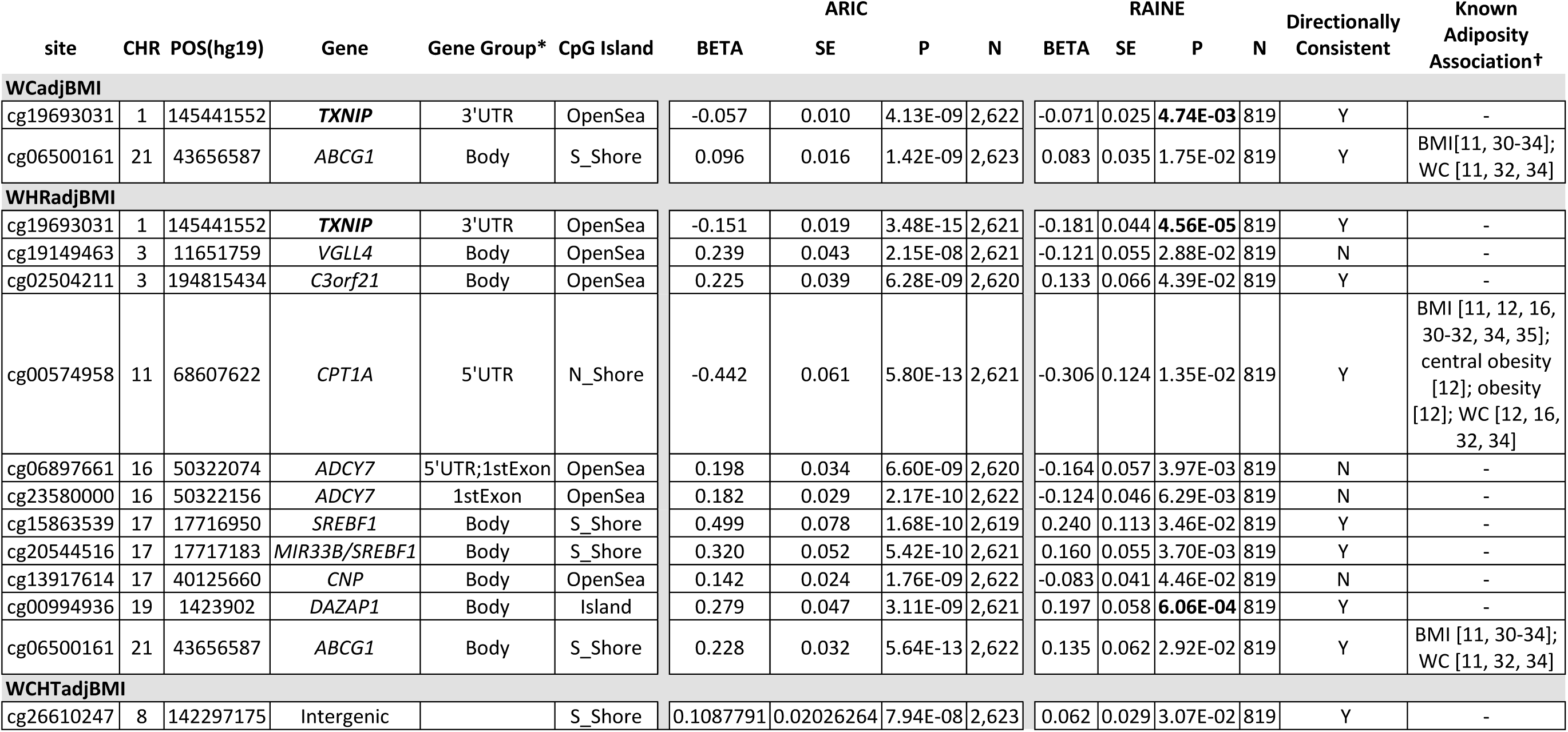
Association results for CpGs significantly associated with central adiposity and body fat distribution in the discovery and nominally associated in The Raine Study. P-values that meet Bonferroni-corrected significance are bolded. *Note that multiple GeneRef Groups denote location respective to CpG sites. New annotation only provided when ref location or ref gene changes. †For any CpG with a known association with an obesity trait, we list the trait and reference. Gene names are bolded if they are significantly associated with all three traits.

### Nearby Genetic Associations

We conducted a search in the NHGRI-EBI GWAS Catalog[39] to determine if any of our significant CpG sites were nearby (<100 MB) genetic variants associated with traits or diseases of interest (**Supplementary Table 4**). We identified several cardiometabolic (lipid levels, C-reactive protein [CRP], blood pressure measures, BMI, birth weight, type 2 diabetes *etc.*), and blood cell (e.g. mean corpuscular volume, hemoglobin, WBC counts, etc.), and other (i.e. height, bone mineral density, etc.) traits potentially related to obesity with GWAS associations near one or more of our WHRadjBMI-associated CpG sites. Only one CpG associated with WCHTadjBMI had nearby associations present in the GWAS Catalog (cg12537003), but these did not include any relevant obesity or cardiometabolic traits. No associations were found nearby WCadjBMI sites.

### Known Obesity-related CpG Associations

Of the 408 CpGs previously associated with an obesity-related trait available for look-up, 35 CpGs were significantly associated (P<1.23E-4) with one or more central adiposity traits in ARIC African Americans (**Figure 2, Supplementary Table 5**). Among these were cg06500161 in *ABCG1* and cg06192883 in *MYO5C* both previously associated with BMI and WC unadjusted for BMI; cg00574958 upstream of *CPT1A* previously associated with BMI, WC unadjusted for BMI, obesity, and central obesity; and cg04816311 in *C7orf50* previously associated with BMI. All four of these were significantly associated with WHRadjBMI in the current analysis, cg06500161 is also associated with WCadjBMI; and cg00574958 is also associated with WCHTadjBMI. Thus, while there are some trait-specific associations with DNA methylation, there is strong evidence for overlap in the influence of DNA methylation across obesity-related traits.

**Figure 2.**
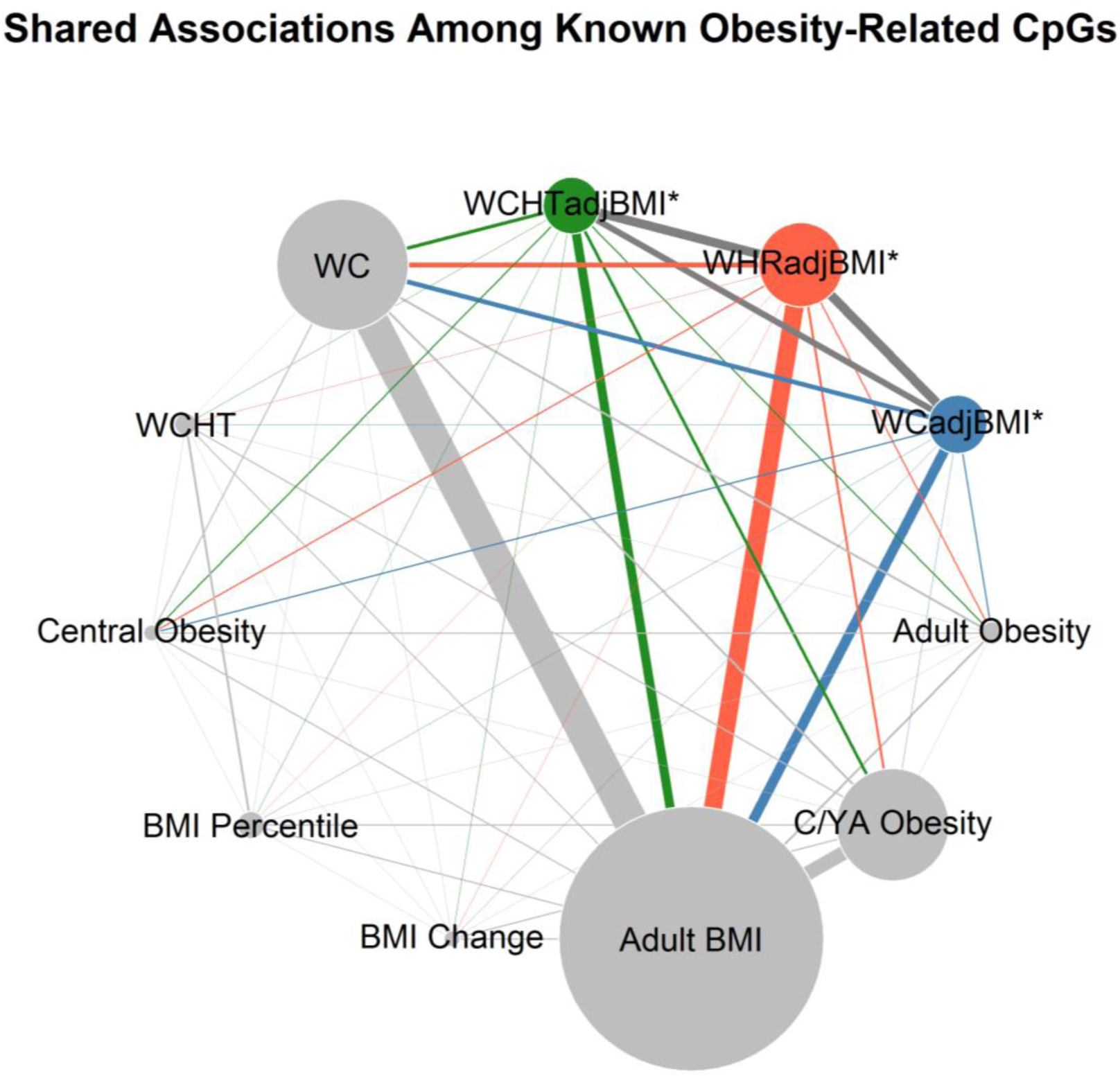
Bubble chart representing the overlap between known adiposity-associated CpGs. Each grey bubble represents a trait reported in the literature for which we conducted a look-up of previously-reported CpGs in our discovery analysis results. Focus traits for the current paper are shown in blue (WCadjBMI), orange (WHRadjBMI), and green (WCHTadjBMI). The size of the bubble is proportional to the number of statistically significant CpGs reported for that trait (array-wide significance for traits from literature, Bonferroni-significance in the lookups within the current analysis). Each pair of bubbles is connected by a line that is proportional to the number significant loci that overlap among the traits. Abbreviations not noted elsewhere: C/YA Obesity – childhood and young adult obesity.

## DISCUSSION

We identified 11 CpGs that were robustly associated with one or more central adiposity traits in adults. Seven of these CpGs are novel central adiposity associations (near *ADCY7, SREBF1, RAP1GAP2*, and one intergenic), and cg19693031 near *TXNIP*, which is apparent across age and cohort. We additionally identified four CpG – central adiposity associations that were apparent across age and cohort, yet these effects were previously published for other obesity-related traits, such as BMI [11, 12, 16, 30-32, 34, 35], central obesity [12], overall obesity [12], and WC unadjusted for BMI [12, 16, 32, 34].

Only a small number of studies have been published showing obesity-related variation in DNA methylation [9-13, 16, 30-34, 36, 37], with most studies generally using either global methylation or candidate gene approaches, with little replication in either independent samples [13] or small to underpowered samples. Fewer studies have examined the epigenetic influence on central adiposity-related traits and none reported adjusting for overall adiposity, in an attempt to identify CPG sites specific to central adiposity. Many of the associations identified in African Americans generalize to European Americans, yet a lack of generalization in Hispanic/Latinos is likely the result of smaller sample size in this stratum as the majority of CpG associations were directionally consistent and of similar magnitude of effect. Most trait-CpG associations were directionally consistent in The Raine Study adolescent cohort, yet all but one displays a larger magnitude of effect in adults.

### Independence of Methylation Effects from Established GWAS Hits for Central Obesity

Notably, there were no nearby GWAS associations with any central adiposity and body fat distribution traits, indicating that our observed CpG associations are not likely the result of confounding by cis-acting SNPs associated with WCadjBMI, WHRadjBMI, or WCHTadjBMI. However, given the large number of SNP-trait associations with white blood cell counts near cg12537003 associated with WCHTadjBMI; and cg26610247, cg22348356, and cg15863539 associated with WHRadjBMI, it is possible that these CpG-trait associations are the result of residual confounding due to WBC proportions. Further analyses in other obesity-relevant tissues is warranted to confirm this finding.

### Biological Interrogation of Nearby Genes Implicate a Role in the Neuroendocrine System

Many of the identified CpGs associated with central adiposity are in or near likely candidate genes (including *TXNIP, SREBF1, MIR33B, MYO5C, C7orf50, ABCG1*, and *CPT1A)* that are important to lipid and/or glucose homeostasis and are highly expressed in tissues within the neuroendocrine system and/or whole blood. Also, methylation at many of these sites is associated with other related cardiometabolic traits. For example, increased methylation at cg19693031 in the 3’ UTR (untranslated region) of *TXNIP* is negatively associated with all three traits (WCadjBMI, WHRadjBMI, and WCHTadjBMI). Methylation at this CpG has been negatively associated with hepatic steatosis [40], prevalent and incident type 2 diabetes (T2D) [41, 42], fasting blood glucose, HOMA-IR (Homeostatic model assessment-insulin resistance) [42], systolic and diastolic blood pressure (SBP and DBP) [43], triglyceride levels (TG) [44], and chylomicrons (Type A) [45]. Methylation at cg19693031 is also positively associated with expression of *TXNIP* in liver tissue [41]. Interestingly, there are no cardiometabolic GWAS associations for T2D near this CpG and no cis-meQTL (methylation quantitative locus) either [43]. However, recent investigations implicate cg19693031 methylation as a mediator between early life famine and adult metabolic disease [46], indicating the relationship between methylation at this site and cardiometabolic disease may be driven through environmental exposures. *TXNIP* is ubiquitously expressed, but exhibits the highest expression in subcutaneous adipose tissue in the Genotype-Tissue Expression (GTEx) database [47].

We identified two new CpGs (cg15863539 and cg20544516) associated with WHRadjBMI in *SREBF1* (sterol regulatory element binding transcription factor 1). Intergenic CpGs near *SREBF1* have been associated with BMI and WC unadjusted for BMI in previous investigations [11]. *SREBF1* exhibits the highest expression in the adrenal glands and to a lesser degree in the salivary and pituitary glands [47]. This gene is well-studied and has been shown to be integral to insulin-dependent cholesterol synthesis and lipid homeostasis [48, 49]. Cg20544516 is also located within *MIR33B* (microRNA 33b). miRNAs are non-coding short RNAs that affect both translational capability and stability of mRNAs by playing a key role in post-transcriptional regulation of gene expression. DNA methylation at these sites within miRNA-33B is associated with blood lipid levels [44, 50]. Further, plasma microRNA 33b levels were found to be associated with lipid disorders [51], especially in T2D patients with dyslipidemia [52].

Additionally, cg04816311 in *C7orf50* (Chromosome 7 Open Reading Frame 50), associated with increased WHRadjBMI in the current study, has been positively associated with BMI [11] and T2D in a Sub-Saharan African population [53] even after adjusting for BMI. GWAS associations exist nearby for total cholesterol (TC), low-density lipoprotein cholesterol (LDL), and pleiotropy between CRP and TC [54, 55]. C7orf50 exhibits the highest expression in the pituitary gland but is also expressed in the thyroid [47]. Methylation at cg04816311 is negatively and significantly associated with expression of *C7orf50* and with *GPER* in monocytes (MESA EpiGenomics eMS Database [56]). *GPER* (G protein-coupled estrogen receptor 1) is a well-studied gene that is important for the estrogen-dependent stimulation of multiple signaling pathways, thus plays a major role in several cellular processes and biological functions, especially cardiometabolic functions such as glucose and lipid homeostasis [57], regulation of blood pressure [58], lean and fat mass [57, 59] as illustrated in murine models. *GPER* is expressed at low levels across several tissue types but exhibits the highest expression in the stomach followed by EBV-transformed lymphocytes, tibial nerve, and thyroid in the GTEX database [47]. Other investigations have shown that *GPER* is also highly expressed in the hypothalamus [60].

Methylation at cg00574958 in the 5’ region of *CPT1A* (carnitine palmitoyltransferase 1A), associated with WHRadjBMI and WCHTadjBMI in the current study, has been previously associated with a number of cardiometabolic traits, including BMI, obesity, central obesity, WC unadjusted for BMI, lipid levels, lipoproteins, blood pressure, T2D, and plasma adiponectin [11, 12, 16, 43, 53, 61-64]. Also, a recent study has shown that blood pressure is causally associated with methylation at this site and methylation at this CpG influences expression of *CPT1A* [43]. CPT1a is the hepatic isoform of CPT 1 that together with CPT2 (carnitine palmitoyltransferase 2) initiates the mitochondrial oxidation of long-chain fatty acids [65, 66]. CPT 1 is the vital enzyme in the carnitine-dependent transport across the inner membrane of mitochondria and reduced fatty acid beta-oxidation rate results in its deficiency. Similarly, mutations in this gene are associated with altered fatty acid oxidation and cancer [67, 68]. *CPT1A* is highly expressed in transverse colon followed by aorta in the GTEx database [47]. *CPT1A* is regulated by PPARα, which is a ligand for drugs used in treating cardiovascular disease (CVD) [64]. The combined evidence, including the current study, indicates a strong epigenetic role of *CPT1A* in metabolic dysfunction.

Methylation at cg06500161 in *ABCG1* (ATP binding cassette (*ABC*) subfamily G member 1) is associated with WHRadjBMI and WCadjBMI. *ABC* genes are members of the superfamily of ABC transporters that are involved in macrophage cholesterol, phospholipids transport, and in extra and intra cellular lipid homeostasis [69]. Mutations and increased methylation in this gene are also associated with metabolic syndrome [15, 69], T2D [70, 71], CVD [71], and lipids [50, 63]. Additionally, studies reported alterations involving the cholesterol metabolism gene network (including *ABCG1*) are associated with molecular mechanisms of obesity/inflammation and T2D [71], complementing the association of CpG sites annotated to *ABCG1* with adiposity measures in this study. Moreover, *ABCG1* is highly expressed in adrenal gland and the spleen [47].

Cg06192883 in *MYO5C* (myosin VC) is associated with WHRadjBMI in the current analysis. Methylation at this site has been previously associated with BMI [11, 30], WC unadjusted for BMI [11], obesity [72], CRP [55], glycan [73], and glycine [45]. *MYO5C* is important for actin binding and likely functions to selectively bundle and transport secretory vesicles. Thus, *MYO5C* is involved in several cellular processes [74, 75], including glucose uptake into muscle cells in response to insulin [76]. *MYO5C* is highly expressed in the thyroid, cerebellum, salivary glands, and subcutaneous adipose tissue, among other tissues [47].

### Prioritization of Genes Involved in Inflammatory Response

In addition to candidate regions with genes related to cardiometabolic disease, several regions harbor genes related to inflammation. For example, cg06897661 and cg23580000 in *ADCY7* (5’UTR/first exon, and first exon, respectively) are associated with WHRadjBMI in the current study. cg23580000 has been associated with obesity in a candidate gene study [77] whereby methylation sites in genes involved in lipolysis were investigated for association with obesity in 15 obese cases and 14 controls; however these findings were never replicated. *ADCY7* (adenylate cyclase 7) is necessary for cyclic AMP synthesis in cells important for human immune response (e.g. macrophages, T cells, B cells) [78, 79], and thus is vital to regulating proinflammatory responses. Mice deficient in Adcy7 exhibit reduced cAMP, B-cell and T-cell production, are more prone to endotoxic shock, and overall display a weakened immune response [78].

Also, cg26610247 associated with WHRadjBMI in the current paper has been positively associated with CRP in whole blood, and has nearby GWAS associations with DBP [80], SBP [81, 82], birth weight [83], and blood cell traits [81, 84]. While this CpG is intergenic and not adjacent to any CpG islands, methylation at cg26610247 is significantly associated with decreased expression of *PTP4A3* in monocytes (MESA EpiGenomics eMS Database [56]). Cg12537003 in *RAP1GAP2* (RAP1 GTPase activating protein 2) is associated with WCHTadjBMI. This gene plays a role in platelet aggregation [85] and release of granulocytes from platelets in human cells [86]. There are multiple GWAS associations with blood cell traits nearby this CpG [84].

## ACKNOWLEDGEMENTS

**The ARIC Study.** The authors thank the staff and participants of the ARIC study for their important contributions. **The Raine Study.** The authors are grateful to The Raine Study participants and their families, and The Raine Study management team for cohort co-ordination and data collection, the National Health & Medical Research Council (NHMRC) for their long-term contribution to funding the study over the last 29 years and The Telethon Kids Institute for long term support of the Study. We also acknowledge The University of Western Australia (UWA), Raine Medical Research Foundation, The Telethon Kids Institute, Women and Infants Research Foundation, Edith Cowan University, Murdoch University, The University of Notre Dame Australia, Raine Medical Research Foundation, and Curtin University for providing funding for Core Management of The Raine Study. For Raine, the DNA methylation work was supported by NHMRC grant 1059711. Collaborative analyses are supported by NHMRC 1142858. Data collection and biological specimens at the 17-year follow-up were funded by the NHMRC Program Grant ID 353514 and Project Grant #403981. RCH is supported by NHMRC Fellowship grant number 1053384. This work was supported by resources provided by The Pawsey Supercomputing Centre with funding from the Australian Government and the Government of Western Australia**. The WHI Study**. The WHI program is supported by contracts from the National Heart, Lung and Blood Institute, NIH. The authors thank the WHI investigators and staff for their dedication, and the study participants for making the program possible. A listing of WHI investigators can be found at http://www.whi.org/researchers/Documents%20%20Write%20a%20Paper/WHI%20Investigator%20Short%20List.pdf.

## FUTURE PERSPECTIVE

We present results from a large, well-powered study examining the relationship between DNA methylation and measures of central adiposity exclusive of overall body size. We identify several new CpG sites and performed replication to confirm CpG-trait associations in adults. Additionally, we generalize two CpGs associated with WHRadjBMI, and WCadjBMI and WHRadjBMI in a predominantly Europeandescent cohort of late adolescents. We also show that the large majority of associations generalize across populations based on self-identified ancestry. These newly implicated CpGs associated with central adiposity traits are in genes or are known to influence expression of genes active in the neuroendocrine system, which function to regulate lipid and glucose homeostasis, as well as genes involved in inflammatory response. Our discovery of 11 methylation sites associated with anthropometric measures of central adiposity and body fat distribution may help to explain differences in central adiposity and/or in downstream cardiometabolic disease risk, like T2D, hyperlipidemia, and hypertension. Taken together, our results highlight the potential importance of lipid and glucose homeostasis, inflammatory response, and metabolism in the relationship between gene methylation and body fat distribution and identify potentially causal genes underlying these relationships. Notably, we identify several genes that have not been implicated in previous GWAS or MWAS of obesity-related traits. Thus, our findings offer potential new therapeutic targets for exploration in studies of risks associated with abdominal fat accumulation and advancement of our understanding of the epigenetic architecture and underlying biology of central adiposity. A minor limitation to the current analyses is the inability to account for some potential mediators that may play a role in both central adiposity and methylation (i.e. physical activity and alcohol consumption). Another limitation to the current study is reverse causation. While we model our associations with adiposity measures as the outcome of interest and methylation as a predictor, adiposity may also affect methylation. Future investigations - preferably with longitudinally assessed methylation and adiposity, and concurrently measured gene expression - are therefore needed to evaluate this possibility and then identify the causal pathways and their downstream cardiometabolic and inflammatory consequences.

