## Supplemental_Materials for "Methylome-Wide Association Study of Central Adiposity Implicate Genes Involved in Immune and Endocrine Systems"

**Supplementary Figure 1.** QQ plot and GC lambda for WCadjBMI (blue), WHRadjBMI (orange), and WCHTadjBMI (green) discovery analyses.

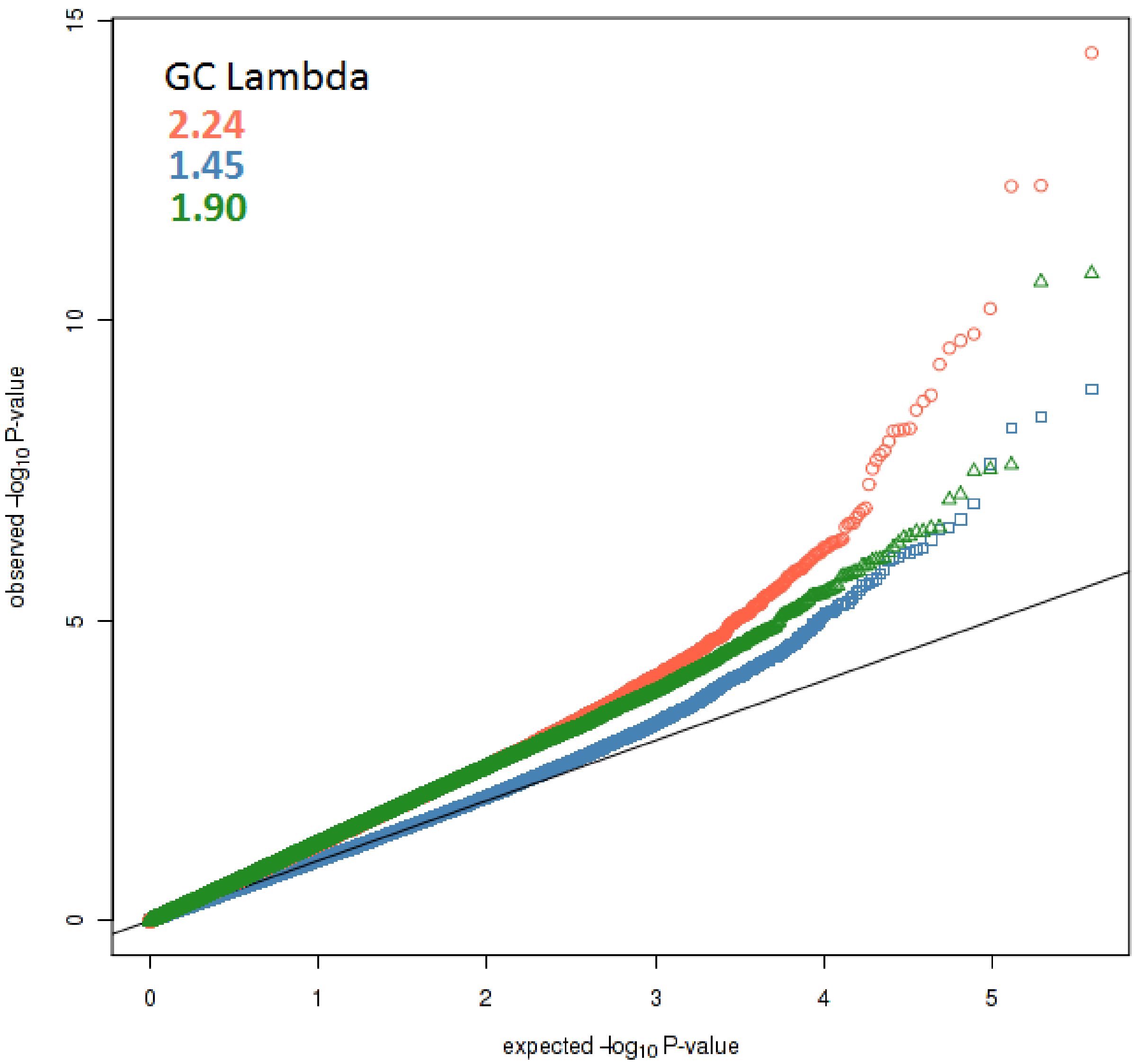

**Supplementary Figure 2**. Heatmaps summarizing the ancestry-specific association results for significant CpGs. Each heatmap square is shaded based on the trait-CpG association z-score.

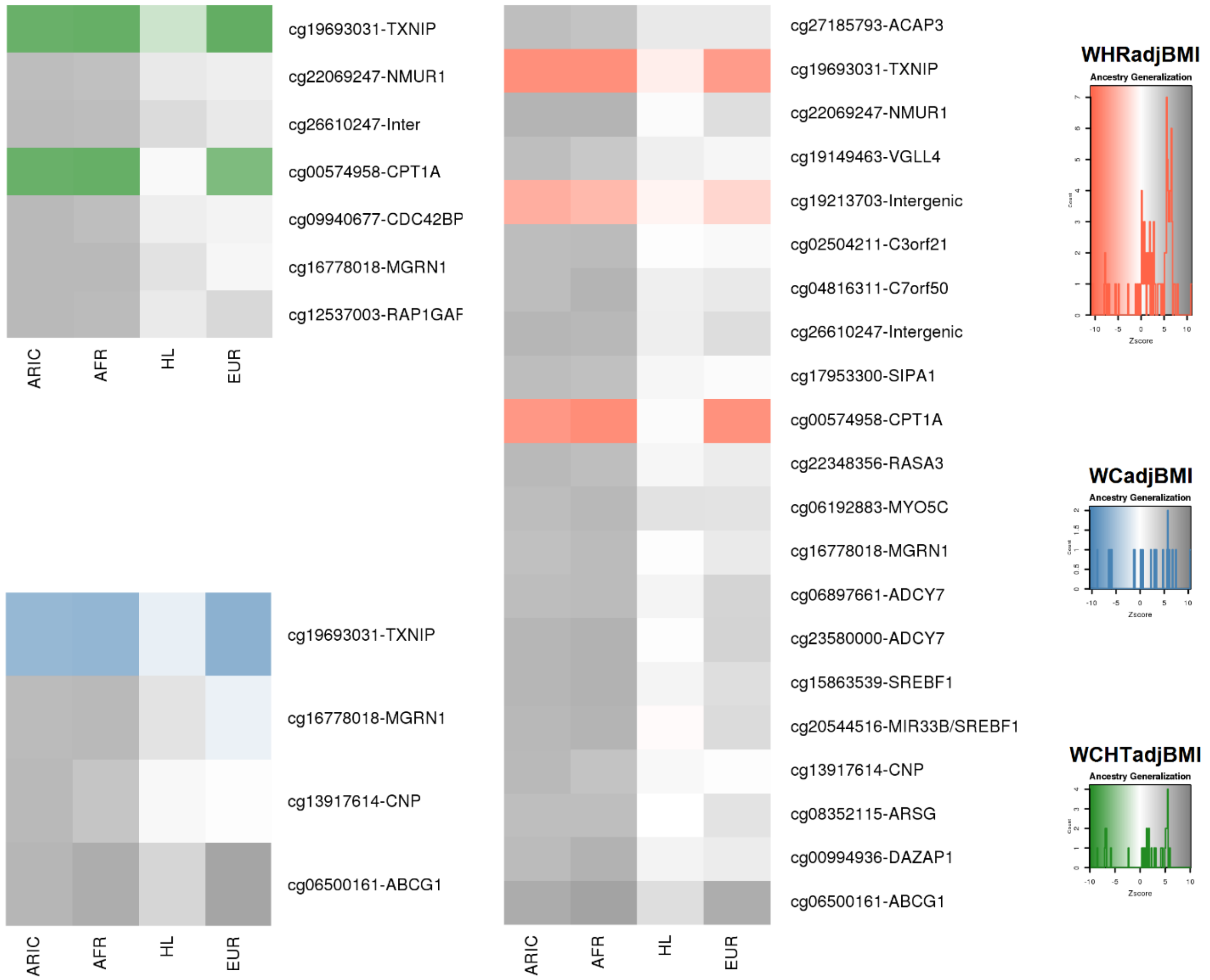

**Supplementary Table 1**. Descriptive statistics for all participating studies. *Sample size for women is 795 and for men is 723.

**Discovery Population**

| | ARIC | | | |
|---|---|---|---|---|
| | Women | | Men | |
| | N = 1,702 | | N = 982 | |
| | Mean | SD | Mean | SD |
| **Age (years)** | 56.51 | 5.81 | 56.79 | 6.00 |
| **WC (cm)** | 102.85 | 16.40 | 99.33 | 13.16 |
| **Weight (kg)** | 183.04 | 39.87 | 192.58 | 38.11 |
| **Height (cm)** | 163.17 | 6.10 | 176.19 | 6.77 |
| **BMI (kg/m$^2$)** | 31.26 | 6.59 | 28.15 | 5.05 |
| **WHR (cm/cm)** | 0.91 | 0.08 | 0.95 | 0.05 |
| **Hip (cm)** | 112.22 | 13.01 | 104.39 | 10.39 |
| **WCHT (cm/cm)** | 0.63 | 0.10 | 0.56 | 0.07 |
| | **N** | **%** | **N** | **%** |
| - | - | - | - | - |
| **Grade school or 0 years education** | 276 | 16.25 | 232 | 23.72 |
| - | - | - | - | - |
| **High school, but no degree** | 385 | 22.67 | 177 | 18.10 |
| **High school graduate** | 383 | 22.56 | 191 | 19.53 |
| **Vocational school** | 107 | 6.30 | 68 | 6.95 |
| - | - | - | - | - |
| **College** | 303 | 17.84 | 175 | 17.89 |
| **Graduate school or Professional school** | 244 | 14.37 | 135 | 13.80 |
| - | - | - | - | - |
| - | - | - | - | - |
| **Current Smoker** | 330 | 19.53 | 329 | 33.88 |
| **Former Smoker** | 417 | 24.67 | 398 | 40.99 |
| **Never Smoker** | 943 | 55.80 | 244 | 25.13 |

**Replication Population**

| | FHS - European American | | | |
|---|---|---|---|---|
| | Women | | Men | |
| | N = 2140 | | N = 1847 | |
| | Mean | SD | Mean | SD |
| **Age (years)** | 58.51 | 13.32 | 58.18 | 13.21 |
| **WC (cm)** | 95.86 | 15.96 | 103.90 | 12.39 |
| **Weight (kg)** | 156.86 | 34.32 | 197.40 | 35.03 |
| **Height (cm)** | 161.65 | 6.46 | 175.79 | 6.87 |
| **BMI (kg/m2)** | 27.28 | 5.88 | 28.97 | 4.79 |
| **WHR (cm/cm)*** | 0.87 | 0.07 | 0.97 | 0.06 |
| **Hip (cm)*** | 105.03 | 12.25 | 105.19 | 8.77 |
| **WCHT (cm/cm)** | 0.59 | 0.10 | 0.59 | 0.07 |
| | **N** | **%** | **N** | **%** |
| - | - | - | - | - |
| - | - | - | - | - |
| - | - | - | - | - |
| **No high school degree** | 38 | 1.78 | 62 | 3.36 |
| **High School Degree** | 511 | 23.88 | 370 | 20.03 |
| - | - | - | - | - |
| **Some college** | 684 | 31.96 | 493 | 26.69 |
| **College grad** | 907 | 42.38 | 922 | 49.92 |
| - | - | - | - | - |
| - | - | - | - | - |
| - | - | - | - | - |
| **Current Smoker** | 179 | 8.71 | 175 | 9.86 |
| **Former Smoker** | 867 | 42.17 | 751 | 42.33 |
| **Never Smoker** | 1010 | 49.12 | 848 | 47.80 |

| | WHI - European American | | WHI - African American | | WHI - Hispanic/Latino | |
|---|---|---|---|---|---|---|
| | All Women | | All Women | | All Women | |
| | N = 990 | | N = 490 | | N = 276 | |
| | Mean | SD | Mean | SD | Mean | SD |
| **Age (years)** | 64.71 | 7.13 | 62.36 | 6.94 | 61.26 | 5.99 |
| **WC (cm)** | 88.22 | 13.51 | 92.72 | 12.44 | 89.70 | 12.53 |
| **Weight (kg)** | 75.08 | 15.50 | 83.13 | 16.04 | 74.46 | 14.43 |
| **Height (cm)** | 161.58 | 6.09 | 162.71 | 5.92 | 157.26 | 5.56 |
| **BMI (kg/m2)** | 28.78 | 5.75 | 31.46 | 5.93 | 30.07 | 5.35 |
| **WHR (cm/cm)** | 0.82 | 0.08 | 0.83 | 0.08 | 0.83 | 0.10 |
| **Hip (cm)** | 108.03 | 11.76 | 112.49 | 12.15 | 107.64 | 10.69 |
| **WCHT (cm/cm)** | 0.55 | 0.08 | 0.57 | 0.08 | 0.57 | 0.08 |
| | **N** | **%** | **N** | **%** | **N** | **%** |
| **No School** | 0 | 0.00 | 0 | 0.00 | 4 | 1.45 |
| **Grade school (1-4 yrs)** | 0 | 0.00 | 0 | 0.00 | 13 | 4.71 |
| **Grade school (5-8 yrs)** | 15 | 1.52 | 10 | 2.04 | 30 | 10.87 |
| **Some highschool (9-11 yrs)** | 33 | 3.33 | 35 | 7.16 | 22 | 7.97 |
| **High school degree/GED** | 223 | 22.53 | 69 | 14.11 | 42 | 15.22 |
| **Vocational/training school** | 121 | 12.22 | 62 | 12.68 | 29 | 10.51 |
| **Some college or associate degree** | 269 | 27.17 | 147 | 30.06 | 64 | 23.19 |
| **College graduate** | 95 | 9.60 | 45 | 9.20 | 22 | 7.97 |
| **Some post-graduate professional** | 84 | 8.48 | 46 | 9.41 | 15 | 5.43 |
| **Master's degree** | 121 | 12.22 | 62 | 12.68 | 31 | 11.23 |
| **Doctoral degree (PhD, MD, JD, etc)** | 29 | 2.93 | 13 | 2.66 | 4 | 1.45 |
| **Current Smoker** | 68 | 6.87 | 51 | 10.43 | 19 | 6.88 |
| **Former Smoker** | 424 | 42.83 | 199 | 40.70 | 81 | 29.35 |
| **Never Smoker** | 498 | 50.30 | 239 | 48.88 | 176 | 63.77 |

**Generalization**

| | RAINE - European Australian | | | |
|---|---|---|---|---|
| | Girls | | Boys | |
| | N = 397 | | N = 423 | |
| | Mean | SD | Mean | SD |
| **Age (years)** | 17.04 | 0.26 | 17.01 | 0.24 |
| **WC (cm)** | 77.86 | 11.55 | 80.70 | 10.46 |
| **Weight (kg)** | 63.65 | 12.21 | 72.42 | 13.88 |
| **Height (cm)** | 166.08 | 6.51 | 178.29 | 7.23 |
| **BMI (kg/m2)** | 23.08 | 4.26 | 22.74 | 3.95 |
| **WHR (cm/cm)** | 0.79 | 0.07 | 0.83 | 0.06 |
| **Hip (cm)** | 98.54 | 8.58 | 96.59 | 8.21 |
| **WCHT (cm/cm)** | 0.47 | 0.07 | 0.45 | 0.06 |
| | **N** | **%** | **N** | **%** |
| - | - | - | - | - |
| - | - | - | - | - |
| - | - | - | - | - |
| - | - | - | - | - |
| - | - | - | - | - |
| - | - | - | - | - |
| - | - | - | - | - |
| - | - | - | - | - |
| - | - | - | - | - |
| - | - | - | - | - |
| - | - | - | - | - |
| **Current Smoker** | 121 | 30.48 | 119 | 28.13 |
| - | - | - | - | - |
| **Never Smoker** | 276 | 69.52 | 304 | 72.10 |

**Supplementary Table 2**. Association results for discover stage, replication, and generalization for all significant CpGs associated with central adiposity and body fat distribution from discovery stage

| site | CHR | POS(hg19) | Gene | Gene Group* | CpG Island | ARIC BETA | ARIC SE | ARIC P | ARIC N | Replication Zscore | Replication P | Replication N | Discovery+Replication Zscore | Discovery+Replication P | Discovery+Replication N | RAINE BETA | RAINE SE | RAINE P | RAINE N | Directionally Consistent with Discovery | Known Adiposity Association† |
|---|---|---|---|---|---|---|---|---|---|---|---|---|---|---|---|---|---|---|---|---|---|
| **WCadjBMI** | | | | | | | | | | | | | | | | | | | | | |
| cg19693031 | 1 | 145441552 | ***TXNIP*** | 3'UTR | OpenSea | -0.057 | 0.010 | 4.13E-09 | 2,622 | -6.812 | **9.63E-12** | 5,741 | -8.936 | **4.05E-19** | 8,363 | -0.071 | 0.025 | **4.74E-03** | 819 | Y | - |
| cg16778018 | 16 | 4736225 | ***MGRN1*** | Body | S_Shelf | 0.123 | 0.022 | 2.48E-08 | 2,620 | -0.115 | 9.08E-01 | 5,741 | 3.025 | 2.48E-03 | 8,361 | 0.035 | 0.036 | 3.39E-01 | 819 | Y | - |
| cg13917614 | 17 | 40125660 | *CNP* | Body | OpenSea | 0.069 | 0.012 | 6.08E-09 | 2,623 | -0.153 | 8.78E-01 | 5,740 | 3.130 | 1.75E-03 | 8,363 | -0.012 | 0.023 | 6.00E-01 | 819 | N | - |
| cg06500161 | 21 | 43656587 | *ABCG1* | Body | S_Shore | 0.096 | 0.016 | 1.42E-09 | 2,623 | 8.563 | **1.10E-17** | 5,741 | 10.484 | **1.03E-25** | 8,364 | 0.083 | 0.035 | 1.75E-02 | 819 | Y | BMI(25935004,27773939, 28002404, 29099282, 29762635, 28095459); WC (29099282, 25935004, 29762635) |
| **WHRadjBMI** | | | | | | | | | | | | | | | | | | | | | |
| cg27185793 | 1 | 1238797 | *ACAP3* | Body | N_Shelf | 0.212 | 0.038 | 1.73E-08 | 2,621 | 2.15 | 3.15E-02 | 3,272 | 5.361 | **8.26E-08** | 5,893 | 0.067 | 0.061 | 2.73E-01 | 819 | Y | - |
| cg19693031 | 1 | 145441552 | ***TXNIP*** | 3'UTR | OpenSea | -0.151 | 0.019 | **3.48E-15** | 2,621 | -6.977 | **3.02E-12** | 3,273 | -10.449 | **1.48E-25** | 5,894 | -0.181 | 0.044 | **4.56E-05** | 819 | Y | - |
| cg22069247 | 2 | 232393256 | *NMUR1* | Body | N_Shore | 0.221 | 0.034 | **6.35E-11** | 2,619 | 2.975 | 2.93E-03 | 3,271 | 6.575 | **4.87E-11** | 5,890 | -0.039 | 0.049 | 4.27E-01 | 819 | N | - |
| cg19149463 | 3 | 11651759 | *VGLL4* | Body | OpenSea | 0.239 | 0.043 | **2.15E-08** | 2,621 | 0.566 | 5.71E-01 | 3,272 | 4.156 | 3.23E-05 | 5,893 | -0.121 | 0.055 | 2.88E-02 | 819 | N | - |
| cg19213703 | 3 | 177554561 | Intergenic | | OpenSea | -0.168 | 0.030 | **2.94E-08** | 2,621 | -2.373 | 1.76E-02 | 3,272 | -5.467 | **4.59E-08** | 5,893 | 0.057 | 0.049 | 2.48E-01 | 819 | N | - |
| cg02504211 | 3 | 194815434 | *C3orf21* | Body | OpenSea | 0.225 | 0.039 | **6.28E-09** | 2,620 | 0.985 | 3.25E-01 | 3,272 | 4.607 | 4.08E-06 | 5,892 | 0.133 | 0.066 | 4.39E-02 | 819 | Y | - |
| cg04816311 | 7 | 1066650 | *C7orf50* | Body | N_Shore | 0.133 | 0.023 | **1.04E-08** | 2,622 | 3.293 | **9.93E-04** | 3,272 | 6.272 | **3.57E-10** | 5,894 | 0.036 | 0.041 | 3.74E-01 | 819 | Y | BMI (25935004, 28095459) |
| cg26610247 | 8 | 142297175 | Intergenic | | S_Shore | 0.197 | 0.031 | **2.90E-10** | 2,622 | 3.384 | **7.15E-04** | 3,272 | 6.726 | **1.75E-11** | 5,894 | -0.016 | 0.055 | 7.70E-01 | 819 | N | - |
| cg17953300 | 11 | 65418265 | *SIPA1* | 3'UTR | N_Shore | 0.241 | 0.042 | **6.93E-09** | 2,620 | 0.624 | 5.33E-01 | 3,273 | 4.327 | 1.51E-05 | 5,893 | 0.025 | 0.087 | 7.74E-01 | 819 | Y | - |
| cg00574958 | 11 | 68607622 | *CPT1A* | 5'UTR | N_Shore | -0.442 | 0.061 | **5.80E-13** | 2,621 | -7.857 | **3.94E-15** | 3,273 | -10.66 | **1.57E-26** | 5,894 | -0.306 | 0.124 | 1.35E-02 | 819 | Y | BMI (28002404, 25935004, 29099282, 27826092, 29762635, 26110892, 28095459, 28947923); central obesity (28947923); obesity (28947923); WC (29762635, 28947923, 29099282, 26110892) |
| cg22348356 | 13 | 114891224 | *RASA3* | Body | OpenSea | 0.280 | 0.047 | **2.23E-09** | 2,620 | 1.765 | 7.76E-02 | 3,273 | 5.302 | 1.14E-07 | 5,893 | -0.043 | 0.058 | 4.60E-01 | 819 | N | - |
| cg06192883 | 15 | 52554171 | *MYO5C* | Body | OpenSea | 0.131 | 0.023 | **6.72E-09** | 2,621 | 3.628 | **2.86E-04** | 3,273 | 6.57 | **5.04E-11** | 5,894 | -0.023 | 0.065 | 7.28E-01 | 819 | N | BMI (28002404, 25935004, 28095459); WC(25935004) |
| cg16778018 | 16 | 4736225 | ***MGRN1*** | Body | S_Shelf | 0.239 | 0.044 | **5.39E-08** | 2,619 | 2.568 | 1.02E-02 | 3,273 | 5.54 | **3.03E-08** | 5,892 | 0.032 | 0.065 | 6.26E-01 | 819 | Y | - |
| cg06897661 | 16 | 50322074 | *ADCY7* | 5'UTR;1stExon | OpenSea | 0.198 | 0.034 | **6.60E-09** | 2,620 | 3.97 | **7.19E-05** | 3,271 | 6.827 | **8.70E-12** | 5,891 | -0.164 | 0.057 | 3.97E-03 | 819 | N | - |
| cg23580000 | 16 | 50322156 | *ADCY7* | 1stExon | OpenSea | 0.182 | 0.029 | **2.17E-10** | 2,622 | 4.261 | **2.04E-05** | 3,273 | 7.409 | **1.28E-13** | 5,895 | -0.124 | 0.046 | 6.29E-03 | 819 | N | - |
| cg15863539 | 17 | 17716950 | *SREBF1* | Body | S_Shore | 0.499 | 0.078 | **1.68E-10** | 2,619 | 3.391 | **6.97E-04** | 3,273 | 6.786 | **1.15E-11** | 5,892 | 0.240 | 0.113 | 3.46E-02 | 819 | Y | - |
| cg20544516 | 17 | 17717183 | *MIR33B/SREBF1* | Body | S_Shore | 0.320 | 0.052 | **5.42E-10** | 2,621 | 3.382 | **7.21E-04** | 3,273 | 6.659 | **2.77E-11** | 5,894 | 0.160 | 0.055 | 3.70E-03 | 819 | Y | - |
| cg13917614 | 17 | 40125660 | *CNP* | Body | OpenSea | 0.142 | 0.024 | **1.76E-09** | 2,622 | -0.064 | 9.49E-01 | 3,272 | 3.967 | 7.28E-05 | 5,894 | -0.083 | 0.041 | 4.46E-02 | 819 | N | - |
| cg08352115 | 17 | 66356057 | *ARSG* | Body | OpenSea | 0.189 | 0.033 | **1.49E-08** | 2,622 | 2.531 | 1.14E-02 | 3,273 | 5.662 | **1.49E-08** | 5,895 | 0.030 | 0.049 | 5.45E-01 | 819 | Y | - |
| cg00994936 | 19 | 1423902 | *DAZAP1* | Body | Island | 0.279 | 0.047 | **3.11E-09** | 2,621 | 2.861 | 4.23E-03 | 3,272 | 6.083 | **1.18E-09** | 5,893 | 0.197 | 0.058 | **6.06E-04** | 819 | Y | - |
| cg06500161 | 21 | 43656587 | *ABCG1* | Body | S_Shore | 0.228 | 0.032 | **5.64E-13** | 2,622 | 8.274 | **1.29E-16** | 3,273 | 10.973 | **5.15E-28** | 5,895 | 0.135 | 0.062 | 2.92E-02 | 819 | Y | BMI(25935004,27773939, 28002404, 29099282, 29762635, 28095459); WC (29099282, 25935004, 29762635) |
| **WCHTadjBMI** | | | | | | | | | | | | | | | | | | | | | |
| cg19693031 | 1 | 145441552 | ***TXNIP*** | 3'UTR | OpenSea | -0.08408221 | 0.01248584 | **1.65E-11** | 2,622 | -7.585 | **3.31E-14** | 5,734 | -10.056 | **8.65E-24** | 8,356 | -0.022 | 0.024 | 3.57E-01 | 819 | Y | - |
| cg22069247 | 2 | 232393256 | *NMUR1* | Body | N_Shore | 0.1176164 | 0.02206056 | **9.74E-08** | 2,620 | 1.707 | 8.78E-02 | 5,732 | 4.4 | 1.08E-05 | 8,352 | 0.034 | 0.026 | 1.87E-01 | 819 | Y | - |
| cg26610247 | 8 | 142297175 | Intergenic | | S_Shore | 0.1087791 | 0.02026264 | **7.94E-08** | 2,623 | 2.482 | 1.31E-02 | 5,734 | 5.064 | 4.11E-07 | 8,357 | 0.062 | 0.029 | 3.07E-02 | 819 | Y | - |
| cg00574958 | 11 | 68607622 | *CPT1A* | 5'UTR | N_Shore | -0.2666759 | 0.03988153 | **2.28E-11** | 2,622 | -5.804 | **6.47E-09** | 5,734 | -8.554 | **1.19E-17** | 8,356 | -0.088 | 0.066 | 1.78E-01 | 819 | Y | BMI (28002404, 25935004, 29099282, 27826092, 29762635, 26110892, 28095459, 28947923); central obesity (28947923); obesity (28947923); WC (29762635, 28947923, 29099282, 26110892) |
| cg09940677 | 14 | 103415458 | *CDC42BPB* | Body | OpenSea | 0.1179211 | 0.02128195 | **3.01E-08** | 2,622 | 1.28 | 2.01E-01 | 5,734 | 4.164 | 3.13E-05 | 8,356 | -0.02 | 0.045 | 6.49E-01 | 819 | N | - |
| cg16778018 | 16 | 4736225 | ***MGRN1*** | Body | S_Shelf | 0.1588406 | 0.02851028 | **2.53E-08** | 2,620 | 1.502 | 1.33E-01 | 5,734 | 4.365 | 1.27E-05 | 8,354 | 0.024 | 0.034 | 4.90E-01 | 819 | Y | - |
| cg12537003 | 17 | 2886453 | *RAP1GAP2* | Body | OpenSea | 0.1782609 | 0.03224328 | **3.23E-08** | 2,623 | 3.482 | **4.98E-04** | 5,734 | 5.981 | **2.21E-09** | 8,357 | 0.002 | 0.031 | 9.46E-01 | 819 | Y | - |

*Note that multiple GeneRef Groups denote location repsective to splice sites. New annotation only provided when ref location or ref gene changes.

†For any CpG with a known association with an obesity-related trait, we list the trait and PubMed PMID.

Gene names are bolded if they are significantly associated with all three traits.

P-values are bolded if they meet significance criteria.

**Supplementary Table 3**. Ancestry-specific meta-analysis results for all significant CpGs from the discovery analysis. Bold indicates statistical sigificance following multiple -test correction (P<1E-7 for African Ancestry analysis with ARIC, and Bonferroni-corrected by trait for other analyses). Z-scores that are not directionally consistent with discovery results are italicized.

| | | | | | | African Ancestry (ARIC + WHI) | | | | African Ancestry (WHI only) | | | European Descent | | | | Hispanic/Latino | | | | |
|---|---|---|---|---|---|---|---|---|---|---|---|---|---|---|---|---|---|---|---|---|---|
| site | CHR | POS(hg19) | Gene | Gene Group* | CpG Island | Zscore | P | $P_{het}$ | N | Zscore | P | N | Zscore | P | $P_{het}$ | N | Zscore | P | N | Significant in Replication | Directionally Consistent |
| **WCadjBMI** | | | | | | | | | | | | | | | | | | | | | |
| cg19693031 | 1 | 145441552 | ***TXNIP*** | 3'UTR | OpenSea | -6.101 | **1.06E-09** | 0.4824 | 3,112 | -1.776 | 7.57E-02 | 490 | -6.472 | **9.67E-11** | 0.0591 | 4,976 | -1.223 | 2.21E-01 | 275 | Yes | Yes |
| cg16778018 | 16 | 4736225 | ***MGRN1*** | Body | S_Shelf | 5.703 | **1.18E-08** | 0.3916 | 3,110 | 1.478 | 1.40E-01 | 490 | *-1.109* | 2.67E-01 | 0.0397 | 4,976 | 2.220 | 2.64E-02 | 275 | No | *No* |
| cg13917614 | 17 | 40125660 | *CNP* | Body | OpenSea | 4.729 | 2.26E-06 | 0.0002 | 3,113 | *-1.535* | 1.25E-01 | 490 | 0.196 | 8.45E-01 | 0.1313 | 4,975 | 0.515 | 6.07E-01 | 275 | No | *No* |
| cg06500161 | 21 | 43656587 | *ABCG1* | Body | S_Shore | 6.775 | **1.25E-11** | 0.6762 | 3,113 | 3.071 | **2.13E-03** | 490 | 7.448 | **9.46E-14** | 0.3036 | 4,976 | 3.341 | **8.34E-04** | 275 | Yes | Yes |
| **WHRadjBMI** | | | | | | | | | | | | | | | | | | | | | |
| cg27185793 | 1 | 1238797 | *ACAP3* | Body | N_Shelf | 5.103 | 3.34E-07 | 0.0164 | 3,110 | *-0.180* | 8.57E-01 | 489 | 1.904 | 5.69E-02 | 0.7466 | 2,507 | 1.905 | 5.68E-02 | 276 | No | No |
| cg19693031 | 1 | 145441552 | ***TXNIP*** | 3'UTR | OpenSea | -7.840 | **4.51E-15** | 0.0886 | 3,110 | -1.545 | 1.22E-01 | 489 | -6.912 | **4.78E-12** | 0.2163 | 2,508 | -1.133 | 2.57E-01 | 276 | Yes | Yes |
| cg22069247 | 2 | 232393256 | *NMUR1* | Body | N_Shore | 6.506 | **7.70E-11** | 0.1561 | 3,108 | 1.279 | 2.01E-01 | 489 | 2.756 | 5.85E-03 | 0.6696 | 2,507 | 0.234 | 8.15E-01 | 275 | No | Yes |
| cg19149463 | 3 | 11651759 | *VGLL4* | Body | OpenSea | 4.698 | 2.63E-06 | 0.0012 | 3,110 | *-1.116* | 2.64E-01 | 489 | 0.716 | 4.74E-01 | 0.9575 | 2,507 | 1.276 | 2.02E-01 | 276 | No | *No* |
| cg19213703 | 3 | 177554561 | Intergenic | | OpenSea | -4.789 | 1.68E-06 | 0.0038 | 3,110 | *0.762* | 4.46E-01 | 489 | -2.809 | 4.98E-03 | 0.4531 | 2,507 | -0.720 | 4.71E-01 | 276 | No | *No* |
| cg02504211 | 3 | 194815434 | *C3orf21* | Body | OpenSea | 5.931 | **3.02E-09** | 0.3577 | 3,109 | 1.508 | 1.32E-01 | 489 | 0.399 | 6.90E-01 | 0.5394 | 2,507 | 0.182 | 8.55E-01 | 276 | No | Yes |
| cg04816311 | 7 | 1066650 | *C7orf50* | Body | N_Shore | 6.457 | **1.07E-10** | 0.6058 | 3,110 | 3.032 | 2.43E-03 | 488 | 1.982 | 4.75E-02 | 0.0871 | 2,508 | 1.330 | 1.83E-01 | 276 | Yes | Yes |
| cg26610247 | 8 | 142297175 | Intergenic | | S_Shore | 6.223 | **4.88E-10** | 0.1361 | 3,111 | 1.099 | 2.72E-01 | 489 | 2.899 | 3.74E-03 | 0.7828 | 2,507 | 1.450 | 1.47E-01 | 276 | Yes | Yes |
| cg17953300 | 11 | 65418265 | *SIPA1* | 3'UTR | N_Shore | 5.507 | **3.65E-08** | 0.0631 | 3,109 | 0.478 | 6.33E-01 | 489 | 0.250 | 8.03E-01 | 0.9046 | 2,508 | 0.758 | 4.48E-01 | 276 | No | Yes |
| cg00574958 | 11 | 68607622 | *CPT1A* | 5'UTR | N_Shore | -7.908 | **2.61E-15** | 0.8894 | 3,110 | -3.264 | **1.10E-03** | 489 | -7.653 | **1.97E-14** | 0.1047 | 2,508 | *0.357* | 7.21E-01 | 276 | Yes | *No* |
| cg22348356 | 13 | 114891224 | *RASA3* | Body | OpenSea | 5.618 | **1.93E-08** | 0.0381 | 3,109 | 0.324 | 7.46E-01 | 489 | 1.648 | 9.93E-02 | 0.7445 | 2,508 | 0.678 | 4.98E-01 | 276 | No | Yes |
| cg06192883 | 15 | 52554171 | *MYO5C* | Body | OpenSea | 6.169 | **6.89E-10** | 0.7339 | 3,110 | 2.134 | 3.29E-02 | 489 | 2.389 | 1.69E-02 | 0.4681 | 2,508 | 2.451 | 1.42E-02 | 276 | Yes | Yes |
| cg16778018 | 16 | 4736225 | ***MGRN1*** | Body | S_Shelf | 5.927 | **3.08E-09** | 0.9933 | 3,108 | 2.359 | 1.83E-02 | 489 | 1.825 | 6.80E-02 | 0.4411 | 2,508 | 0.203 | 8.39E-01 | 276 | No | Yes |
| cg06897661 | 16 | 50322074 | *ADCY7* | 5'UTR;1stExon | OpenSea | 5.879 | **4.13E-09** | 0.3093 | 3,108 | 1.396 | 1.63E-01 | 488 | 3.614 | **3.02E-04** | 0.7465 | 2,507 | 0.919 | 3.58E-01 | 276 | Yes | Yes |
| cg23580000 | 16 | 50322156 | *ADCY7* | 1stExon | OpenSea | 6.676 | **2.46E-11** | 0.5793 | 3,111 | 2.138 | 3.25E-02 | 489 | 3.852 | **1.17E-04** | 0.3300 | 2,508 | 0.214 | 8.31E-01 | 276 | Yes | Yes |
| cg15863539 | 17 | 17716950 | *SREBF1* | Body | S_Shore | 6.601 | **4.08E-11** | 0.4079 | 3,108 | 1.859 | 6.31E-02 | 489 | 2.770 | 5.60E-03 | 0.5171 | 2,508 | 0.852 | 3.94E-01 | 276 | Yes | Yes |
| cg20544516 | 17 | 17717183 | *MIR33B/SREBF1* | Body | S_Shore | 6.517 | **7.18E-11** | 0.5728 | 3,110 | 2.066 | 3.88E-02 | 489 | 3.045 | **2.32E-03** | 0.4373 | 2,508 | *-0.286* | 7.75E-01 | 276 | Yes | *No* |
| cg13917614 | 17 | 40125660 | *CNP* | Body | OpenSea | 5.094 | 3.51E-07 | 0.0007 | 3,111 | *-1.089* | 2.76E-01 | 489 | 0.200 | 8.42E-01 | 0.7903 | 2,507 | 0.628 | 5.30E-01 | 276 | No | *No* |
| cg08352115 | 17 | 66356057 | *ARSG* | Body | OpenSea | 5.632 | **1.78E-08** | 0.2142 | 3,111 | 1.093 | 2.75E-01 | 489 | 2.390 | 1.69E-02 | 0.8737 | 2,508 | 0.056 | 9.55E-01 | 276 | No | Yes |
| cg00994936 | 19 | 1423902 | *DAZAP1* | Body | Island | 6.567 | **5.15E-11** | 0.7939 | 3,109 | 2.841 | 4.49E-03 | 488 | 1.651 | 9.86E-02 | 0.9899 | 2,508 | 1.093 | 2.74E-01 | 276 | No | Yes |
| cg06500161 | 21 | 43656587 | *ABCG1* | Body | S_Shore | 7.884 | **3.16E-15** | 0.9408 | 3,111 | 3.194 | **1.40E-03** | 489 | 7.134 | **9.76E-13** | 0.1979 | 2,508 | 2.737 | 6.20E-03 | 276 | Yes | Yes |
| **WCHTadjBMI** | | | | | | | | | | | | | | | | | | | | | |
| cg19693031 | 1 | 145441552 | ***TXNIP*** | 3'UTR | OpenSea | -6.927 | **4.31E-12** | 0.3440 | 3,111 | -1.877 | 6.05E-02 | 489 | -7.031 | **2.05E-12** | 0.1314 | 4,970 | -2.243 | 2.49E-02 | 275 | Yes | Yes |
| cg22069247 | 2 | 232393256 | *NMUR1* | Body | N_Shore | 4.990 | 6.05E-07 | 0.0582 | 3,109 | 0.240 | 8.10E-01 | 489 | 1.340 | 1.80E-01 | 0.0098 | 4,969 | 1.780 | 7.50E-02 | 274 | No | Yes |
| cg26610247 | 8 | 142297175 | Intergenic | | S_Shore | 5.318 | 1.05E-07 | 0.2199 | 3,112 | 0.982 | 3.26E-01 | 489 | 1.689 | 9.12E-02 | 0.7573 | 4,970 | 2.846 | **4.43E-03** | 275 | No | Yes |
| cg00574958 | 11 | 68607622 | *CPT1A* | 5'UTR | N_Shore | -6.874 | **6.26E-12** | 0.3424 | 3,111 | -1.854 | 6.38E-02 | 489 | -5.763 | **8.26E-09** | 0.7363 | 4,970 | *0.468* | 6.40E-01 | 275 | Yes | *No* |
| cg09940677 | 14 | 103415458 | *CDC42BPB* | Body | OpenSea | 5.137 | 2.80E-07 | 0.0374 | 3,111 | 0.126 | 9.00E-01 | 489 | 1.010 | 3.13E-01 | 0.1355 | 4,970 | 1.382 | 1.67E-01 | 275 | No | Yes |
| cg16778018 | 16 | 4736225 | ***MGRN1*** | Body | S_Shelf | 5.584 | **2.35E-08** | 0.2617 | 3,109 | 1.184 | 2.36E-01 | 489 | 0.714 | 4.75E-01 | 0.3131 | 4,970 | 2.243 | 2.49E-02 | 275 | No | Yes |
| cg12537003 | 17 | 2886453 | *RAP1GAP2* | Body | OpenSea | 5.427 | **5.72E-08** | 0.1685 | 3,112 | 0.887 | 3.75E-01 | 489 | 3.095 | **1.97E-03** | 0.1540 | 4,970 | 1.560 | 1.19E-01 | 275 | Yes | Yes |

**Supplementary Table 4**. NHGRI-EBI GWAS Catalog lookups for genetic associations within 100 kb (+/-) of significantly associated CpGs

| Current Trait | Site | CHR | POS (hg19) | Distance | Disease/Trait in Catalog | Mapped Gene | SNP(s) | EAF | P | PMID |
|---|---|---|---|---|---|---|---|---|---|---|
| WCHTadjBMI | cg12537003 | 17 | 2886453 | 4448 | White blood cell count | RAP1GAP2 | rs62092000 | NR | 1.00E-09 | 30595370 |
| WCHTadjBMI | cg12537003 | 17 | 2886453 | 20664 | Eosinophil counts | RAP1GAP2, AC015921.1 | rs34224666 | NR | 5.00E-14 | 30595370 |
| WCHTadjBMI | cg12537003 | 17 | 2886453 | 23301 | Male-pattern baldness | RAP1GAP2 | rs9901627 | 0.335373 | 4.00E-09 | 30573740 |
| WCHTadjBMI | cg12537003 | 17 | 2886453 | 2865 | Lymphocyte counts | RAP1GAP2 | rs17762452 | 0.3188 | 8.00E-14 | 27863252 |
| WHRadjBMI | cg04816311 | 7 | 1066650 | 39362 | Mean corpuscular hemoglobin | C7orf50 | rs1104888 | NR | 4.00E-12 | 30595370 |
| **WHRadjBMI** | **cg04816311** | **7** | **1066650** | **17127** | **Cholesterol, total** | **C7orf50** | **rs1997243** | **0.16** | **3.00E-10** | **24097068** |
| **WHRadjBMI** | **cg04816311** | **7** | **1066650** | **1256** | **Total cholesterol levels** | **C7orf50** | **rs2362529** | **0.7754** | **8.00E-19** | **30275531** |
| **WHRadjBMI** | **cg04816311** | **7** | **1066650** | **16268** | **LDL cholesterol** | **AC073957.1, C7orf50** | **rs10275712** | **0.198** | **2.00E-16** | **30275531** |
| **WHRadjBMI** | **cg04816311** | **7** | **1066650** | **8457** | **C-reactive protein levels or total cholesterol levels (pleiotropy)** | **C7orf50** | **rs6951245** | **NR** | **3.00E-09** | **27286809** |
| WHRadjBMI | cg26610247 | 8 | 142297175 | 1004 | Heel bone mineral density | AC011676.3, SLC45A4 | rs7000279 | NR | 3.00E-10 | 30595370 |
| WHRadjBMI | cg26610247 | 8 | 142297175 | 40615 | Sum neutrophil eosinophil counts | AC011676.4 - LINC01300 | rs6986779 | 0.3592 | 5.00E-15 | 27863252 |
| WHRadjBMI | cg26610247 | 8 | 142297175 | 68266 | High light scatter reticulocyte count | SLC45A4 | rs753778 | 0.2935 | 5.00E-10 | 27863252 |
| **WHRadjBMI** | **cg26610247** | **8** | **142297175** | **99305** | **Systolic blood pressure** | **GPR20 - AC100803.1** | **rs76735299** | **NR** | **4.00E-11** | **30595370** |
| WHRadjBMI | cg26610247 | 8 | 142297175 | 68266 | Reticulocyte fraction of red cells | SLC45A4 | rs753778 | 0.2935 | 3.00E-09 | 27863252 |
| **WHRadjBMI** | **cg26610247** | **8** | **142297175** | **69911** | **Systolic blood pressure** | **GPR20** | **rs34591516** | **0.0587** | **1.00E-09** | **27618447** |
| **WHRadjBMI** | **cg26610247** | **8** | **142297175** | **99305** | **Systolic blood pressure** | **GPR20 - AC100803.1** | **rs76735299** | | **4.00E-08** | **27841878** |
| WHRadjBMI | cg26610247 | 8 | 142297175 | 68266 | High light scatter reticulocyte percentage of red cells | SLC45A4 | rs753778 | 0.2935 | 1.00E-10 | 27863252 |
| WHRadjBMI | cg26610247 | 8 | 142297175 | 40615 | Neutrophil count | AC011676.4 - LINC01300 | rs6986779 | 0.3592 | 1.00E-14 | 27863252 |
| **WHRadjBMI** | **cg26610247** | **8** | **142297175** | **77897** | **Diastolic blood pressure** | **GPR20** | **rs78192203** | **NR** | **4.00E-11** | **28498854** |
| WHRadjBMI | cg26610247 | 8 | 142297175 | 64919 | Immature fraction of reticulocytes | SLC45A4 | rs59918340 | 0.477 | 8.00E-10 | 27863252 |
| WHRadjBMI | cg26610247 | 8 | 142297175 | 1004 | Heel bone mineral density | AC011676.3, SLC45A4 | rs7000279 | NR | 8.00E-12 | 30048462 |
| WHRadjBMI | cg26610247 | 8 | 142297175 | 96708 | White blood cell count | DENND3 | rs1045303 | NR | 5.00E-10 | 30595370 |
| WHRadjBMI | cg26610247 | 8 | 142297175 | 40615 | Granulocyte count | AC011676.4 - LINC01300 | rs6986779 | 0.3592 | 6.00E-15 | 27863252 |
| WHRadjBMI | cg26610247 | 8 | 142297175 | 66988 | Lung function (FEV1/FVC) | AC100803.2 | rs60772015 | NR | 3.00E-08 | 30595370 |
| WHRadjBMI | cg26610247 | 8 | 142297175 | 31544 | White blood cell count | SLC45A4 - AC011676.4 | rs13261848 | 0.3614 | 7.00E-15 | 27863252 |
| WHRadjBMI | cg26610247 | 8 | 142297175 | 40615 | Myeloid white cell count | AC011676.4 - LINC01300 | rs6986779 | 0.3592 | 1.00E-14 | 27863252 |
| **WHRadjBMI** | **cg26610247** | **8** | **142297175** | **77897** | **Blood pressure traits (multi-trait analysis)** | **GPR20** | **rs78192203** | **NR** | **2.00E-08** | **28498854** |
| WHRadjBMI | cg26610247 | 8 | 142297175 | 40615 | Sum basophil neutrophil counts | AC011676.4 - LINC01300 | rs6986779 | 0.3592 | 1.00E-14 | 27863252 |
| WHRadjBMI | cg26610247 | 8 | 142297175 | 32285 | White blood cell count | SLC45A4 - AC011676.4 | rs34634548 | NR | 0.00E+00 | 30595370 |
| WHRadjBMI | cg26610247 | 8 | 142297175 | 1004 | Heel bone mineral density | AC011676.3, SLC45A4 | rs7000279 | 0.736828 | 4.00E-10 | 30598549 |
| WHRadjBMI | cg26610247 | 8 | 142297175 | 4849 | Plateletcrit | AC011676.3, SLC45A4 | rs12677618 | 0.2611 | 3.00E-11 | 27863252 |
| WHRadjBMI | cg26610247 | 8 | 142297175 | 17763 | Mean corpuscular hemoglobin | SLC45A4 | rs6992286 | NR | 5.00E-08 | 30595370 |
| **WHRadjBMI** | **cg26610247** | **8** | **142297175** | **49196** | **Birth weight** | **SLC45A4** | **rs12543725** | **0.6** | **2.00E-09** | **27680694** |
| **WHRadjBMI** | **cg00574958** | **11** | **68607622** | **45294** | **Lipid metabolism phenotypes** | **CPT1A** | **rs17610395** | **NR** | **8.00E-12** | **22286219** |
| WHRadjBMI | cg00574958 | 11 | 68607622 | 74166 | Heel bone mineral density | IGHMBP2 | rs557266652 | 0.998646 | 3.00E-08 | 28869591 |
| WHRadjBMI | cg00574958 | 11 | 68607622 | 61090 | Heel bone mineral density | CPT1A | rs186065533 | NR | 1.00E-15 | 30595370 |
| WHRadjBMI | cg00574958 | 11 | 68607622 | 83569 | Total body bone mineral density | CPT1A | rs2278907 | 0.5455 | 3.00E-08 | 29304378 |
| WHRadjBMI | cg00574958 | 11 | 68607622 | 74166 | Heel bone mineral density | IGHMBP2 | rs557266652 | NR | 3.00E-20 | 30048462 |
| WHRadjBMI | cg00574958 | 11 | 68607622 | 85569 | Total body bone mineral density | TESMIN - CPT1A | rs3018712 | 0.1787 | 4.00E-11 | 29304378 |
| **WHRadjBMI** | **cg00574958** | **11** | **68607622** | **9568** | **Lipid traits (pleiotropy) (HIPO component 1)** | **CPT1A** | **rs7938117** | **NR** | **3.00E-11** | **30289880** |
| WHRadjBMI | cg00574958 | 11 | 68607622 | 74166 | Heel bone mineral density | IGHMBP2 | rs557266652 | 0.998715 | 5.00E-11 | 28869591 |
| **WHRadjBMI** | **cg00574958** | **11** | **68607622** | **92153** | **Body mass index** | **IGHMBP2** | **rs653264** | **NR** | **5.00E-10** | **30595370** |
| WHRadjBMI | cg22348356 | 13 | 114891224 | 70259 | Eosinophil counts | CFAP97D2, AL160396.1 | rs9590475 | NR | 5.00E-14 | 30595370 |
| WHRadjBMI | cg22348356 | 13 | 114891224 | 73963 | Hair color | RASA3 | rs6560957 | NR | 3.00E-09 | 30595370 |
| WHRadjBMI | cg22348356 | 13 | 114891224 | 56286 | Lymphocyte counts | CFAP97D2, AL160396.1 | rs547320937 | 0.0064 | 3.00E-09 | 27863252 |
| WHRadjBMI | cg06897661 | 16 | 50322074 | 13000 | Autoimmune traits | ADCY7 | rs78534766 | NR | 5.00E-13 | 30595370 |
| WHRadjBMI | cg06897661 | 16 | 50322074 | 13000 | Inflammatory bowel disease | ADCY7 | rs78534766 | | 2.00E-12 | 28067908 |
| WHRadjBMI | cg06897661 | 16 | 50322074 | 13000 | Hypothyroidism | ADCY7 | rs78534766 | NR | 1.00E-16 | 30595370 |

| | | | | | | | | | | |
|---|---|---|---|---|---|---|---|---|---|---|
| WHRadjBMI | cg06897661 | 16 | 50322074 | 72994 | Adolescent idiopathic scoliosis | TENT4B | rs3814284 | NR | 2.00E-10 | 30019117 |
| WHRadjBMI | cg06897661 | 16 | 50322074 | 13000 | Ulcerative colitis | ADCY7 | rs78534766 | | 3.00E-13 | 28067908 |
| WHRadjBMI | cg06897661 | 16 | 50322074 | 17825 | Pediatric autoimmune diseases | ADCY7 | rs77150043 | 0.23 | 6.00E-09 | 26301688 |
| WHRadjBMI | cg15863539 | 17 | 17716950 | 85156 | Eczema | RAI1 | rs62064086 | NR | 2.00E-10 | 30595370 |
| WHRadjBMI | cg15863539 | 17 | 17716950 | 5680 | Schizophrenia | RAI1 | rs4925114 | NR | 3.00E-08 | 29483656 |
| WHRadjBMI | cg15863539 | 17 | 17716950 | 33957 | TB-LM or TBLH-BMD (pleiotropy) | TOM1L2 | rs7501812 | 0.41 | 1.00E-10 | 28743860 |
| WHRadjBMI | cg15863539 | 17 | 17716950 | 9845 | Lymphocyte counts | RAI1 | rs3818717 | 0.5924 | 1.00E-12 | 27863252 |
| WHRadjBMI | cg15863539 | 17 | 17716950 | 87775 | Total body bone mineral density | TOM1L2 | rs8070128 | 0.5763 | 2.00E-11 | 29304378 |
| WHRadjBMI | cg15863539 | 17 | 17716950 | 55148 | Educational attainment (years of education) | RAI1 | rs4925109 | 0.3172 | 2.00E-10 | 30038396 |
| WHRadjBMI | cg15863539 | 17 | 17716950 | 59439 | Lung function (FVC) | TOM1L2 | rs8080061 | NR | 1.00E-08 | 30595370 |
| **WHRadjBMI** | **cg15863539** | **17** | **17716950** | **63539** | **Type 2 diabetes** | **RAI1** | **rs12945601** | **0.386396556** | **2.00E-09** | **30054458** |
| WHRadjBMI | cg15863539 | 17 | 17716950 | 9845 | White blood cell count | RAI1 | rs3818717 | NR | 4.00E-23 | 30595370 |
| WHRadjBMI | cg15863539 | 17 | 17716950 | 5680 | General cognitive ability | RAI1 | rs4925114 | NR | 2.00E-08 | 29844566 |
| WHRadjBMI | cg15863539 | 17 | 17716950 | 78189 | Eosinophil counts | RAI1 | rs7219213 | NR | 9.00E-12 | 30595370 |
| WHRadjBMI | cg15863539 | 17 | 17716950 | 419 | Resting heart rate | SREBF1 | rs12941356 | 0.42 | 5.00E-08 | 27798624 |
| **WHRadjBMI** | **cg15863539** | **17** | **17716950** | **5680** | **Body mass index** | **RAI1** | **rs4925114** | **0.6** | **3.00E-08** | **28892062** |

**Supplementary Table 5**. Lookup of known CpGs associated with obesity-related traits. P-values that remain significant following Bonferroni correction (P<0.05/408 = 1.23E-4) are bolded and shaded with grey. CpGs that were significantly associated with WCadjBMI, WHRadjBMI, or WCHTadjBMI in the current paper are shaded in yellow.

| | | | | | | WCadjBMI | | | | WHRadjBMI | | | | WCHTadjBMI | | | |
|---|---|---|---|---|---|---|---|---|---|---|---|---|---|---|---|---|---|
| Trait | Site | CHR | POS(hg19) | Gene | Gene | BETA | SE | P | N | BETA | SE | P | N | BETA | SE | P | N |
| Childhood/Young Adult Obesity | cg21815220 | 1 | 1140273 | *TNFRSF4* | TSS1500 | -0.0599 | 0.0210 | 4.40E-03 | 2,623 | -0.1257 | 0.0420 | 2.77E-03 | 2,622 | -0.0834 | 0.0272 | 2.22E-03 | 2,623 |
| BMI | cg09315878 | 1 | 1142443 | *SDF4* | 3UTR | -0.0091 | 0.0121 | 4.54E-01 | 2,623 | -0.0225 | 0.0241 | 3.51E-01 | 2,622 | -0.0009 | 0.0157 | 9.52E-01 | 2,623 |
| Childhood/Young Adult Obesity | cg14531564 | 1 | 1144716 | *SDF4* | Body | -0.0245 | 0.0177 | 1.66E-01 | 2,622 | -0.0605 | 0.0354 | 8.76E-02 | 2,621 | -0.0354 | 0.0230 | 1.24E-01 | 2,622 |
| Childhood/Young Adult Obesity | cg01979157 | 1 | 2150873 | *SKI* | 1stExon | -0.0823 | 0.0321 | 1.03E-02 | 2,621 | -0.1452 | 0.0641 | 2.35E-02 | 2,620 | -0.1580 | 0.0415 | 1.38E-04 | 2,621 |
| Childhood/Young Adult Obesity | cg05603985 | 1 | 2150909 | *SKI* | 1stExon | -0.0428 | 0.0168 | 1.09E-02 | 2,618 | -0.0937 | 0.0335 | 5.24E-03 | 2,617 | -0.0843 | 0.0217 | **1.05E-04** | 2,618 |
| Childhood/Young Adult Obesity | cg17942763 | 1 | 2153332 | *SKI* | Body | -0.0163 | 0.0148 | 2.71E-01 | 2,621 | -0.0336 | 0.0295 | 2.55E-01 | 2,620 | -0.0232 | 0.0191 | 2.25E-01 | 2,621 |
| BMI | cg08648047 | 1 | 10951148 | *C1orf127* | Body | 0.0276 | 0.0233 | 2.38E-01 | 2,623 | 0.0945 | 0.0466 | 4.26E-02 | 2,622 | 0.0515 | 0.0302 | 8.85E-02 | 2,623 |
| BMI | cg03885055 | 1 | 16595819 | *C1orf144* | 3UTR | -0.0088 | 0.0186 | 6.37E-01 | 2,623 | -0.0194 | 0.0372 | 6.02E-01 | 2,622 | -0.0241 | 0.0241 | 3.17E-01 | 2,623 |
| BMI | cg08540100 | 1 | 23761573 | | | 0.0449 | 0.0129 | 5.00E-04 | 2,623 | 0.0869 | 0.0258 | 7.50E-04 | 2,622 | 0.0548 | 0.0167 | 1.05E-03 | 2,623 |
| BMI | cg11673687 | 1 | 27299142 | *SLC9A1* | 3UTR | 0.0466 | 0.0313 | 1.37E-01 | 2,621 | 0.0940 | 0.0625 | 1.33E-01 | 2,620 | 0.0342 | 0.0405 | 3.98E-01 | 2,621 |
| BMI | cg26161057 | 1 | 27754187 | *AHDC1* | 5UTR | -0.0062 | 0.0341 | 8.56E-01 | 2,623 | -0.0601 | 0.0681 | 3.77E-01 | 2,622 | -0.0415 | 0.0442 | 3.48E-01 | 2,623 |
| BMI | cg12484113 | 1 | 27771344 | *AHDC1* | 5UTR | 0.0370 | 0.0175 | 3.46E-02 | 2,617 | 0.0867 | 0.0350 | 1.32E-02 | 2,616 | 0.0263 | 0.0227 | 2.46E-01 | 2,617 |
| BMI, WC | cg17822325 | 1 | 31669049 | *SERINC2* | Body | 0.0262 | 0.0100 | 9.09E-03 | 2,621 | 0.0697 | 0.0200 | 5.01E-04 | 2,620 | 0.0391 | 0.0130 | 2.64E-03 | 2,621 |
| BMI | cg16815882 | 1 | 35681196 | *KIAA0319L* | Body | 0.0153 | 0.0151 | 3.11E-01 | 2,623 | 0.0481 | 0.0301 | 1.10E-01 | 2,622 | 0.0370 | 0.0195 | 5.80E-02 | 2,623 |
| BMI WC | cg17901584 | 1 | 55126294 | *DHCR24* | TSS1500 | -0.0228 | 0.0103 | 2.76E-02 | 2,623 | -0.0595 | 0.0206 | 3.93E-03 | 2,622 | -0.0276 | 0.0134 | 3.91E-02 | 2,623 |
| BMI | cg10092518 | 1 | 57442642 | *DAB1* | Body | -0.0342 | 0.0195 | 7.87E-02 | 2,622 | -0.1139 | 0.0388 | 3.31E-03 | 2,621 | -0.0556 | 0.0252 | 2.74E-02 | 2,622 |
| Childhood/Young Adult Obesity | cg20849150 | 1 | 59295581 | | | -0.0200 | 0.0176 | 2.55E-01 | 2,620 | -0.0518 | 0.0351 | 1.40E-01 | 2,619 | -0.0332 | 0.0228 | 1.45E-01 | 2,620 |
| BMI | cg25001190 | 1 | 61441423 | *NFIA* | Body | -0.0360 | 0.0124 | 3.66E-03 | 2,623 | -0.0929 | 0.0247 | 1.65E-04 | 2,622 | -0.0507 | 0.0160 | 1.54E-03 | 2,623 |
| BMI | cg06872964 | 1 | 78857838 | *IFI44L* | TSS1500 | 0.0047 | 0.0072 | 5.15E-01 | 2,611 | -0.0023 | 0.0143 | 8.70E-01 | 2,610 | -0.0013 | 0.0093 | 8.86E-01 | 2,611 |
| BMI, WC | cg15997518 | 1 | 81587462 | | | -0.0173 | 0.0114 | 1.29E-01 | 2,622 | -0.0552 | 0.0228 | 1.53E-02 | 2,621 | -0.0422 | 0.0148 | 4.27E-03 | 2,622 |
| BMI | cg03421440 | 1 | 92187108 | *BRDT* | TSS1500 | 0.0017 | 0.0111 | 8.78E-01 | 2,623 | 0.0009 | 0.0221 | 9.66E-01 | 2,622 | -0.0048 | 0.0143 | 7.36E-01 | 2,623 |
| BMI | cg03050965 | 1 | 101477825 | *S1PR1* | Body | -0.0147 | 0.0224 | 5.13E-01 | 2,623 | -0.0356 | 0.0447 | 4.27E-01 | 2,622 | -0.0276 | 0.0290 | 3.42E-01 | 2,623 |
| BMI | cg03725309 | 1 | 109559108 | *SARS* | Body | -0.0470 | 0.0135 | 4.98E-04 | 2,620 | -0.1160 | 0.0269 | **1.64E-05** | 2,619 | -0.0855 | 0.0174 | **9.50E-07** | 2,620 |
| BMI, WC | cg14476101 | 1 | 120057515 | *PHGDH* | Body | -0.0155 | 0.0085 | 6.86E-02 | 2,615 | -0.0394 | 0.0170 | 2.06E-02 | 2,614 | -0.0301 | 0.0110 | 6.33E-03 | 2,615 |
| BMI | cg18149207 | 1 | 150072258 | *RORC* | TSS1500 | 0.0186 | 0.0221 | 3.99E-01 | 2,623 | 0.0322 | 0.0442 | 4.66E-01 | 2,622 | 0.0354 | 0.0286 | 2.17E-01 | 2,623 |
| BMI | cg24678869 | 1 | 152186262 | *DENND4B* | TSS1500 | 0.0546 | 0.0236 | 2.10E-02 | 2,623 | 0.1432 | 0.0472 | 2.40E-03 | 2,622 | 0.0915 | 0.0306 | 2.77E-03 | 2,623 |
| BMI | cg23998749 | 1 | 153235405 | | | 0.0151 | 0.0195 | 4.40E-01 | 2,622 | 0.0573 | 0.0390 | 1.42E-01 | 2,621 | 0.0171 | 0.0253 | 4.98E-01 | 2,622 |
| BMI, Childhood/Young Adult Obesity, WC | cg12593793 | 1 | 154340759 | | | -0.0233 | 0.0137 | 8.83E-02 | 2,622 | -0.0806 | 0.0273 | 3.15E-03 | 2,621 | -0.0452 | 0.0177 | 1.07E-02 | 2,622 |
| BMI | cg25217710 | 1 | 154876147 | | | 0.0105 | 0.0120 | 3.81E-01 | 2,623 | 0.0657 | 0.0240 | 6.18E-03 | 2,622 | 0.0178 | 0.0156 | 2.54E-01 | 2,623 |
| BMI | cg04869770 | 1 | 162828174 | *PBX1* | Body | -0.0200 | 0.0182 | 2.70E-01 | 2,620 | -0.0333 | 0.0363 | 3.59E-01 | 2,619 | -0.0459 | 0.0235 | 5.10E-02 | 2,620 |
| BMI | cg09554443 | 1 | 165754386 | *CD247* | 1stExon/5UTR | -0.0361 | 0.0127 | 4.36E-03 | 2,623 | -0.0554 | 0.0253 | 2.87E-02 | 2,622 | -0.0117 | 0.0164 | 4.78E-01 | 2,623 |
| BMI | cg22534374 | 1 | 199778233 | | | -0.0204 | 0.0125 | 1.01E-01 | 2,623 | -0.0454 | 0.0248 | 6.77E-02 | 2,622 | -0.0267 | 0.0161 | 9.83E-02 | 2,623 |
| BMI | cg23172671 | 1 | 201749146 | | | 0.0039 | 0.0093 | 6.75E-01 | 2,619 | 0.0094 | 0.0187 | 6.13E-01 | 2,618 | 0.0141 | 0.0121 | 2.43E-01 | 2,619 |
| WC | cg13558971 | 1 | 201863708 | *ATP2B4* | 5UTR | -0.2552 | 0.1365 | 6.16E-02 | 2,620 | -0.2455 | 0.2728 | 3.68E-01 | 2,619 | -0.2428 | 0.1770 | 1.70E-01 | 2,620 |
| BMI | cg12458003 | 1 | 203227449 | *NFASC* | Body | 0.0525 | 0.0194 | 6.65E-03 | 2,623 | 0.1032 | 0.0386 | 7.58E-03 | 2,622 | 0.0786 | 0.0251 | 1.70E-03 | 2,623 |
| BMI | cg01455178 | 1 | 203914810 | *SLC45A3* | 5UTR | -0.0065 | 0.0150 | 6.64E-01 | 2,623 | 0.0084 | 0.0300 | 7.80E-01 | 2,622 | -0.0023 | 0.0195 | 9.04E-01 | 2,623 |
| BMI | cg10717869 | 1 | 204047535 | *SLC41A1* | 5UTR | 0.0143 | 0.0186 | 4.43E-01 | 2,623 | 0.0343 | 0.0371 | 3.56E-01 | 2,622 | 0.0247 | 0.0241 | 3.05E-01 | 2,623 |
| BMI | cg15323828 | 1 | 224120296 | *TMEM63A* | Body | -0.0097 | 0.0146 | 5.08E-01 | 2,619 | -0.0061 | 0.0293 | 8.35E-01 | 2,618 | -0.0040 | 0.0190 | 8.35E-01 | 2,619 |
| BMI | cg01101459 | 1 | 232938100 | | | 0.0013 | 0.0131 | 9.24E-01 | 2,623 | 0.0305 | 0.0261 | 2.43E-01 | 2,622 | 0.0088 | 0.0169 | 6.05E-01 | 2,623 |
| BMI, WC | cg00851028 | 1 | 232972395 | | | 0.0342 | 0.0158 | 3.03E-02 | 2,623 | 0.0884 | 0.0315 | 4.93E-03 | 2,622 | 0.0720 | 0.0204 | 4.12E-04 | 2,623 |
| BMI, WC | cg13139542 | 2 | 8160266 | | | 0.0382 | 0.0267 | 1.53E-01 | 2,622 | 0.0753 | 0.0533 | 1.58E-01 | 2,621 | 0.0558 | 0.0346 | 1.06E-01 | 2,622 |
| BMI | cg17214023 | 2 | 10094199 | | | -0.0225 | 0.0132 | 8.84E-02 | 2,621 | -0.0279 | 0.0264 | 2.89E-01 | 2,620 | -0.0214 | 0.0171 | 2.10E-01 | 2,621 |
| BMI | cg02560388 | 2 | 11887409 | | | -0.0328 | 0.0138 | 1.78E-02 | 2,623 | -0.0592 | 0.0276 | 3.18E-02 | 2,622 | -0.0278 | 0.0179 | 1.21E-01 | 2,623 |
| BMI, Childhood/Young Adult Obesity | cg04011474 | 2 | 28757959 | | | -0.0258 | 0.0149 | 8.35E-02 | 2,623 | -0.1109 | 0.0297 | 1.83E-04 | 2,622 | -0.0684 | 0.0192 | 3.76E-04 | 2,623 |
| BMI | cg16163382 | 2 | 37792144 | | | -0.0128 | 0.0234 | 5.85E-01 | 2,621 | -0.0752 | 0.0468 | 1.08E-01 | 2,620 | -0.0322 | 0.0304 | 2.90E-01 | 2,621 |
| BMI | cg19017142 | 2 | 64284749 | | | -0.0080 | 0.0098 | 4.12E-01 | 2,620 | -0.0440 | 0.0195 | 2.40E-02 | 2,619 | -0.0299 | 0.0127 | 1.80E-02 | 2,620 |
| BMI | cg26164488 | 2 | 64293799 | | | 0.0107 | 0.0115 | 3.53E-01 | 2,623 | -0.0133 | 0.0230 | 5.64E-01 | 2,622 | -0.0294 | 0.0149 | 4.84E-02 | 2,623 |
| BMI | cg26253134 | 2 | 70605229 | *TGFA* | Body | -0.0078 | 0.0110 | 4.78E-01 | 2,623 | -0.0234 | 0.0220 | 2.87E-01 | 2,622 | -0.0247 | 0.0142 | 8.31E-02 | 2,623 |
| BMI | cg00234616 | 2 | 74594080 | *TLX2* | TSS1500 | 0.0319 | 0.0142 | 2.49E-02 | 2,622 | 0.0712 | 0.0284 | 1.20E-02 | 2,621 | 0.0530 | 0.0184 | 3.96E-03 | 2,622 |
| BMI | cg07622521 | 2 | 85846133 | *ATOH8* | Body | 0.0154 | 0.0201 | 4.44E-01 | 2,622 | 0.0655 | 0.0401 | 1.02E-01 | 2,621 | 0.0470 | 0.0260 | 7.05E-02 | 2,622 |
| BMI | cg14017402 | 2 | 86079113 | | | 0.0374 | 0.0135 | 5.70E-03 | 2,622 | 0.0732 | 0.0270 | 6.69E-03 | 2,621 | 0.0439 | 0.0175 | 1.22E-02 | 2,622 |
| BMI | cg23417875 | 2 | 101679501 | *MAP4K4* | TSS1500 | 0.0198 | 0.0144 | 1.69E-01 | 2,622 | 0.0576 | 0.0287 | 4.51E-02 | 2,621 | 0.0397 | 0.0186 | 3.31E-02 | 2,622 |
| BMI | cg21282997 | 2 | 102402267 | *IL18RAP* | 5UTR | 0.0194 | 0.0106 | 6.76E-02 | 2,617 | 0.0525 | 0.0212 | 1.34E-02 | 2,616 | 0.0122 | 0.0138 | 3.75E-01 | 2,617 |

| | | | | | | | | | | | | | | | | | |
|---|---|---|---|---|---|---|---|---|---|---|---|---|---|---|---|---|---|
| BMI, WC | cg00585790 | 2 | 108571277 | *LIMS1* | 1stExon/5UTR | 0.0134 | 0.0126 | 2.90E-01 | 2,623 | 0.0503 | 0.0252 | 4.56E-02 | 2,622 | 0.0425 | 0.0163 | 9.16E-03 | 2,623 |
| BMI, Childhood/Young Adult Obesity | cg09152259 | 2 | 127872584 | | | -0.0287 | 0.0112 | 1.02E-02 | 2,620 | -0.0564 | 0.0223 | 1.15E-02 | 2,619 | -0.0414 | 0.0145 | 4.21E-03 | 2,620 |
| Childhood/Young Adult Obesity | cg06202737 | 2 | 127882749 | | | -0.0197 | 0.0107 | 6.51E-02 | 2,623 | -0.0455 | 0.0213 | 3.24E-02 | 2,622 | -0.0275 | 0.0138 | 4.61E-02 | 2,623 |
| BMI | cg15357118 | 2 | 128644442 | *UGGT1* | Body | 0.0037 | 0.0152 | 8.06E-01 | 2,623 | 0.0245 | 0.0303 | 4.19E-01 | 2,622 | 0.0284 | 0.0196 | 1.47E-01 | 2,623 |
| WC | cg25349939 | 2 | 144571555 | *GTDC1* | Body | 0.0329 | 0.0147 | 2.58E-02 | 2,621 | 0.0542 | 0.0294 | 6.54E-02 | 2,620 | 0.0105 | 0.0191 | 5.81E-01 | 2,621 |
| BMI | cg03327570 | 2 | 145021353 | | | -0.0796 | 0.0209 | 1.34E-04 | 2,620 | -0.1643 | 0.0416 | **7.84E-05** | 2,619 | -0.0629 | 0.0270 | 2.01E-02 | 2,620 |
| BMI | cg08035749 | 2 | 146233804 | | | 0.0510 | 0.0290 | 7.89E-02 | 2,621 | 0.1258 | 0.0579 | 2.98E-02 | 2,620 | 0.0585 | 0.0376 | 1.20E-01 | 2,621 |
| BMI | cg16721489 | 2 | 164901076 | | | -0.0126 | 0.0122 | 3.03E-01 | 2,616 | -0.0638 | 0.0243 | 8.65E-03 | 2,615 | -0.0468 | 0.0157 | 2.93E-03 | 2,616 |
| BMI | cg11775828 | 2 | 168809161 | *STK39* | Body | 0.0132 | 0.0133 | 3.21E-01 | 2,622 | 0.0316 | 0.0266 | 2.35E-01 | 2,621 | 0.0076 | 0.0173 | 6.62E-01 | 2,622 |
| BMI, Childhood/Young Adult Obesity | cg17178175 | 2 | 177818219 | *NFE2L2* | Body/5UTR | -0.0050 | 0.0133 | 7.07E-01 | 2,622 | -0.0118 | 0.0266 | 6.57E-01 | 2,621 | 0.0023 | 0.0173 | 8.95E-01 | 2,622 |
| BMI | cg09613192 | 2 | 181096783 | | | 0.0397 | 0.0113 | 4.44E-04 | 2,619 | 0.0770 | 0.0226 | 6.40E-04 | 2,618 | 0.0424 | 0.0146 | 3.80E-03 | 2,619 |
| BMI, WC | cg05918312 | 2 | 183078329 | *PDE1A* | Body | -0.0217 | 0.0129 | 9.26E-02 | 2,620 | -0.0622 | 0.0257 | 1.54E-02 | 2,619 | -0.0453 | 0.0167 | 6.56E-03 | 2,620 |
| BMI | cg12760041 | 2 | 210745135 | *C2orf67* | TSS1500 | 0.0686 | 0.0385 | 7.47E-02 | 2,613 | 0.1492 | 0.0769 | 5.23E-02 | 2,612 | 0.0723 | 0.0499 | 1.47E-01 | 2,613 |
| BMI | cg00634542 | 2 | 218962832 | *SLC11A1* | Body | 0.0209 | 0.0097 | 3.10E-02 | 2,619 | 0.0583 | 0.0193 | 2.56E-03 | 2,618 | 0.0213 | 0.0125 | 8.89E-02 | 2,619 |
| BMI | cg06096336 | 2 | 231698044 | *PSMD1;HTR2B* | Body;1stExon/5UTR | 0.0102 | 0.0091 | 2.65E-01 | 2,622 | 0.0339 | 0.0182 | 6.28E-02 | 2,621 | 0.0130 | 0.0118 | 2.72E-01 | 2,622 |
| BMI | cg04286697 | 2 | 231967867 | *B3GNT7* | TSS1500 | 0.0553 | 0.0180 | 2.18E-03 | 2,623 | 0.1382 | 0.0360 | **1.22E-04** | 2,622 | 0.0428 | 0.0234 | 6.74E-02 | 2,623 |
| BMI | cg20954977 | 2 | 231968360 | *B3GNT7* | TSS1500 | 0.0190 | 0.0092 | 3.77E-02 | 2,623 | 0.0332 | 0.0183 | 6.90E-02 | 2,622 | 0.0131 | 0.0119 | 2.71E-01 | 2,623 |
| BMI, WC | cg20034202 | 2 | 231968380 | *B3GNT7* | TSS200 | 0.0269 | 0.0114 | 1.80E-02 | 2,623 | 0.0649 | 0.0227 | 4.21E-03 | 2,622 | 0.0347 | 0.0147 | 1.83E-02 | 2,623 |
| BMI | cg12001357 | 2 | 233119096 | *CHRNG* | 3UTR | -0.0239 | 0.0132 | 7.10E-02 | 2,623 | -0.0576 | 0.0264 | 2.88E-02 | 2,622 | -0.0052 | 0.0171 | 7.62E-01 | 2,623 |
| BMI | cg06118217 | 2 | 239765935 | *HDAC4* | Body | 0.1127 | 0.0492 | 2.19E-02 | 2,623 | 0.1732 | 0.0982 | 7.78E-02 | 2,622 | 0.0938 | 0.0637 | 1.41E-01 | 2,623 |
| BMI | cg00144180 | 2 | 239959299 | *HDAC4* | 5UTR | 0.0192 | 0.0157 | 2.20E-01 | 2,616 | 0.0538 | 0.0313 | 8.49E-02 | 2,615 | 0.0388 | 0.0203 | 5.55E-02 | 2,616 |
| BMI | cg23032421 | 3 | 3127038 | *IL5RA* | 5UTR/1stExon | -0.0248 | 0.0142 | 8.10E-02 | 2,622 | -0.0865 | 0.0284 | 2.31E-03 | 2,621 | -0.0239 | 0.0184 | 1.95E-01 | 2,622 |
| BMI | cg04924511 | 3 | 10309731 | *GHRLOS;GHRL* | Body;TSS200 | 0.0303 | 0.0144 | 3.51E-02 | 2,619 | 0.0825 | 0.0287 | 3.98E-03 | 2,618 | 0.0579 | 0.0186 | 1.82E-03 | 2,619 |
| BMI | cg05628049 | 3 | 42088628 | | | 0.0226 | 0.0125 | 7.17E-02 | 2,622 | 0.0425 | 0.0250 | 8.99E-02 | 2,621 | 0.0416 | 0.0162 | 1.03E-02 | 2,622 |
| BMI, WC | cg18030453 | 3 | 45481220 | *LARS2* | Body | 0.0291 | 0.0188 | 1.22E-01 | 2,621 | 0.0954 | 0.0375 | 1.10E-02 | 2,620 | 0.0758 | 0.0243 | 1.84E-03 | 2,621 |
| BMI | cg00138407 | 3 | 47361509 | *KLHL18* | 3UTR | 0.0480 | 0.0162 | 2.99E-03 | 2,623 | 0.1195 | 0.0323 | 2.12E-04 | 2,622 | 0.0585 | 0.0209 | 5.21E-03 | 2,623 |
| BMI | cg17641710 | 3 | 50254042 | *GNAI2* | Body | 0.0183 | 0.0174 | 2.93E-01 | 2,621 | 0.0567 | 0.0347 | 1.02E-01 | 2,620 | 0.0135 | 0.0225 | 5.49E-01 | 2,621 |
| Childhood/Young Adult Obesity | cg21585138 | 3 | 50620110 | *CISH* | Body | -0.0222 | 0.0138 | 1.06E-01 | 2,618 | -0.0753 | 0.0274 | 6.08E-03 | 2,617 | -0.0501 | 0.0178 | 4.90E-03 | 2,618 |
| Childhood/Young Adult Obesity | cg23005227 | 3 | 50620430 | *CISH* | Body | -0.0244 | 0.0133 | 6.62E-02 | 2,623 | -0.0766 | 0.0265 | 3.86E-03 | 2,622 | -0.0459 | 0.0172 | 7.59E-03 | 2,623 |
| BMI, WC | cg08996521 | 3 | 50624998 | *CISH* | TSS1500 | -0.0210 | 0.0221 | 3.43E-01 | 2,622 | -0.0133 | 0.0442 | 7.63E-01 | 2,621 | -0.0502 | 0.0287 | 7.98E-02 | 2,622 |
| Childhood/Young Adult Obesity | cg03929796 | 3 | 52206880 | *ALAS1* | TSS1500 | 0.0039 | 0.0114 | 7.34E-01 | 2,623 | -0.0054 | 0.0227 | 8.12E-01 | 2,622 | -0.0214 | 0.0147 | 1.47E-01 | 2,623 |
| BMI | cg00108715 | 3 | 52540055 | *NT5DC2* | Body;Body | 0.0132 | 0.0188 | 4.81E-01 | 2,622 | 0.0634 | 0.0374 | 9.02E-02 | 2,621 | -0.0106 | 0.0243 | 6.64E-01 | 2,622 |
| BMI | cg01368219 | 3 | 54974831 | *CACNA2D3* | Body | 0.0307 | 0.0148 | 3.80E-02 | 2,623 | 0.0596 | 0.0295 | 4.36E-02 | 2,622 | 0.0226 | 0.0192 | 2.39E-01 | 2,623 |
| BMI | cg22012981 | 3 | 58497729 | *ACOX2* | 5UTR | 0.0510 | 0.0224 | 2.28E-02 | 2,618 | 0.1616 | 0.0447 | 2.99E-04 | 2,617 | 0.0592 | 0.0290 | 4.16E-02 | 2,618 |
| BMI | cg10549088 | 3 | 64252194 | | | 0.0226 | 0.0120 | 6.02E-02 | 2,622 | 0.0420 | 0.0240 | 7.98E-02 | 2,621 | 0.0057 | 0.0156 | 7.13E-01 | 2,622 |
| BMI | cg23232188 | 3 | 123039233 | *EAF2* | Body | 0.0211 | 0.0167 | 2.07E-01 | 2,622 | 0.0681 | 0.0334 | 4.16E-02 | 2,621 | -0.0001 | 0.0217 | 9.95E-01 | 2,622 |
| BMI | cg25197194 | 3 | 130241477 | *CCDC48* | 3UTR | -0.0105 | 0.0110 | 3.41E-01 | 2,623 | -0.0004 | 0.0220 | 9.85E-01 | 2,622 | 0.0045 | 0.0142 | 7.51E-01 | 2,623 |
| BMI | cg07730360 | 3 | 130328316 | | | 0.0404 | 0.0162 | 1.25E-02 | 2,622 | 0.1185 | 0.0323 | 2.42E-04 | 2,621 | 0.0350 | 0.0210 | 9.51E-02 | 2,622 |
| BMI, WC | cg13180098 | 3 | 130729613 | *RHO* | TSS1500 | -0.0338 | 0.0205 | 9.89E-02 | 2,622 | -0.1151 | 0.0408 | 4.76E-03 | 2,621 | -0.0624 | 0.0265 | 1.84E-02 | 2,622 |
| BMI, Childhood/Young Adult Obesity | cg01671681 | 3 | 156904429 | *PLCH1* | 5UTR | -0.0520 | 0.0144 | 3.04E-04 | 2,622 | -0.1144 | 0.0287 | **6.93E-05** | 2,621 | -0.0910 | 0.0186 | **1.00E-06** | 2,622 |
| BMI | cg00673344 | 3 | 158290385 | | | -0.0067 | 0.0092 | 4.66E-01 | 2,622 | -0.0137 | 0.0183 | 4.55E-01 | 2,621 | -0.0045 | 0.0119 | 7.03E-01 | 2,622 |
| Childhood/Young Adult Obesity | cg14986890 | 3 | 159910329 | *RARRES1* | Body | 0.0042 | 0.0118 | 7.23E-01 | 2,623 | 0.0139 | 0.0236 | 5.56E-01 | 2,622 | 0.0068 | 0.0153 | 6.59E-01 | 2,623 |
| BMI | cg15721584 | 3 | 182809449 | *SOX2OT* | TSS1500 | 0.0072 | 0.0080 | 3.73E-01 | 2,617 | 0.0278 | 0.0160 | 8.31E-02 | 2,616 | 0.0043 | 0.0104 | 6.83E-01 | 2,617 |
| BMI | cg09831562 | 3 | 182809819 | *SOX2OT* | TSS1500 | 0.0174 | 0.0124 | 1.63E-01 | 2,619 | 0.0636 | 0.0248 | 1.03E-02 | 2,618 | 0.0163 | 0.0161 | 3.10E-01 | 2,619 |
| BMI | cg19266387 | 3 | 185078817 | *PARL* | Body | 0.0105 | 0.0106 | 3.19E-01 | 2,621 | 0.0351 | 0.0211 | 9.57E-02 | 2,620 | 0.0264 | 0.0137 | 5.31E-02 | 2,621 |
| BMI | cg10513161 | 3 | 185188421 | *ABCC5* | Body | 0.0564 | 0.0269 | 3.62E-02 | 2,621 | 0.1556 | 0.0537 | 3.79E-03 | 2,620 | 0.0900 | 0.0348 | 9.79E-03 | 2,621 |
| BMI | cg23679085 | 3 | 185375467 | *AP2M1* | 5UTR | -0.0533 | 0.1536 | 7.29E-01 | 2,623 | 0.1111 | 0.3066 | 7.17E-01 | 2,622 | -0.0478 | 0.1989 | 8.10E-01 | 2,623 |
| BMI | cg15548101 | 3 | 187561334 | *DGKG* | 5UTR | 0.0414 | 0.0156 | 8.09E-03 | 2,605 | 0.0702 | 0.0312 | 2.46E-02 | 2,604 | 0.0334 | 0.0203 | 1.00E-01 | 2,605 |
| BMI | cg18513344 | 3 | 197015695 | *MUC4* | Body | 0.0005 | 0.0116 | 9.66E-01 | 2,623 | -0.0068 | 0.0232 | 7.71E-01 | 2,622 | 0.0026 | 0.0150 | 8.65E-01 | 2,623 |
| BMI, Childhood/Young Adult Obesity | cg07094298 | 4 | 2717824 | *TNIP2* | Body | -0.0141 | 0.0111 | 2.03E-01 | 2,623 | -0.0610 | 0.0221 | 5.73E-03 | 2,622 | -0.0422 | 0.0143 | 3.18E-03 | 2,623 |
| Childhood/Young Adult Obesity | cg00741986 | 4 | 2718130 | *TNIP2* | Body | -0.0076 | 0.0150 | 6.14E-01 | 2,623 | -0.0416 | 0.0299 | 1.64E-01 | 2,622 | -0.0346 | 0.0194 | 7.46E-02 | 2,623 |
| BMI | cg10438589 | 4 | 14140591 | | | 0.0449 | 0.0130 | 5.65E-04 | 2,622 | 0.1031 | 0.0260 | **7.17E-05** | 2,621 | 0.0409 | 0.0169 | 1.53E-02 | 2,622 |
| BMI, WC | cg04332373 | 4 | 15388740 | *CD38* | TSS1500 | 0.0385 | 0.0153 | 1.19E-02 | 2,622 | 0.0833 | 0.0306 | 6.46E-03 | 2,621 | 0.0276 | 0.0199 | 1.64E-01 | 2,622 |
| BMI, WC | cg10094443 | 4 | 39206261 | *UGDH* | TSS1500 | 0.0368 | 0.0149 | 1.33E-02 | 2,622 | 0.0976 | 0.0297 | 9.93E-04 | 2,621 | 0.0568 | 0.0192 | 3.16E-03 | 2,622 |
| BMI | cg18500988 | 4 | 54654465 | | | 0.0210 | 0.0113 | 6.32E-02 | 2,623 | 0.0383 | 0.0226 | 8.96E-02 | 2,622 | 0.0319 | 0.0146 | 2.94E-02 | 2,623 |
| BMI | cg27577928 | 4 | 54670145 | | | 0.0210 | 0.0137 | 1.25E-01 | 2,621 | 0.0599 | 0.0273 | 2.86E-02 | 2,620 | 0.0420 | 0.0177 | 1.80E-02 | 2,621 |
| BMI | cg26542660 | 4 | 56508617 | *CEP135* | TSS1500 | -0.0206 | 0.0198 | 2.98E-01 | 2,620 | -0.0496 | 0.0396 | 2.11E-01 | 2,619 | -0.0360 | 0.0257 | 1.61E-01 | 2,620 |
| Childhood/Young Adult Obesity | cg02734358 | 4 | 90446097 | *GPRIN3* | 5UTR | -0.0006 | 0.0096 | 9.48E-01 | 2,622 | -0.0039 | 0.0191 | 8.40E-01 | 2,621 | -0.0155 | 0.0124 | 2.10E-01 | 2,622 |

| | | | | | | | | | | | | | | | | | |
|---|---|---|---|---|---|---|---|---|---|---|---|---|---|---|---|---|---|
| BMI | cg13084458 | 4 | 128773367 | *INTU* | TSS1500 | 0.0075 | 0.0168 | 6.54E-01 | 2,622 | 0.0453 | 0.0334 | 1.76E-01 | 2,621 | 0.0409 | 0.0217 | 5.94E-02 | 2,622 |
| BMI, WC | cg06690548 | 4 | 139382258 | *SLC7A11* | Body | -0.0888 | 0.0211 | **2.55E-05** | 2,619 | -0.1676 | 0.0421 | **6.90E-05** | 2,618 | -0.1249 | 0.0273 | **4.73E-06** | 2,619 |
| BMI, WC | cg01300684 | 4 | 141449568 | *SCOC* | 5UTR | 0.0130 | 0.0163 | 4.23E-01 | 2,623 | -0.0106 | 0.0325 | 7.44E-01 | 2,622 | -0.0207 | 0.0211 | 3.26E-01 | 2,623 |
| BMI, WC | cg24968721 | 4 | 154931070 | *SFRP2* | TSS1500 | 0.0334 | 0.0166 | 4.40E-02 | 2,622 | 0.0748 | 0.0331 | 2.37E-02 | 2,621 | 0.0492 | 0.0214 | 2.17E-02 | 2,622 |
| BMI | cg05119988 | 4 | 166470639 | *SC4MOL* | 5UTR | -0.0124 | 0.0123 | 3.12E-01 | 2,621 | -0.0256 | 0.0245 | 2.96E-01 | 2,620 | -0.0166 | 0.0159 | 2.96E-01 | 2,621 |
| BMI | cg15442888 | 4 | 176025714 | | | -0.0305 | 0.0174 | 7.99E-02 | 2,618 | -0.0926 | 0.0348 | 7.82E-03 | 2,617 | -0.0536 | 0.0226 | 1.77E-02 | 2,618 |
| BMI, WC | cg02846963 | 4 | 182667809 | | | -0.0223 | 0.0164 | 1.73E-01 | 2,622 | -0.0695 | 0.0327 | 3.37E-02 | 2,621 | -0.0472 | 0.0212 | 2.63E-02 | 2,622 |
| BMI | cg17287155 | 5 | 446347 | *AHRR* | Body | 0.0069 | 0.0201 | 7.32E-01 | 2,623 | 0.0005 | 0.0401 | 9.90E-01 | 2,622 | 0.0231 | 0.0260 | 3.74E-01 | 2,623 |
| BMI | cg06940720 | 5 | 1579929 | | | 0.0135 | 0.0167 | 4.19E-01 | 2,623 | 0.0596 | 0.0333 | 7.30E-02 | 2,622 | 0.0306 | 0.0216 | 1.57E-01 | 2,623 |
| BMI | cg11080651 | 5 | 10498523 | *ROPN1L* | Body | 0.0058 | 0.0163 | 7.23E-01 | 2,622 | 0.0171 | 0.0326 | 5.99E-01 | 2,621 | -0.0006 | 0.0211 | 9.77E-01 | 2,622 |
| BMI | cg13276570 | 5 | 10620643 | *ANKRD33B* | Body | 0.0004 | 0.0269 | 9.89E-01 | 2,621 | 0.0317 | 0.0538 | 5.55E-01 | 2,620 | -0.0043 | 0.0349 | 9.03E-01 | 2,621 |
| Childhood/Young Adult Obesity | cg22143698 | 5 | 10661058 | *ANKRD33B* | Body | 0.0206 | 0.0098 | 3.47E-02 | 2,623 | 0.0530 | 0.0195 | 6.60E-03 | 2,622 | 0.0413 | 0.0126 | 1.09E-03 | 2,623 |
| BMI | cg10179300 | 5 | 14200618 | *TRIO* | Body | 0.0000 | 0.0150 | 9.99E-01 | 2,623 | -0.0082 | 0.0299 | 7.83E-01 | 2,622 | -0.0225 | 0.0194 | 2.45E-01 | 2,623 |
| BMI | cg06820412 | 5 | 135414195 | *TGFBI* | Body | -0.0184 | 0.0345 | 5.94E-01 | 2,623 | -0.0501 | 0.0688 | 4.67E-01 | 2,622 | 0.0163 | 0.0447 | 7.16E-01 | 2,623 |
| BMI | cg09047573 | 5 | 137502804 | *NME5* | 5UTR | 0.0335 | 0.0127 | 8.09E-03 | 2,622 | 0.0694 | 0.0253 | 6.00E-03 | 2,621 | 0.0516 | 0.0164 | 1.63E-03 | 2,622 |
| BMI | cg12153755 | 5 | 141042755 | *ARAP3* | TSS1500 | 0.0243 | 0.0152 | 1.11E-01 | 2,619 | 0.0415 | 0.0304 | 1.72E-01 | 2,618 | 0.0299 | 0.0198 | 1.30E-01 | 2,619 |
| BMI | cg13305415 | 5 | 141677648 | *SPRY4* | Body/5UTR | 0.0126 | 0.0120 | 2.94E-01 | 2,622 | 0.0203 | 0.0240 | 3.98E-01 | 2,621 | -0.0035 | 0.0156 | 8.21E-01 | 2,622 |
| BMI, WC | cg15674825 | 5 | 150032542 | *MYOZ3* | Body | 0.0103 | 0.0121 | 3.96E-01 | 2,620 | 0.0340 | 0.0242 | 1.60E-01 | 2,619 | 0.0332 | 0.0157 | 3.40E-02 | 2,620 |
| BMI, WC | cg26403843 | 5 | 158566663 | *RNF145* | Body | 0.0197 | 0.0080 | 1.37E-02 | 2,623 | 0.0551 | 0.0159 | 5.38E-04 | 2,622 | 0.0100 | 0.0103 | 3.32E-01 | 2,623 |
| BMI | cg18307303 | 5 | 158690034 | *IL12B* | 1stExon/5UTR | 0.0807 | 0.0239 | 7.24E-04 | 2,623 | 0.1639 | 0.0476 | 5.82E-04 | 2,622 | 0.0808 | 0.0309 | 8.96E-03 | 2,623 |
| BMI | cg11927233 | 5 | 170749147 | *NPM1* | Body | 0.0078 | 0.0084 | 3.58E-01 | 2,623 | 0.0106 | 0.0169 | 5.30E-01 | 2,622 | -0.0212 | 0.0109 | 5.25E-02 | 2,623 |
| BMI | cg04483863 | 5 | 173104343 | | | 0.0239 | 0.0325 | 4.61E-01 | 2,623 | 0.0346 | 0.0648 | 5.93E-01 | 2,622 | -0.0130 | 0.0421 | 7.57E-01 | 2,623 |
| BMI | cg02286155 | 5 | 176758868 | | | 0.0412 | 0.0242 | 8.90E-02 | 2,622 | 0.1153 | 0.0483 | 1.70E-02 | 2,621 | 0.0453 | 0.0313 | 1.49E-01 | 2,622 |
| BMI | cg22590032 | 5 | 179983171 | *FLT4* | Body | 0.0113 | 0.0178 | 5.27E-01 | 2,623 | 0.0775 | 0.0355 | 2.90E-02 | 2,622 | 0.0125 | 0.0230 | 5.88E-01 | 2,623 |
| BMI, WC | cg03717755 | 6 | 16244518 | *MYLIP* | Body | 0.0491 | 0.0125 | **8.62E-05** | 2,619 | 0.1306 | 0.0250 | **1.65E-07** | 2,618 | 0.0552 | 0.0162 | 6.74E-04 | 2,619 |
| BMI | cg00094412 | 6 | 29700833 | *GABBR1* | Body | -0.0234 | 0.0152 | 1.23E-01 | 2,622 | -0.0685 | 0.0303 | 2.38E-02 | 2,621 | -0.0558 | 0.0196 | 4.47E-03 | 2,622 |
| BMI | cg14352682 | 6 | 30722147 | *C6orf136* | TSS1500 | 0.0021 | 0.0200 | 9.18E-01 | 2,621 | 0.0490 | 0.0400 | 2.21E-01 | 2,620 | -0.0075 | 0.0260 | 7.74E-01 | 2,621 |
| BMI | cg20117675 | 6 | 30749199 | *DHX16* | TSS1500 | -0.0048 | 0.0117 | 6.80E-01 | 2,623 | -0.0262 | 0.0234 | 2.63E-01 | 2,622 | 0.0037 | 0.0152 | 8.09E-01 | 2,623 |
| Childhood/Young Adult Obesity | cg11554650 | 6 | 30761170 | *KIAA1949* | Body/1stExon | -0.0438 | 0.0231 | 5.77E-02 | 2,623 | -0.0841 | 0.0461 | 6.81E-02 | 2,622 | -0.0660 | 0.0299 | 2.72E-02 | 2,623 |
| WC | cg23533285 | 6 | 31430327 | *HLA-B* | Body | 0.0222 | 0.0110 | 4.28E-02 | 2,623 | 0.0481 | 0.0219 | 2.82E-02 | 2,622 | 0.0096 | 0.0142 | 4.98E-01 | 2,623 |
| BMI, WC | cg25843003 | 6 | 31539291 | *HCP5* | 3UTR | 0.0066 | 0.0078 | 3.97E-01 | 2,620 | 0.0125 | 0.0156 | 4.25E-01 | 2,619 | -0.0087 | 0.0101 | 3.93E-01 | 2,620 |
| BMI, WC | cg00218406 | 6 | 31539386 | *HCP5* | 3UTR | 0.0016 | 0.0064 | 7.98E-01 | 2,619 | -0.0050 | 0.0128 | 6.94E-01 | 2,618 | -0.0157 | 0.0083 | 5.91E-02 | 2,619 |
| BMI, WC | cg13123009 | 6 | 31789861 | *LY6G6E;LY6G6D* | TSS200;TSS1500 | 0.0316 | 0.0168 | 5.96E-02 | 2,623 | 0.0927 | 0.0335 | 5.67E-03 | 2,622 | 0.0276 | 0.0218 | 2.05E-01 | 2,623 |
| BMI | cg16599983 | 6 | 32276947 | *NOTCH4* | Body | -0.0179 | 0.0170 | 2.93E-01 | 2,622 | -0.0716 | 0.0339 | 3.43E-02 | 2,621 | -0.0422 | 0.0220 | 5.49E-02 | 2,622 |
| BMI | cg23893346 | 6 | 32277856 | *NOTCH4* | Body | -0.0175 | 0.0221 | 4.28E-01 | 2,623 | -0.0625 | 0.0441 | 1.57E-01 | 2,622 | -0.0348 | 0.0286 | 2.25E-01 | 2,623 |
| BMI, WC | cg04797846 | 6 | 32470685 | *BTNL2* | Body | -0.0146 | 0.0194 | 4.51E-01 | 2,619 | -0.0699 | 0.0387 | 7.08E-02 | 2,618 | -0.0393 | 0.0251 | 1.17E-01 | 2,619 |
| WC | cg22940798 | 6 | 32913532 | *TAP2* | Body | -0.0177 | 0.0129 | 1.70E-01 | 2,623 | -0.0506 | 0.0257 | 4.94E-02 | 2,622 | -0.0494 | 0.0167 | 3.04E-03 | 2,623 |
| BMI, WC | cg08099136 | 6 | 32919229 | *PSMB8* | Body | 0.0050 | 0.0106 | 6.40E-01 | 2,618 | 0.0018 | 0.0211 | 9.31E-01 | 2,617 | -0.0198 | 0.0137 | 1.48E-01 | 2,618 |
| BMI, WC | cg01309328 | 6 | 32919231 | *PSMB8* | Body | 0.0033 | 0.0121 | 7.86E-01 | 2,622 | 0.0029 | 0.0242 | 9.06E-01 | 2,621 | -0.0130 | 0.0157 | 4.09E-01 | 2,622 |
| BMI, WC | cg08818207 | 6 | 32928333 | *TAP1* | Body | -0.0031 | 0.0108 | 7.72E-01 | 2,616 | -0.0104 | 0.0216 | 6.28E-01 | 2,615 | -0.0243 | 0.0140 | 8.18E-02 | 2,616 |
| BMI | cg09132634 | 6 | 33082100 | *HLA-DOA* | 3UTR | -0.0030 | 0.0194 | 8.77E-01 | 2,620 | -0.0324 | 0.0388 | 4.04E-01 | 2,619 | 0.0024 | 0.0250 | 9.23E-01 | 2,620 |
| BMI, WC | cg09572125 | 6 | 33508455 | *SYNGAP1* | Body | 0.0116 | 0.0114 | 3.09E-01 | 2,622 | 0.0549 | 0.0228 | 1.60E-02 | 2,621 | 0.0339 | 0.0148 | 2.18E-02 | 2,622 |
| BMI, WC | cg20118717 | 6 | 33508483 | *SYNGAP1* | Body | 0.0144 | 0.0143 | 3.13E-01 | 2,620 | 0.0346 | 0.0286 | 2.25E-01 | 2,619 | 0.0411 | 0.0185 | 2.62E-02 | 2,620 |
| BMI | cg22875823 | 6 | 33508521 | *SYNGAP1* | Body | -0.0012 | 0.0147 | 9.37E-01 | 2,613 | 0.0201 | 0.0293 | 4.93E-01 | 2,612 | 0.0158 | 0.0190 | 4.05E-01 | 2,613 |
| BMI, Childhood/Young Adult Obesity | cg03957124 | 6 | 37124847 | | | -0.0261 | 0.0171 | 1.27E-01 | 2,619 | -0.1008 | 0.0340 | 3.04E-03 | 2,618 | -0.0434 | 0.0221 | 4.92E-02 | 2,619 |
| BMI | cg08666707 | 6 | 41564419 | | | -0.0114 | 0.0186 | 5.38E-01 | 2,622 | -0.0490 | 0.0370 | 1.86E-01 | 2,621 | -0.0472 | 0.0240 | 4.91E-02 | 2,622 |
| Childhood/Young Adult Obesity | cg04163119 | 6 | 41861984 | *PRICKLE4;TOMM6* | Body;TSS1500 | -0.0140 | 0.0211 | 5.06E-01 | 2,623 | -0.0715 | 0.0421 | 8.94E-02 | 2,622 | -0.0380 | 0.0273 | 1.64E-01 | 2,623 |
| BMI | cg18120259 | 6 | 44002617 | *LOC100132354* | Body | -0.0284 | 0.0147 | 5.38E-02 | 2,623 | -0.0675 | 0.0294 | 2.16E-02 | 2,622 | -0.0460 | 0.0191 | 1.59E-02 | 2,623 |
| BMI | cg18862566 | 6 | 45499894 | *RUNX2* | Body | 0.0200 | 0.0125 | 1.11E-01 | 2,623 | 0.0422 | 0.0250 | 9.14E-02 | 2,622 | 0.0412 | 0.0162 | 1.11E-02 | 2,623 |
| BMI, WC | cg15835542 | 6 | 45817786 | | | 0.0254 | 0.0106 | 1.70E-02 | 2,620 | 0.0564 | 0.0212 | 7.93E-03 | 2,619 | 0.0288 | 0.0138 | 3.71E-02 | 2,620 |
| BMI | cg07800670 | 6 | 56873046 | *DST* | Body | -0.0025 | 0.0182 | 8.91E-01 | 2,623 | -0.0422 | 0.0364 | 2.46E-01 | 2,622 | -0.0438 | 0.0236 | 6.36E-02 | 2,623 |
| BMI, WC | cg13010621 | 6 | 92401444 | | | -0.0104 | 0.0164 | 5.25E-01 | 2,622 | -0.0715 | 0.0326 | 2.83E-02 | 2,621 | -0.0303 | 0.0212 | 1.53E-01 | 2,622 |
| BMI | cg17478979 | 6 | 149813843 | *ZC3H12D* | Body | 0.0182 | 0.0117 | 1.18E-01 | 2,621 | 0.0465 | 0.0232 | 4.53E-02 | 2,620 | 0.0227 | 0.0151 | 1.32E-01 | 2,621 |
| BMI | cg06012428 | 6 | 157518896 | *ARID1B* | Body | -0.0455 | 0.0198 | 2.15E-02 | 2,623 | -0.0761 | 0.0395 | 5.40E-02 | 2,622 | -0.0537 | 0.0256 | 3.61E-02 | 2,623 |
| BMI, Childhood/Young Adult Obesity | cg03940776 | 6 | 158410001 | *SYNJ2* | Body | -0.0090 | 0.0197 | 6.48E-01 | 2,621 | -0.0314 | 0.0392 | 4.24E-01 | 2,620 | -0.0330 | 0.0254 | 1.95E-01 | 2,621 |
| BMI | cg17501210 | 6 | 166890242 | *RPS6KA2* | Body | -0.0184 | 0.0078 | 1.83E-02 | 2,622 | -0.0343 | 0.0155 | 2.73E-02 | 2,621 | -0.0256 | 0.0101 | 1.10E-02 | 2,622 |
| BMI | cg04816311 | 7 | 1033176 | *C7orf50* | Body | 0.0503 | 0.0117 | **1.65E-05** | 2,623 | 0.1331 | 0.0232 | **1.04E-08** | 2,622 | 0.0682 | 0.0151 | **6.45E-06** | 2,623 |
| BMI | cg23647610 | 7 | 1324339 | | | -0.0345 | 0.0322 | 2.84E-01 | 2,623 | -0.0920 | 0.0642 | 1.52E-01 | 2,622 | -0.0790 | 0.0416 | 5.78E-02 | 2,623 |

| | | | | | | | | | | | | | | | | | |
|---|---|---|---|---|---|---|---|---|---|---|---|---|---|---|---|---|---|
| BMI | cg08972190 | 7 | 2105521 | *MAD1L1* | Body | 0.0406 | 0.0178 | 2.25E-02 | 2,623 | 0.0958 | 0.0355 | 6.93E-03 | 2,622 | 0.0483 | 0.0230 | 3.57E-02 | 2,623 |
| BMI | cg05095590 | 7 | 2105785 | *MAD1L1* | Body | 0.0168 | 0.0081 | 3.75E-02 | 2,600 | 0.0574 | 0.0161 | 3.52E-04 | 2,599 | 0.0218 | 0.0104 | 3.64E-02 | 2,600 |
| BMI, WC | cg09956615 | 7 | 2639515 | *TTYH3* | Body | 0.0133 | 0.0094 | 1.56E-01 | 2,623 | 0.0321 | 0.0187 | 8.63E-02 | 2,622 | 0.0292 | 0.0121 | 1.64E-02 | 2,623 |
| BMI | cg13290371 | 7 | 5484597 | *FBXL18* | 3UTR | 0.0250 | 0.0128 | 5.16E-02 | 2,622 | 0.0579 | 0.0256 | 2.38E-02 | 2,621 | 0.0307 | 0.0166 | 6.50E-02 | 2,622 |
| Childhood/Young Adult Obesity | cg00232092 | 7 | 5485413 | *FBXL18* | 3UTR | 0.0058 | 0.0140 | 6.80E-01 | 2,622 | 0.0087 | 0.0280 | 7.56E-01 | 2,621 | 0.0057 | 0.0182 | 7.52E-01 | 2,622 |
| BMI | cg24469729 | 7 | 27127045 | *HOXA3* | 5UTR/TSS1500 | 0.0317 | 0.0181 | 8.00E-02 | 2,623 | 0.0633 | 0.0361 | 7.97E-02 | 2,622 | 0.0181 | 0.0234 | 4.40E-01 | 2,623 |
| BMI | cg21429551 | 7 | 30602287 | *GARS* | Body | -0.0164 | 0.0070 | 1.87E-02 | 2,623 | -0.0437 | 0.0139 | 1.73E-03 | 2,622 | -0.0308 | 0.0090 | 6.48E-04 | 2,623 |
| BMI, Childhood/Young Adult Obesity | cg13134297 | 7 | 30704081 | | | -0.0188 | 0.0139 | 1.75E-01 | 2,621 | -0.0352 | 0.0277 | 2.03E-01 | 2,620 | -0.0050 | 0.0179 | 7.78E-01 | 2,621 |
| BMI | cg04577162 | 7 | 73305333 | *RFC2* | Body | 0.0332 | 0.0178 | 6.25E-02 | 2,623 | 0.0767 | 0.0356 | 3.11E-02 | 2,622 | 0.0271 | 0.0231 | 2.41E-01 | 2,623 |
| BMI | cg19566658 | 7 | 100304177 | *TRIP6* | Body | 0.0054 | 0.0169 | 7.51E-01 | 2,622 | 0.0255 | 0.0337 | 4.50E-01 | 2,621 | 0.0152 | 0.0219 | 4.88E-01 | 2,622 |
| BMI | cg03310939 | 7 | 101475450 | *CUX1* | Body | 0.0315 | 0.0117 | 7.02E-03 | 2,623 | 0.0792 | 0.0233 | 6.74E-04 | 2,622 | 0.0560 | 0.0151 | 2.12E-04 | 2,623 |
| BMI | cg22103219 | 7 | 101721612 | *SH2B2* | Body | -0.0369 | 0.0123 | 2.73E-03 | 2,623 | -0.1108 | 0.0245 | **6.22E-06** | 2,622 | -0.0737 | 0.0159 | **3.58E-06** | 2,623 |
| BMI, WC | cg15857470 | 7 | 104732484 | *SRPK2* | Body | -0.0120 | 0.0131 | 3.59E-01 | 2,620 | -0.0431 | 0.0261 | 9.86E-02 | 2,619 | -0.0264 | 0.0169 | 1.19E-01 | 2,620 |
| BMI | cg05720226 | 7 | 116573833 | *ST7;ST7OT2* | Body;TSS1500 | 0.0356 | 0.0207 | 8.59E-02 | 2,622 | 0.0428 | 0.0414 | 3.01E-01 | 2,621 | 0.0204 | 0.0268 | 4.46E-01 | 2,622 |
| BMI | cg12976145 | 7 | 117289988 | *CTTNBP2* | Body | -0.0070 | 0.0155 | 6.52E-01 | 2,620 | -0.0353 | 0.0309 | 2.53E-01 | 2,619 | -0.0326 | 0.0200 | 1.03E-01 | 2,620 |
| BMI | cg27269962 | 7 | 127328233 | *SND1* | Body | 0.0581 | 0.0239 | 1.48E-02 | 2,619 | 0.1527 | 0.0476 | 1.33E-03 | 2,618 | 0.0656 | 0.0309 | 3.37E-02 | 2,619 |
| BMI | cg01844514 | 7 | 149188054 | *ZNF862* | Body | 0.0197 | 0.0244 | 4.18E-01 | 2,619 | 0.0548 | 0.0486 | 2.60E-01 | 2,618 | 0.0118 | 0.0316 | 7.08E-01 | 2,619 |
| BMI | cg25435714 | 7 | 156776142 | | | -0.0020 | 0.0159 | 8.99E-01 | 2,622 | 0.0176 | 0.0318 | 5.80E-01 | 2,621 | 0.0020 | 0.0206 | 9.22E-01 | 2,622 |
| Childhood/Young Adult Obesity | cg21163717 | 8 | 21825849 | *DOK2* | Body | -0.0126 | 0.0184 | 4.91E-01 | 2,622 | -0.0425 | 0.0366 | 2.46E-01 | 2,621 | -0.0315 | 0.0238 | 1.84E-01 | 2,622 |
| BMI | cg17560136 | 8 | 21971456 | *EPB49* | 5UTR/TSS1500 | 0.0119 | 0.0152 | 4.34E-01 | 2,622 | 0.0343 | 0.0303 | 2.58E-01 | 2,621 | -0.0086 | 0.0197 | 6.61E-01 | 2,622 |
| BMI, Childhood/Young Adult Obesity | cg24531955 | 8 | 23210636 | *LOXL2* | 3UTR | -0.0272 | 0.0137 | 4.62E-02 | 2,623 | -0.0718 | 0.0273 | 8.44E-03 | 2,622 | -0.0531 | 0.0177 | 2.66E-03 | 2,623 |
| BMI | cg02571142 | 8 | 42353960 | *DKK4* | TSS200 | 0.0180 | 0.0112 | 1.07E-01 | 2,623 | 0.0343 | 0.0223 | 1.23E-01 | 2,622 | 0.0266 | 0.0145 | 6.55E-02 | 2,623 |
| BMI | cg05490029 | 8 | 79881570 | *IL7* | TSS1500 | 0.0077 | 0.0108 | 4.73E-01 | 2,623 | 0.0104 | 0.0215 | 6.28E-01 | 2,622 | -0.0027 | 0.0139 | 8.47E-01 | 2,623 |
| BMI, Childhood/Young Adult Obesity | cg19589396 | 8 | 104006550 | | | -0.0363 | 0.0135 | 7.11E-03 | 2,616 | -0.0976 | 0.0269 | 2.89E-04 | 2,615 | -0.0637 | 0.0175 | 2.66E-04 | 2,616 |
| BMI | cg07471614 | 8 | 125924333 | | | 0.0267 | 0.0255 | 2.93E-01 | 2,622 | 0.0230 | 0.0508 | 6.51E-01 | 2,621 | 0.0060 | 0.0329 | 8.55E-01 | 2,622 |
| BMI | cg26140475 | 8 | 126594740 | | | -0.0030 | 0.0153 | 8.46E-01 | 2,622 | -0.0161 | 0.0305 | 5.98E-01 | 2,621 | -0.0411 | 0.0198 | 3.75E-02 | 2,622 |
| Childhood/Young Adult Obesity | cg03183540 | 8 | 135000938 | | | -0.0130 | 0.0139 | 3.48E-01 | 2,622 | -0.0618 | 0.0277 | 2.58E-02 | 2,621 | -0.0476 | 0.0180 | 8.09E-03 | 2,622 |
| BMI | cg26952928 | 8 | 142299415 | *SLC45A4* | Body | 0.0532 | 0.0287 | 6.37E-02 | 2,622 | 0.1640 | 0.0572 | 4.12E-03 | 2,621 | 0.0703 | 0.0371 | 5.82E-02 | 2,622 |
| BMI | cg25392060 | 8 | 142366303 | | | 0.0571 | 0.0150 | 1.45E-04 | 2,623 | 0.1546 | 0.0299 | **2.40E-07** | 2,622 | 0.0672 | 0.0195 | 5.59E-04 | 2,623 |
| BMI | cg26361535 | 8 | 144647747 | *ZC3H3* | Body | 0.0057 | 0.0110 | 6.02E-01 | 2,623 | 0.0189 | 0.0219 | 3.87E-01 | 2,622 | 0.0007 | 0.0142 | 9.58E-01 | 2,623 |
| BMI | cg14286682 | 9 | 6318493 | *TPD52L3* | 5UTR/1stExon | -0.0144 | 0.0157 | 3.60E-01 | 2,623 | -0.0619 | 0.0313 | 4.81E-02 | 2,622 | -0.0492 | 0.0203 | 1.54E-02 | 2,623 |
| BMI | cg02716826 | 9 | 33437032 | *SUGT1P1;AQP3* | Body;Body | -0.0258 | 0.0091 | 4.80E-03 | 2,620 | -0.0593 | 0.0183 | 1.16E-03 | 2,619 | -0.0428 | 0.0118 | 3.02E-04 | 2,620 |
| BMI | cg13591783 | 9 | 74958688 | *ANXA1* | 5UTR | -0.0015 | 0.0101 | 8.83E-01 | 2,623 | -0.0236 | 0.0202 | 2.42E-01 | 2,622 | -0.0185 | 0.0131 | 1.58E-01 | 2,623 |
| BMI, WC | cg13840239 | 9 | 88408212 | | | -0.0212 | 0.0156 | 1.75E-01 | 2,622 | -0.0801 | 0.0311 | 1.01E-02 | 2,621 | -0.0544 | 0.0202 | 6.99E-03 | 2,622 |
| Childhood/Young Adult Obesity | cg03257930 | 9 | 109439114 | | | -0.0131 | 0.0156 | 4.01E-01 | 2,612 | -0.0356 | 0.0312 | 2.55E-01 | 2,611 | -0.0388 | 0.0202 | 5.50E-02 | 2,612 |
| BMI | cg14264316 | 9 | 133270624 | | | -0.0073 | 0.0120 | 5.42E-01 | 2,621 | -0.0135 | 0.0240 | 5.74E-01 | 2,620 | -0.0218 | 0.0155 | 1.61E-01 | 2,621 |
| Childhood/Young Adult Obesity | cg13823169 | 9 | 138896714 | | | -0.0018 | 0.0138 | 8.95E-01 | 2,622 | -0.0399 | 0.0275 | 1.47E-01 | 2,621 | -0.0384 | 0.0178 | 3.14E-02 | 2,622 |
| BMI | cg19695507 | 10 | 13566199 | *BEND7* | Body | 0.0193 | 0.0131 | 1.42E-01 | 2,623 | 0.0683 | 0.0261 | 9.02E-03 | 2,622 | 0.0446 | 0.0170 | 8.49E-03 | 2,623 |
| BMI, WC | cg00134210 | 10 | 14684138 | *FAM107B* | Body | -0.0548 | 0.0351 | 1.18E-01 | 2,623 | -0.1440 | 0.0700 | 3.97E-02 | 2,622 | -0.0867 | 0.0454 | 5.61E-02 | 2,623 |
| BMI | cg05176551 | 10 | 32741592 | | | 0.0226 | 0.0174 | 1.95E-01 | 2,622 | 0.0631 | 0.0348 | 6.96E-02 | 2,621 | 0.0398 | 0.0226 | 7.74E-02 | 2,622 |
| BMI | cg14333542 | 10 | 75317567 | | | 0.0134 | 0.0093 | 1.52E-01 | 2,620 | 0.0470 | 0.0186 | 1.16E-02 | 2,619 | 0.0162 | 0.0121 | 1.80E-01 | 2,620 |
| BMI | cg16578636 | 10 | 92977437 | *PCGF5* | Body | 0.0026 | 0.0060 | 6.63E-01 | 2,620 | 0.0080 | 0.0119 | 4.99E-01 | 2,619 | -0.0029 | 0.0077 | 7.07E-01 | 2,620 |
| BMI | cg15903032 | 10 | 101287595 | | | 0.0110 | 0.0122 | 3.67E-01 | 2,619 | 0.0290 | 0.0243 | 2.31E-01 | 2,618 | 0.0007 | 0.0157 | 9.64E-01 | 2,619 |
| BMI, WC | cg07504977 | 10 | 102121002 | | | 0.0329 | 0.0083 | **7.29E-05** | 2,623 | 0.0697 | 0.0166 | **2.61E-05** | 2,622 | 0.0226 | 0.0108 | 3.57E-02 | 2,623 |
| BMI | cg00431050 | 10 | 103975720 | *ELOVL3* | TSS1500 | -0.0356 | 0.0208 | 8.69E-02 | 2,623 | -0.0941 | 0.0415 | 2.34E-02 | 2,622 | -0.0492 | 0.0269 | 6.75E-02 | 2,623 |
| Childhood/Young Adult Obesity | cg09701700 | 10 | 104184833 | *MIR146B* | TSS1500 | -0.0134 | 0.0106 | 2.09E-01 | 2,623 | -0.0266 | 0.0212 | 2.11E-01 | 2,622 | -0.0289 | 0.0138 | 3.58E-02 | 2,623 |
| BMI | cg17782974 | 10 | 104396980 | *TRIM8* | Body | 0.0197 | 0.0100 | 4.86E-02 | 2,617 | 0.0508 | 0.0199 | 1.06E-02 | 2,616 | 0.0201 | 0.0129 | 1.20E-01 | 2,617 |
| BMI, WC | cg25104397 | 10 | 104525910 | *C10orf26* | 5UTR/Body/1stExon | 0.0212 | 0.0116 | 6.86E-02 | 2,622 | 0.0518 | 0.0232 | 2.59E-02 | 2,621 | 0.0451 | 0.0151 | 2.76E-03 | 2,622 |
| BMI | cg26955383 | 10 | 105208650 | *CALHM1* | TSS200 | 0.0475 | 0.0146 | 1.13E-03 | 2,621 | 0.1316 | 0.0291 | **5.97E-06** | 2,620 | 0.0576 | 0.0189 | 2.30E-03 | 2,621 |
| BMI | cg26878209 | 10 | 112365465 | | | 0.0494 | 0.0159 | 1.92E-03 | 2,622 | 0.0627 | 0.0318 | 4.86E-02 | 2,621 | 0.0610 | 0.0206 | 3.08E-03 | 2,622 |
| BMI | cg18954700 | 10 | 124210844 | *HTRA1* | TSS200 | 0.0087 | 0.0303 | 7.74E-01 | 2,619 | 0.0132 | 0.0604 | 8.27E-01 | 2,618 | -0.0166 | 0.0392 | 6.72E-01 | 2,619 |
| BMI | cg04726013 | 10 | 126213226 | *LHPP* | Body | 0.0393 | 0.0097 | **4.65E-05** | 2,622 | 0.0691 | 0.0193 | 3.39E-04 | 2,621 | 0.0471 | 0.0125 | 1.64E-04 | 2,622 |
| BMI | cg00244001 | 10 | 126326795 | *FAM53B* | Body | -0.0220 | 0.0153 | 1.51E-01 | 2,622 | -0.0569 | 0.0305 | 6.23E-02 | 2,621 | -0.0444 | 0.0198 | 2.48E-02 | 2,622 |
| BMI | cg00238353 | 10 | 129675527 | *PTPRE* | 5UTR | -0.0232 | 0.0131 | 7.79E-02 | 2,622 | -0.0469 | 0.0262 | 7.38E-02 | 2,621 | -0.0374 | 0.0170 | 2.77E-02 | 2,622 |
| WC | cg01971407 | 11 | 303624 | *IFITM1* | TSS1500 | 0.0130 | 0.0116 | 2.62E-01 | 2,623 | 0.0003 | 0.0232 | 9.91E-01 | 2,622 | -0.0089 | 0.0150 | 5.53E-01 | 2,623 |
| WC | cg23570810 | 11 | 305102 | *IFITM1* | Body | 0.0005 | 0.0081 | 9.50E-01 | 2,623 | -0.0208 | 0.0161 | 1.97E-01 | 2,622 | -0.0194 | 0.0105 | 6.31E-02 | 2,623 |
| BMI | cg10927968 | 11 | 1763909 | | | 0.0503 | 0.0142 | 4.10E-04 | 2,621 | 0.1188 | 0.0284 | **2.93E-05** | 2,620 | 0.0564 | 0.0185 | 2.23E-03 | 2,621 |
| BMI, WC | cg02743674 | 11 | 2390044 | *TRPM5* | Body | 0.0192 | 0.0188 | 3.09E-01 | 2,623 | -0.0042 | 0.0376 | 9.10E-01 | 2,622 | -0.0023 | 0.0244 | 9.24E-01 | 2,623 |

| | | | | | | | | | | | | | | | | | |
|---|---|---|---|---|---|---|---|---|---|---|---|---|---|---|---|---|---|
| BMI, WC | cg19936757 | 11 | 2408794 | | | -0.0120 | 0.0218 | 5.82E-01 | 2,623 | -0.0413 | 0.0434 | 3.41E-01 | 2,622 | -0.0377 | 0.0282 | 1.81E-01 | 2,623 |
| BMI | cg06603309 | 11 | 2680720 | *KCNQ1* | Body | -0.0142 | 0.0135 | 2.92E-01 | 2,621 | -0.0093 | 0.0269 | 7.29E-01 | 2,620 | 0.0079 | 0.0175 | 6.53E-01 | 2,621 |
| BMI, Childhood/Young Adult Obesity | cg17061862 | 11 | 9547007 | | | -0.0238 | 0.0112 | 3.33E-02 | 2,621 | -0.0480 | 0.0223 | 3.12E-02 | 2,620 | -0.0129 | 0.0145 | 3.74E-01 | 2,621 |
| Childhood/Young Adult Obesity | cg08239103 | 11 | 16580394 | | | -0.0293 | 0.0139 | 3.54E-02 | 2,617 | -0.0770 | 0.0278 | 5.67E-03 | 2,616 | -0.0589 | 0.0180 | 1.09E-03 | 2,617 |
| BMI, WC | cg17526229 | 11 | 20134322 | | | 0.0213 | 0.0138 | 1.22E-01 | 2,623 | 0.0434 | 0.0274 | 1.14E-01 | 2,622 | 0.0373 | 0.0178 | 3.64E-02 | 2,623 |
| BMI, WC | cg06734985 | 11 | 26806077 | | | -0.0228 | 0.0119 | 5.57E-02 | 2,623 | -0.0707 | 0.0238 | 2.97E-03 | 2,622 | -0.0523 | 0.0154 | 6.90E-04 | 2,623 |
| BMI, Childhood/Young Adult Obesity, BMI change | cg07136133 | 11 | 36378953 | *PRR5L* | 5UTR/TSS200/Body | -0.0051 | 0.0164 | 7.53E-01 | 2,623 | 0.0176 | 0.0327 | 5.89E-01 | 2,622 | -0.0118 | 0.0212 | 5.76E-01 | 2,623 |
| BMI | cg11376147 | 11 | 57017774 | *SLC43A1* | Body | -0.0921 | 0.0222 | **3.31E-05** | 2,622 | -0.2139 | 0.0442 | **1.33E-06** | 2,621 | -0.1324 | 0.0287 | **4.00E-06** | 2,622 |
| BMI | cg03433986 | 11 | 62234200 | *BSCL2* | TSS1500 | 0.0090 | 0.0147 | 5.42E-01 | 2,621 | 0.0074 | 0.0294 | 8.02E-01 | 2,620 | -0.0096 | 0.0191 | 6.14E-01 | 2,621 |
| Childhood/Young Adult Obesity | cg27209729 | 11 | 64185501 | *NRXN2* | Body | -0.0060 | 0.0096 | 5.33E-01 | 2,623 | -0.0200 | 0.0192 | 2.96E-01 | 2,622 | -0.0159 | 0.0124 | 2.00E-01 | 2,623 |
| BMI, WC | cg26661640 | 11 | 64215422 | *NRXN2* | Body | -0.0059 | 0.0159 | 7.10E-01 | 2,623 | -0.0407 | 0.0316 | 1.99E-01 | 2,622 | -0.0233 | 0.0205 | 2.57E-01 | 2,623 |
| BMI | cg26800893 | 11 | 66941172 | *ATPGD1* | 5UTR/Body | -0.1555 | 0.0347 | **7.35E-06** | 2,622 | -0.2408 | 0.0693 | 5.16E-04 | 2,621 | -0.1476 | 0.0450 | 1.04E-03 | 2,622 |
| BMI, WC, Obesity, Central Obesity | cg00574958 | 11 | 68364198 | *CPT1A* | 5UTR | -0.1538 | 0.0309 | **6.58E-07** | 2,622 | -0.4424 | 0.0614 | **5.81E-13** | 2,621 | -0.2667 | 0.0399 | **2.28E-11** | 2,622 |
| BMI | cg17058475 | 11 | 68364313 | *CPT1A* | 5UTR | -0.0493 | 0.0174 | 4.59E-03 | 2,621 | -0.1340 | 0.0347 | **1.11E-04** | 2,620 | -0.1040 | 0.0225 | **3.68E-06** | 2,621 |
| BMI | cg11152384 | 11 | 68690876 | | | -0.0012 | 0.0143 | 9.34E-01 | 2,623 | 0.0022 | 0.0286 | 9.37E-01 | 2,622 | -0.0088 | 0.0185 | 6.34E-01 | 2,623 |
| BMI | cg11261850 | 11 | 75998618 | | | -0.0023 | 0.0095 | 8.07E-01 | 2,621 | 0.0186 | 0.0190 | 3.28E-01 | 2,620 | -0.0036 | 0.0123 | 7.70E-01 | 2,621 |
| BMI | cg19574327 | 11 | 95971416 | | | -0.0073 | 0.0156 | 6.37E-01 | 2,623 | -0.0468 | 0.0311 | 1.33E-01 | 2,622 | -0.0312 | 0.0202 | 1.22E-01 | 2,623 |
| BMI | cg09777883 | 11 | 111598906 | | | 0.0286 | 0.0130 | 2.82E-02 | 2,623 | 0.0724 | 0.0260 | 5.40E-03 | 2,622 | 0.0327 | 0.0169 | 5.28E-02 | 2,623 |
| BMI | cg12978214 | 11 | 112650384 | *NCAM1* | Body | -0.0197 | 0.0145 | 1.74E-01 | 2,622 | -0.0641 | 0.0289 | 2.67E-02 | 2,621 | -0.0324 | 0.0188 | 8.48E-02 | 2,622 |
| BMI | cg17260706 | 11 | 118288089 | *BCL9L* | TSS1500 | -0.0115 | 0.0157 | 4.64E-01 | 2,623 | -0.0275 | 0.0312 | 3.79E-01 | 2,622 | -0.0247 | 0.0203 | 2.22E-01 | 2,623 |
| BMI, WC | cg07217499 | 12 | 2286600 | *CACNA1C* | Body | -0.0108 | 0.0135 | 4.22E-01 | 2,623 | -0.0456 | 0.0269 | 8.97E-02 | 2,622 | -0.0333 | 0.0174 | 5.56E-02 | 2,623 |
| BMI, Childhood/Young Adult Obesity | cg10601624 | 12 | 6274638 | | | -0.0202 | 0.0162 | 2.10E-01 | 2,621 | -0.0728 | 0.0323 | 2.42E-02 | 2,620 | -0.0359 | 0.0209 | 8.69E-02 | 2,621 |
| WC | cg06538684 | 12 | 12402490 | *LOH12CR2* | TSS1500/Body | 0.0035 | 0.0123 | 7.78E-01 | 2,623 | 0.0042 | 0.0245 | 8.64E-01 | 2,622 | -0.0245 | 0.0159 | 1.22E-01 | 2,623 |
| BMI, WC, Obesity | cg20399616 | 12 | 24947234 | *BCAT1* | Body | -0.0327 | 0.0247 | 1.86E-01 | 2,622 | -0.0301 | 0.0493 | 5.41E-01 | 2,621 | -0.0190 | 0.0320 | 5.52E-01 | 2,622 |
| BMI | cg06898549 | 12 | 39369857 | | | 0.0221 | 0.0109 | 4.18E-02 | 2,623 | 0.0422 | 0.0217 | 5.17E-02 | 2,622 | 0.0044 | 0.0141 | 7.54E-01 | 2,623 |
| BMI | cg06559575 | 12 | 51776619 | *IGFBP6* | TSS1500 | -0.0140 | 0.0168 | 4.05E-01 | 2,622 | -0.0232 | 0.0336 | 4.88E-01 | 2,621 | -0.0097 | 0.0218 | 6.55E-01 | 2,622 |
| BMI | cg09689944 | 12 | 54677219 | *SUOX* | TSS200 | 0.0494 | 0.0157 | 1.65E-03 | 2,622 | 0.1078 | 0.0313 | 5.80E-04 | 2,621 | 0.0727 | 0.0203 | 3.49E-04 | 2,622 |
| BMI | cg20722088 | 12 | 88267017 | *DUSP6* | 3UTR | 0.0421 | 0.0182 | 2.04E-02 | 2,623 | 0.1159 | 0.0362 | 1.37E-03 | 2,622 | 0.0271 | 0.0235 | 2.50E-01 | 2,623 |
| Childhood/Young Adult Obesity | cg06647068 | 12 | 103377404 | *CHST11* | Body | -0.0131 | 0.0094 | 1.64E-01 | 2,623 | -0.0320 | 0.0188 | 8.92E-02 | 2,622 | -0.0321 | 0.0122 | 8.48E-03 | 2,623 |
| BMI | cg08693490 | 12 | 115242279 | | | 0.0150 | 0.0108 | 1.66E-01 | 2,623 | 0.0355 | 0.0216 | 1.00E-01 | 2,622 | 0.0310 | 0.0140 | 2.71E-02 | 2,623 |
| BMI, WC | cg13708645 | 12 | 120458688 | *KDM2B* | Body | 0.0186 | 0.0097 | 5.62E-02 | 2,622 | 0.0431 | 0.0194 | 2.65E-02 | 2,621 | 0.0229 | 0.0126 | 6.91E-02 | 2,622 |
| BMI | cg08120831 | 12 | 121235139 | *LRRC43* | 5UTR/Body | 0.0344 | 0.0133 | 9.48E-03 | 2,623 | 0.0792 | 0.0265 | 2.79E-03 | 2,622 | 0.0641 | 0.0172 | 1.89E-04 | 2,623 |
| WC | cg05899984 | 12 | 123604716 | | | 0.0159 | 0.0226 | 4.81E-01 | 2,623 | 0.0600 | 0.0452 | 1.84E-01 | 2,622 | 0.0451 | 0.0293 | 1.23E-01 | 2,623 |
| BMI, Childhood/Young Adult Obesity | cg19750657 | 13 | 37833967 | *UFM1* | 3UTR | 0.0171 | 0.0096 | 7.43E-02 | 2,622 | 0.0380 | 0.0191 | 4.64E-02 | 2,621 | 0.0153 | 0.0124 | 2.16E-01 | 2,622 |
| BMI | cg26687842 | 13 | 39953491 | *LOC646982* | TSS1500 | 0.0184 | 0.0133 | 1.66E-01 | 2,623 | 0.0284 | 0.0266 | 2.86E-01 | 2,622 | 0.0408 | 0.0172 | 1.79E-02 | 2,623 |
| BMI | cg11650298 | 13 | 43588989 | | | -0.0250 | 0.0244 | 3.05E-01 | 2,621 | -0.0690 | 0.0488 | 1.57E-01 | 2,620 | -0.0085 | 0.0316 | 7.88E-01 | 2,621 |
| BMI | cg25215047 | 13 | 50631080 | | | -0.0296 | 0.0316 | 3.49E-01 | 2,623 | -0.0540 | 0.0631 | 3.92E-01 | 2,622 | -0.0116 | 0.0409 | 7.76E-01 | 2,623 |
| BMI | cg21390682 | 13 | 112791752 | *MCF2L* | Body | -0.0075 | 0.0145 | 6.05E-01 | 2,623 | -0.0415 | 0.0289 | 1.52E-01 | 2,622 | -0.0295 | 0.0187 | 1.15E-01 | 2,623 |
| BMI | cg19881557 | 14 | 20037266 | | | 0.0355 | 0.0164 | 2.99E-02 | 2,623 | 0.0823 | 0.0326 | 1.17E-02 | 2,622 | 0.0296 | 0.0212 | 1.63E-01 | 2,623 |
| BMI, WC | cg03508235 | 14 | 22515834 | *JUB* | 5UTR/Body/1stExon | -0.0309 | 0.0178 | 8.15E-02 | 2,623 | -0.0938 | 0.0354 | 8.10E-03 | 2,622 | -0.0563 | 0.0230 | 1.44E-02 | 2,623 |
| BMI | cg12917475 | 14 | 22845627 | *BCL2L2* | TSS1500 | 0.0337 | 0.0145 | 2.05E-02 | 2,621 | 0.0876 | 0.0290 | 2.54E-03 | 2,620 | 0.0621 | 0.0188 | 9.45E-04 | 2,621 |
| BMI | cg03523676 | 14 | 23610075 | *CPNE6* | TSS1500 | 0.0222 | 0.0162 | 1.72E-01 | 2,623 | 0.0637 | 0.0324 | 4.89E-02 | 2,622 | 0.0370 | 0.0210 | 7.84E-02 | 2,623 |
| BMI | cg13097800 | 14 | 46173890 | | | -0.0133 | 0.0150 | 3.75E-01 | 2,622 | -0.0393 | 0.0299 | 1.88E-01 | 2,621 | -0.0281 | 0.0194 | 1.47E-01 | 2,622 |
| BMI | cg26357885 | 14 | 64075957 | *HSPA2* | TSS1500 | -0.0199 | 0.0163 | 2.21E-01 | 2,623 | -0.0183 | 0.0325 | 5.74E-01 | 2,622 | -0.0116 | 0.0211 | 5.81E-01 | 2,623 |
| BMI | cg16398761 | 14 | 73289991 | *C14orf43* | 5UTR | -0.1141 | 0.0285 | **6.30E-05** | 2,621 | -0.2271 | 0.0569 | **6.55E-05** | 2,620 | -0.1113 | 0.0370 | 2.61E-03 | 2,621 |
| BMI, Childhood/Young Adult Obesity | cg10919522 | 14 | 73297194 | *C14orf43* | 5UTR/TSS1500 | -0.0100 | 0.0113 | 3.78E-01 | 2,623 | -0.0403 | 0.0226 | 7.50E-02 | 2,622 | -0.0366 | 0.0147 | 1.27E-02 | 2,623 |
| BMI | cg19818308 | 14 | 75016498 | | | 0.0234 | 0.0122 | 5.51E-02 | 2,622 | 0.0396 | 0.0243 | 1.04E-01 | 2,621 | 0.0171 | 0.0158 | 2.79E-01 | 2,622 |
| BMI | cg19998073 | 14 | 88148196 | *ZC3H14* | 3UTR | 0.0346 | 0.0181 | 5.62E-02 | 2,623 | 0.0700 | 0.0361 | 5.29E-02 | 2,622 | 0.0044 | 0.0235 | 8.50E-01 | 2,623 |
| BMI | cg10814005 | 14 | 90780794 | *GPR68* | 5UTR | -0.0156 | 0.0161 | 3.33E-01 | 2,622 | -0.0218 | 0.0321 | 4.97E-01 | 2,621 | -0.0218 | 0.0209 | 2.96E-01 | 2,622 |
| BMI | cg21498220 | 14 | 101469461 | | | -0.1437 | 0.0999 | 1.51E-01 | 2,622 | -0.3592 | 0.1996 | 7.19E-02 | 2,621 | -0.0950 | 0.1293 | 4.63E-01 | 2,622 |
| BMI | cg10044470 | 14 | 103937329 | | | 0.0228 | 0.0128 | 7.53E-02 | 2,623 | 0.0691 | 0.0255 | 6.84E-03 | 2,622 | 0.0159 | 0.0166 | 3.37E-01 | 2,623 |
| BMI, WC | cg00808648 | 14 | 104850955 | *PACS2* | TSS1500 | 0.0275 | 0.0120 | 2.15E-02 | 2,622 | 0.0650 | 0.0239 | 6.49E-03 | 2,621 | 0.0555 | 0.0155 | 3.34E-04 | 2,622 |
| BMI | cg10734665 | 15 | 23658503 | *ATP10A* | Body | 0.0072 | 0.0116 | 5.34E-01 | 2,623 | -0.0154 | 0.0231 | 5.03E-01 | 2,622 | -0.0115 | 0.0150 | 4.42E-01 | 2,623 |
| BMI | cg27184903 | 15 | 27073019 | *APBA2* | 5UTR | 0.0482 | 0.0146 | 9.87E-04 | 2,599 | 0.1407 | 0.0292 | **1.43E-06** | 2,598 | 0.0652 | 0.0190 | 5.95E-04 | 2,599 |
| BMI | cg07814318 | 15 | 29411876 | *KLF13* | Body | 0.0423 | 0.0118 | 3.28E-04 | 2,620 | 0.0867 | 0.0235 | 2.27E-04 | 2,619 | 0.0281 | 0.0153 | 6.61E-02 | 2,620 |
| BMI, WC | cg15159104 | 15 | 41597157 | *MAP1A* | 5UTR/1stExon | 0.0331 | 0.0134 | 1.37E-02 | 2,623 | 0.0940 | 0.0268 | 4.52E-04 | 2,622 | 0.0610 | 0.0174 | 4.51E-04 | 2,623 |
| BMI, WC | cg22107533 | 15 | 42815375 | *TRIM69* | TSS1500 | -0.0038 | 0.0104 | 7.14E-01 | 2,623 | -0.0021 | 0.0208 | 9.19E-01 | 2,622 | -0.0251 | 0.0135 | 6.31E-02 | 2,623 |
| WC | cg05439368 | 15 | 42815390 | *TRIM69* | TSS1500 | -0.0032 | 0.0102 | 7.54E-01 | 2,623 | -0.0074 | 0.0204 | 7.18E-01 | 2,622 | -0.0237 | 0.0132 | 7.29E-02 | 2,623 |

| | | | | | | | | | | | | | | | | | |
|---|---|---|---|---|---|---|---|---|---|---|---|---|---|---|---|---|---|
| BMI | cg21670987 | 15 | 48137724 | *ATP8B4* | Body | -0.0026 | 0.0108 | 8.11E-01 | 2,621 | -0.0337 | 0.0216 | 1.19E-01 | 2,620 | -0.0193 | 0.0140 | 1.68E-01 | 2,621 |
| BMI, WC | cg06192883 | 15 | 50341463 | *MYO5C* | Body | 0.0484 | 0.0113 | **1.87E-05** | 2,622 | 0.1306 | 0.0225 | **6.72E-09** | 2,621 | 0.0752 | 0.0146 | **2.82E-07** | 2,622 |
| BMI, WC | cg24340572 | 15 | 67674107 | | | 0.0196 | 0.0125 | 1.19E-01 | 2,623 | 0.0546 | 0.0250 | 2.91E-02 | 2,622 | 0.0395 | 0.0162 | 1.50E-02 | 2,623 |
| BMI | cg07728579 | 15 | 81272067 | *FSD2* | TSS1500 | 0.0243 | 0.0137 | 7.63E-02 | 2,623 | 0.0458 | 0.0274 | 9.40E-02 | 2,622 | 0.0252 | 0.0177 | 1.55E-01 | 2,623 |
| BMI | cg11592786 | 15 | 87334585 | | | -0.0124 | 0.0278 | 6.57E-01 | 2,622 | -0.0870 | 0.0555 | 1.17E-01 | 2,621 | -0.0356 | 0.0360 | 3.24E-01 | 2,622 |
| BMI | cg18568872 | 15 | 88407498 | *ZNF710* | 5UTR | 0.0399 | 0.0179 | 2.62E-02 | 2,616 | 0.1374 | 0.0357 | **1.19E-04** | 2,615 | 0.0454 | 0.0232 | 5.08E-02 | 2,616 |
| BMI | cg11183227 | 15 | 89256411 | *MAN2A2* | Body | 0.0267 | 0.0193 | 1.65E-01 | 2,620 | 0.1172 | 0.0384 | 2.26E-03 | 2,619 | 0.0723 | 0.0249 | 3.65E-03 | 2,620 |
| BMI | cg27614723 | 15 | 90200901 | *SLCO3A1* | Body | 0.0175 | 0.0122 | 1.52E-01 | 2,622 | 0.0335 | 0.0244 | 1.70E-01 | 2,621 | 0.0088 | 0.0158 | 5.79E-01 | 2,622 |
| BMI, WC | cg16003913 | 16 | 67072 | *MPG* | 5UTR/TSS1500/1stExon | 0.0172 | 0.0152 | 2.58E-01 | 2,622 | 0.0501 | 0.0303 | 9.79E-02 | 2,621 | 0.0391 | 0.0197 | 4.70E-02 | 2,622 |
| BMI | cg00973118 | 16 | 314571 | *AXIN1* | Body | 0.0204 | 0.0117 | 8.04E-02 | 2,623 | 0.0712 | 0.0232 | 2.19E-03 | 2,622 | 0.0411 | 0.0151 | 6.48E-03 | 2,623 |
| BMI | cg05063895 | 16 | 2013519 | | | -0.0328 | 0.0284 | 2.48E-01 | 2,623 | -0.0365 | 0.0568 | 5.20E-01 | 2,622 | -0.0289 | 0.0368 | 4.33E-01 | 2,623 |
| BMI | cg23813257 | 16 | 3055287 | *IL32* | TSS1500/TSS200 | 0.0025 | 0.0146 | 8.64E-01 | 2,623 | -0.0108 | 0.0293 | 7.11E-01 | 2,622 | -0.0067 | 0.0190 | 7.23E-01 | 2,623 |
| BMI | cg26680760 | 16 | 3095881 | | | 0.0436 | 0.0210 | 3.80E-02 | 2,623 | 0.1173 | 0.0419 | 5.11E-03 | 2,622 | 0.0578 | 0.0272 | 3.37E-02 | 2,623 |
| BMI | cg01352090 | 16 | 4043534 | *ADCY9* | Body | 0.0102 | 0.0141 | 4.68E-01 | 2,623 | 0.0525 | 0.0281 | 6.23E-02 | 2,622 | 0.0386 | 0.0183 | 3.43E-02 | 2,623 |
| BMI | cg03500056 | 16 | 8722008 | *ABAT* | 5UTR/TSS200/5UTR | 0.0242 | 0.0118 | 4.08E-02 | 2,623 | 0.0650 | 0.0236 | 5.90E-03 | 2,622 | 0.0170 | 0.0153 | 2.67E-01 | 2,623 |
| BMI | cg03746015 | 16 | 11048501 | *CLEC16A* | Body | -0.0096 | 0.0121 | 4.27E-01 | 2,622 | -0.0417 | 0.0241 | 8.39E-02 | 2,621 | -0.0176 | 0.0156 | 2.61E-01 | 2,622 |
| BMI, Childhood/Young Adult Obesity | cg06946797 | 16 | 11329910 | | | -0.0166 | 0.0111 | 1.34E-01 | 2,620 | -0.0719 | 0.0221 | 1.14E-03 | 2,619 | -0.0453 | 0.0143 | 1.57E-03 | 2,620 |
| BMI | cg09607047 | 16 | 14309658 | *MIR365-1* | TSS1500 | 0.0375 | 0.0218 | 8.64E-02 | 2,621 | 0.1201 | 0.0435 | 5.82E-03 | 2,620 | 0.0742 | 0.0283 | 8.67E-03 | 2,621 |
| BMI | cg26663590 | 16 | 28866811 | | | 0.0368 | 0.0143 | 9.81E-03 | 2,622 | 0.1251 | 0.0284 | **1.06E-05** | 2,621 | 0.0578 | 0.0185 | 1.73E-03 | 2,622 |
| BMI | cg08877257 | 16 | 29728093 | *MAZ* | Body | -0.0397 | 0.0176 | 2.41E-02 | 2,618 | -0.1090 | 0.0351 | 1.92E-03 | 2,617 | -0.0588 | 0.0228 | 9.88E-03 | 2,618 |
| BMI | cg00711896 | 16 | 30317552 | *ZNF48* | Body | 0.1124 | 0.0248 | **5.68E-06** | 2,623 | 0.2385 | 0.0494 | **1.40E-06** | 2,622 | 0.1083 | 0.0321 | 7.48E-04 | 2,623 |
| BMI% | cg04502490 | 16 | 30337212 | *ZNF771* | 3UTR/3UTR | -0.0139 | 0.0123 | 2.59E-01 | 2,621 | -0.0552 | 0.0246 | 2.52E-02 | 2,620 | -0.0180 | 0.0160 | 2.60E-01 | 2,621 |
| BMI, WC | cg01172150 | 16 | 30724944 | | | 0.0236 | 0.0125 | 5.92E-02 | 2,623 | 0.0669 | 0.0249 | 7.29E-03 | 2,622 | 0.0459 | 0.0162 | 4.56E-03 | 2,623 |
| BMI | cg01243823 | 16 | 49289713 | *NOD2* | Body | -0.0026 | 0.0065 | 6.90E-01 | 2,623 | -0.0200 | 0.0130 | 1.24E-01 | 2,622 | -0.0109 | 0.0084 | 1.96E-01 | 2,623 |
| BMI | cg00863378 | 16 | 55107258 | *BBS2* | Body | 0.0301 | 0.0172 | 8.01E-02 | 2,623 | 0.1056 | 0.0343 | 2.08E-03 | 2,622 | 0.0211 | 0.0223 | 3.44E-01 | 2,623 |
| BMI | cg09018739 | 16 | 55737608 | *CPNE2* | Body | 0.0210 | 0.0153 | 1.68E-01 | 2,623 | 0.0933 | 0.0304 | 2.14E-03 | 2,622 | 0.0510 | 0.0197 | 9.72E-03 | 2,623 |
| BMI | cg10922280 | 16 | 66591728 | *DPEP2* | TSS1500 | 0.0149 | 0.0134 | 2.67E-01 | 2,621 | 0.0655 | 0.0267 | 1.43E-02 | 2,620 | 0.0434 | 0.0173 | 1.23E-02 | 2,621 |
| BMI | cg26899718 | 16 | 66962298 | *SMPD3* | Body | 0.0225 | 0.0216 | 2.99E-01 | 2,620 | 0.0723 | 0.0431 | 9.37E-02 | 2,619 | 0.0414 | 0.0280 | 1.39E-01 | 2,620 |
| BMI | cg08305942 | 16 | 78249855 | | | -0.0165 | 0.0143 | 2.48E-01 | 2,623 | -0.0556 | 0.0285 | 5.08E-02 | 2,622 | -0.0265 | 0.0185 | 1.51E-01 | 2,623 |
| BMI | cg16739178 | 16 | 84028175 | | | 0.0329 | 0.0176 | 6.15E-02 | 2,623 | 0.1085 | 0.0351 | 2.02E-03 | 2,622 | 0.0642 | 0.0228 | 4.89E-03 | 2,623 |
| BMI | cg03159676 | 16 | 84158037 | | | 0.0267 | 0.0099 | 7.11E-03 | 2,623 | 0.0594 | 0.0198 | 2.70E-03 | 2,622 | 0.0270 | 0.0128 | 3.57E-02 | 2,623 |
| BMI | cg07955474 | 16 | 84493057 | *IRF8* | 5UTR | -0.0053 | 0.0109 | 6.23E-01 | 2,623 | -0.0306 | 0.0217 | 1.58E-01 | 2,622 | -0.0145 | 0.0141 | 3.04E-01 | 2,623 |
| BMI | cg07021906 | 16 | 86424334 | *SLC7A5* | Body | 0.0294 | 0.0138 | 3.35E-02 | 2,623 | 0.1062 | 0.0275 | **1.14E-04** | 2,622 | 0.0592 | 0.0179 | 9.25E-04 | 2,623 |
| BMI | cg08443038 | 16 | 87534378 | *CBFA2T3* | 5UTR/Body | -0.0106 | 0.0202 | 6.00E-01 | 2,620 | -0.0219 | 0.0403 | 5.87E-01 | 2,619 | -0.0119 | 0.0261 | 6.50E-01 | 2,620 |
| BMI | cg02426464 | 17 | 1428055 | *SLC43A2* | Body | -0.0242 | 0.0262 | 3.55E-01 | 2,622 | -0.0429 | 0.0522 | 4.11E-01 | 2,621 | -0.0441 | 0.0338 | 1.93E-01 | 2,622 |
| Childhood/Young Adult Obesity | cg06568880 | 17 | 2113333 | *SMG6* | Body | -0.0217 | 0.0273 | 4.26E-01 | 2,623 | -0.0592 | 0.0544 | 2.76E-01 | 2,622 | -0.0024 | 0.0353 | 9.45E-01 | 2,623 |
| BMI | cg09664445 | 17 | 2559156 | *KIAA0664* | 5UTR | 0.0245 | 0.0174 | 1.59E-01 | 2,621 | 0.0792 | 0.0347 | 2.24E-02 | 2,620 | 0.0396 | 0.0225 | 7.86E-02 | 2,621 |
| BMI | cg16611352 | 17 | 3766178 | *P2RX1* | 1stExon | 0.0284 | 0.0114 | 1.28E-02 | 2,623 | 0.0624 | 0.0228 | 6.20E-03 | 2,622 | 0.0461 | 0.0148 | 1.81E-03 | 2,623 |
| BMI | cg01798813 | 17 | 3853423 | | | 0.1051 | 0.0423 | 1.30E-02 | 2,622 | 0.2523 | 0.0844 | 2.79E-03 | 2,621 | 0.1211 | 0.0548 | 2.71E-02 | 2,622 |
| BMI | cg19217955 | 17 | 7064718 | *DLG4;ACADVL* | TSS1500;Body | -0.0669 | 0.0673 | 3.20E-01 | 2,622 | -0.0687 | 0.1344 | 6.09E-01 | 2,621 | -0.0961 | 0.0871 | 2.70E-01 | 2,622 |
| BMI | cg22695339 | 17 | 7732355 | *CHD3* | Body/TSS1500 | -0.0470 | 0.0285 | 9.95E-02 | 2,623 | -0.0966 | 0.0569 | 8.97E-02 | 2,622 | -0.0779 | 0.0369 | 3.48E-02 | 2,623 |
| BMI | cg25649826 | 17 | 20879332 | *USP22* | Body | 0.0242 | 0.0180 | 1.80E-01 | 2,623 | 0.0612 | 0.0360 | 8.88E-02 | 2,622 | 0.0254 | 0.0233 | 2.76E-01 | 2,623 |
| WC | cg15416179 | 17 | 21130452 | *MAP2K3* | Body | -0.0467 | 0.0300 | 1.20E-01 | 2,622 | -0.0997 | 0.0600 | 9.65E-02 | 2,621 | -0.0867 | 0.0389 | 2.58E-02 | 2,622 |
| Childhood/Young Adult Obesity | cg00760203 | 17 | 34508447 | *PLXDC1* | Body | 0.0242 | 0.0124 | 5.14E-02 | 2,622 | 0.0636 | 0.0248 | 1.01E-02 | 2,621 | 0.0446 | 0.0160 | 5.40E-03 | 2,622 |
| BMI | cg13274938 | 17 | 35747348 | *RARA* | Body | 0.0944 | 0.0232 | **4.57E-05** | 2,617 | 0.1820 | 0.0463 | **8.34E-05** | 2,616 | 0.1092 | 0.0300 | 2.73E-04 | 2,617 |
| BMI | cg24457403 | 17 | 37023863 | *KRT16* | TSS1500 | -0.0278 | 0.0140 | 4.78E-02 | 2,621 | -0.0713 | 0.0280 | 1.10E-02 | 2,620 | -0.0524 | 0.0182 | 3.94E-03 | 2,621 |
| BMI | cg01597398 | 17 | 37504253 | | | -0.0009 | 0.0138 | 9.50E-01 | 2,623 | -0.0013 | 0.0276 | 9.64E-01 | 2,622 | -0.0209 | 0.0179 | 2.43E-01 | 2,623 |
| BMI, WC | cg03078551 | 17 | 39011824 | | | -0.0494 | 0.0274 | 7.17E-02 | 2,622 | -0.0483 | 0.0548 | 3.79E-01 | 2,621 | -0.0368 | 0.0356 | 3.01E-01 | 2,622 |
| BMI | cg27050612 | 17 | 43488197 | *NFE2L1* | Body | -0.0253 | 0.0188 | 1.79E-01 | 2,623 | -0.0831 | 0.0376 | 2.70E-02 | 2,622 | -0.0634 | 0.0243 | 9.20E-03 | 2,623 |
| BMI | cg01130991 | 17 | 43865391 | | | 0.0220 | 0.0154 | 1.54E-01 | 2,623 | 0.0627 | 0.0308 | 4.19E-02 | 2,622 | 0.0264 | 0.0200 | 1.86E-01 | 2,623 |
| Childhood/Young Adult Obesity | cg05487507 | 17 | 44026860 | *LOC404266;HOXB5* | Body/TSS1500;TSS1500 | -0.0233 | 0.0116 | 4.41E-02 | 2,623 | -0.0453 | 0.0231 | 4.99E-02 | 2,622 | -0.0259 | 0.0150 | 8.36E-02 | 2,623 |
| BMI | cg14509967 | 17 | 44034134 | *HOXB6;LOC404266* | 5UTR;Body | -0.0053 | 0.0102 | 6.00E-01 | 2,621 | 0.0087 | 0.0203 | 6.67E-01 | 2,620 | -0.0140 | 0.0132 | 2.87E-01 | 2,621 |
| BMI | cg02650017 | 17 | 44656613 | *PHOSPHO1* | Body | -0.0763 | 0.0613 | 2.13E-01 | 2,622 | -0.2340 | 0.1223 | 5.56E-02 | 2,621 | -0.0588 | 0.0794 | 4.59E-01 | 2,622 |
| BMI, WC | cg21139312 | 17 | 53018224 | *MSI2* | Body | 0.0816 | 0.0303 | 6.97E-03 | 2,621 | 0.1873 | 0.0604 | 1.92E-03 | 2,620 | 0.1192 | 0.0392 | 2.34E-03 | 2,621 |
| BMI, WC, Childhood/Young Adult Obesity | cg24174557 | 17 | 55258326 | *TMEM49* | Body | -0.0198 | 0.0090 | 2.73E-02 | 2,623 | -0.0605 | 0.0179 | 7.28E-04 | 2,622 | -0.0353 | 0.0116 | 2.35E-03 | 2,623 |
| Childhood/Young Adult Obesity | cg16936953 | 17 | 55270447 | *TMEM49* | Body | 0.0120 | 0.0077 | 1.20E-01 | 2,623 | 0.0140 | 0.0154 | 3.63E-01 | 2,622 | -0.0041 | 0.0100 | 6.83E-01 | 2,623 |
| Childhood/Young Adult Obesity | cg12054453 | 17 | 55270499 | *TMEM49* | Body | 0.0139 | 0.0069 | 4.38E-02 | 2,621 | 0.0202 | 0.0138 | 1.45E-01 | 2,620 | 0.0023 | 0.0090 | 7.95E-01 | 2,621 |
| Childhood/Young Adult Obesity | cg01409343 | 17 | 55270522 | *TMEM49* | Body | 0.0060 | 0.0117 | 6.07E-01 | 2,623 | -0.0211 | 0.0234 | 3.68E-01 | 2,622 | -0.0252 | 0.0152 | 9.62E-02 | 2,623 |

| | | | | | | | | | | | | | | | | | |
|---|---|---|---|---|---|---|---|---|---|---|---|---|---|---|---|---|---|
| Childhood/Young Adult Obesity | cg18942579 | 17 | 55270555 | *TMEM49* | Body | 0.0033 | 0.0092 | 7.21E-01 | 2,622 | -0.0025 | 0.0183 | 8.93E-01 | 2,621 | -0.0134 | 0.0119 | 2.61E-01 | 2,622 |
| WCHT, BMI% | cg02988947 | 17 | 59132545 | *LIMD2* | TSS1500 | -0.0173 | 0.0085 | 4.10E-02 | 2,618 | -0.0341 | 0.0169 | 4.41E-02 | 2,617 | -0.0227 | 0.0110 | 3.84E-02 | 2,618 |
| BMI | cg07918509 | 17 | 59437139 | *ICAM2* | Body | 0.0318 | 0.0204 | 1.20E-01 | 2,622 | 0.0488 | 0.0408 | 2.32E-01 | 2,621 | 0.0488 | 0.0264 | 6.48E-02 | 2,622 |
| BMI, WC | cg18772573 | 17 | 68769575 | *CPSF4L* | 1stExon;5UTR | 0.0614 | 0.0193 | 1.49E-03 | 2,623 | 0.1273 | 0.0386 | 9.80E-04 | 2,622 | 0.0700 | 0.0251 | 5.24E-03 | 2,623 |
| BMI | cg08813944 | 17 | 68770184 | *CPSF4L* | TSS1500 | 0.0271 | 0.0145 | 6.11E-02 | 2,593 | 0.0613 | 0.0289 | 3.37E-02 | 2,592 | 0.0392 | 0.0187 | 3.64E-02 | 2,593 |
| BMI | cg21486834 | 17 | 71989137 | *RHBDF2* | Body | 0.0611 | 0.0233 | 8.65E-03 | 2,622 | 0.1678 | 0.0464 | 2.98E-04 | 2,621 | 0.1024 | 0.0301 | 6.69E-04 | 2,622 |
| BMI, Childhood/Young Adult Obesity, WCHT, BMI% | cg18181703 | 17 | 73866216 | *SOCS3* | Body | 0.0056 | 0.0109 | 6.09E-01 | 2,623 | -0.0225 | 0.0218 | 3.03E-01 | 2,622 | -0.0330 | 0.0141 | 1.96E-02 | 2,623 |
| BMI, Childhood/Young Adult Obesity | cg10508317 | 17 | 73866741 | *SOCS3* | Body | 0.0240 | 0.0244 | 3.26E-01 | 2,623 | -0.0154 | 0.0488 | 7.52E-01 | 2,622 | -0.0316 | 0.0316 | 3.17E-01 | 2,623 |
| BMI | cg27637521 | 17 | 73866797 | *SOCS3* | 5UTR | 0.0359 | 0.0296 | 2.26E-01 | 2,623 | 0.0429 | 0.0592 | 4.69E-01 | 2,622 | -0.0123 | 0.0384 | 7.49E-01 | 2,623 |
| BMI | cg27470213 | 17 | 74479290 | *LGALS3BP* | Body | -0.0282 | 0.0189 | 1.36E-01 | 2,622 | -0.0742 | 0.0377 | 4.90E-02 | 2,621 | -0.0401 | 0.0244 | 1.01E-01 | 2,622 |
| BMI, WC | cg11202345 | 17 | 74487652 | *LGALS3BP* | 1stExon/5UTR | 0.0175 | 0.0099 | 7.58E-02 | 2,622 | 0.0654 | 0.0197 | 8.68E-04 | 2,621 | 0.0182 | 0.0128 | 1.55E-01 | 2,622 |
| BMI, WC | cg04927537 | 17 | 74487686 | *LGALS3BP* | TSS200 | 0.0097 | 0.0082 | 2.33E-01 | 2,622 | 0.0381 | 0.0163 | 1.93E-02 | 2,621 | 0.0128 | 0.0106 | 2.26E-01 | 2,622 |
| BMI | cg22713958 | 17 | 74487840 | *LGALS3BP* | TSS200 | 0.0354 | 0.0208 | 8.92E-02 | 2,623 | 0.0819 | 0.0416 | 4.88E-02 | 2,622 | 0.0392 | 0.0270 | 1.45E-01 | 2,623 |
| BMI, WC | cg25178683 | 17 | 74487862 | *LGALS3BP* | TSS1500 | 0.0162 | 0.0108 | 1.34E-01 | 2,623 | 0.0526 | 0.0216 | 1.47E-02 | 2,622 | 0.0221 | 0.0140 | 1.14E-01 | 2,623 |
| BMI | cg17836612 | 17 | 74487952 | *LGALS3BP* | TSS1500 | 0.0071 | 0.0161 | 6.57E-01 | 2,623 | 0.0408 | 0.0321 | 2.03E-01 | 2,622 | 0.0137 | 0.0208 | 5.11E-01 | 2,623 |
| BMI | cg18091083 | 17 | 76433687 | *RPTOR* | Body | 0.0124 | 0.0092 | 1.81E-01 | 2,621 | 0.0312 | 0.0185 | 9.11E-02 | 2,620 | 0.0201 | 0.0120 | 9.37E-02 | 2,621 |
| BMI, WC | cg26766064 | 17 | 76714306 | *MIR657;AATK;MIR338* | TSS1500;Body;Body | -0.0136 | 0.0168 | 4.19E-01 | 2,623 | -0.0580 | 0.0335 | 8.36E-02 | 2,622 | -0.0368 | 0.0217 | 9.08E-02 | 2,623 |
| BMI | cg11969813 | 17 | 77409848 | *P4HB* | Body | 0.0419 | 0.0156 | 7.02E-03 | 2,619 | 0.1225 | 0.0310 | **7.71E-05** | 2,618 | 0.0382 | 0.0201 | 5.76E-02 | 2,619 |
| Childhood/Young Adult Obesity | cg23018755 | 17 | 77474822 | *MAFG* | 5UTR/TSS200 | 0.0352 | 0.0148 | 1.75E-02 | 2,623 | 0.0800 | 0.0296 | 6.87E-03 | 2,622 | 0.0613 | 0.0192 | 1.38E-03 | 2,623 |
| BMI | cg07012687 | 17 | 77788469 | *SLC16A3* | Body | 0.0278 | 0.0131 | 3.31E-02 | 2,621 | 0.1219 | 0.0260 | **2.78E-06** | 2,620 | 0.0469 | 0.0169 | 5.55E-03 | 2,621 |
| BMI | cg16755922 | 17 | 78129503 | *FOXK2* | Body | 0.0149 | 0.0131 | 2.53E-01 | 2,622 | 0.0412 | 0.0261 | 1.14E-01 | 2,621 | 0.0193 | 0.0169 | 2.53E-01 | 2,622 |
| BMI | cg15871086 | 18 | 54677575 | | | 0.0235 | 0.0191 | 2.17E-01 | 2,621 | 0.0429 | 0.0381 | 2.60E-01 | 2,620 | -0.0259 | 0.0247 | 2.95E-01 | 2,621 |
| WC | cg13348877 | 18 | 76106228 | *PARD6G* | 1stExon/5UTR | 0.0423 | 0.0435 | 3.31E-01 | 2,619 | 0.1391 | 0.0868 | 1.09E-01 | 2,618 | 0.0746 | 0.0563 | 1.85E-01 | 2,619 |
| BMI | cg24824917 | 19 | 1052716 | | | 0.0229 | 0.0156 | 1.41E-01 | 2,623 | 0.0406 | 0.0312 | 1.93E-01 | 2,622 | 0.0427 | 0.0202 | 3.43E-02 | 2,623 |
| BMI, Childhood/Young Adult Obesity | cg18608055 | 19 | 1081866 | *SBNO2* | Body | 0.0127 | 0.0127 | 3.19E-01 | 2,619 | 0.0169 | 0.0253 | 5.05E-01 | 2,618 | -0.0035 | 0.0165 | 8.32E-01 | 2,619 |
| BMI | cg04524040 | 19 | 4104364 | *CREB3L3* | TSS1500 | -0.0048 | 0.0069 | 4.85E-01 | 2,623 | -0.0080 | 0.0138 | 5.62E-01 | 2,622 | -0.0084 | 0.0090 | 3.49E-01 | 2,623 |
| BMI | cg22950899 | 19 | 5204997 | *PTPRS* | Body | 0.0323 | 0.0138 | 1.93E-02 | 2,618 | 0.0899 | 0.0276 | 1.11E-03 | 2,617 | 0.0384 | 0.0179 | 3.19E-02 | 2,618 |
| Childhood/Young Adult Obesity | cg03650189 | 19 | 10266083 | *ICAM5* | Body | 0.0081 | 0.0069 | 2.38E-01 | 2,623 | 0.0185 | 0.0137 | 1.76E-01 | 2,622 | 0.0137 | 0.0089 | 1.23E-01 | 2,623 |
| Childhood/Young Adult Obesity | cg15011409 | 19 | 10266226 | *ICAM5* | Body | 0.0068 | 0.0064 | 2.86E-01 | 2,623 | 0.0167 | 0.0127 | 1.88E-01 | 2,622 | 0.0109 | 0.0082 | 1.85E-01 | 2,623 |
| BMI | cg07769588 | 19 | 10516622 | *ATG4D* | Body | 0.0373 | 0.0131 | 4.34E-03 | 2,618 | 0.0727 | 0.0261 | 5.43E-03 | 2,617 | 0.0069 | 0.0170 | 6.84E-01 | 2,618 |
| BMI, WC | cg01581222 | 19 | 10819952 | *C19orf38* | TSS200 | 0.0104 | 0.0119 | 3.79E-01 | 2,622 | 0.0360 | 0.0237 | 1.28E-01 | 2,621 | 0.0312 | 0.0153 | 4.17E-02 | 2,622 |
| BMI | cg01751802 | 19 | 11170639 | *KANK2* | TSS1500 | 0.0209 | 0.0115 | 7.02E-02 | 2,623 | 0.0715 | 0.0230 | 1.89E-03 | 2,622 | 0.0332 | 0.0149 | 2.60E-02 | 2,623 |
| BMI, WC | cg24679890 | 19 | 17107356 | *MYO9B* | Body | 0.0250 | 0.0132 | 5.88E-02 | 2,623 | 0.0734 | 0.0264 | 5.36E-03 | 2,622 | 0.0483 | 0.0171 | 4.74E-03 | 2,623 |
| BMI, WC | cg20981127 | 19 | 17218587 | *NR2F6* | TSS1500 | 0.0236 | 0.0134 | 7.78E-02 | 2,622 | 0.0685 | 0.0267 | 1.04E-02 | 2,621 | 0.0469 | 0.0173 | 6.89E-03 | 2,622 |
| BMI | cg04557677 | 19 | 17820082 | *JAK3* | TSS1500 | -0.0401 | 0.0453 | 3.76E-01 | 2,623 | -0.0156 | 0.0904 | 8.63E-01 | 2,622 | -0.0810 | 0.0586 | 1.67E-01 | 2,623 |
| BMI | cg07682160 | 19 | 18820935 | *UPF1* | Body | 0.0249 | 0.0163 | 1.26E-01 | 2,622 | 0.0483 | 0.0325 | 1.37E-01 | 2,621 | 0.0202 | 0.0211 | 3.37E-01 | 2,622 |
| BMI | cg26950531 | 19 | 43396355 | *DPF1* | Body | -0.0079 | 0.0077 | 3.09E-01 | 2,607 | -0.0016 | 0.0155 | 9.19E-01 | 2,606 | 0.0061 | 0.0100 | 5.40E-01 | 2,607 |
| BMI | cg26836479 | 19 | 47398193 | *DEDD2* | Body | -0.0352 | 0.0221 | 1.11E-01 | 2,622 | -0.0995 | 0.0441 | 2.39E-02 | 2,621 | -0.0448 | 0.0286 | 1.17E-01 | 2,622 |
| BMI, Childhood/Young Adult Obesity | cg26470501 | 19 | 49944795 | *BCL3* | Body | -0.0152 | 0.0146 | 2.98E-01 | 2,623 | -0.0544 | 0.0291 | 6.17E-02 | 2,622 | -0.0588 | 0.0189 | 1.81E-03 | 2,623 |
| BMI | cg27087650 | 19 | 49947636 | *BCL3* | Body | -0.0228 | 0.0156 | 1.43E-01 | 2,622 | -0.0425 | 0.0311 | 1.71E-01 | 2,621 | -0.0533 | 0.0201 | 8.14E-03 | 2,622 |
| BMI | cg27146050 | 19 | 51493397 | *HIF3A* | Body/TSS200 | -0.0021 | 0.0094 | 8.22E-01 | 2,623 | 0.0064 | 0.0189 | 7.36E-01 | 2,622 | -0.0116 | 0.0122 | 3.44E-01 | 2,623 |
| BMI | cg22891070 | 19 | 51493482 | *HIF3A* | Body/TSS200 | -0.0006 | 0.0056 | 9.19E-01 | 2,621 | -0.0010 | 0.0111 | 9.27E-01 | 2,620 | -0.0077 | 0.0072 | 2.82E-01 | 2,621 |
| BMI, WC | cg22304262 | 19 | 51979618 | *SLC1A5* | Body/5UTR | -0.0324 | 0.0148 | 2.85E-02 | 2,619 | -0.0987 | 0.0295 | 8.06E-04 | 2,618 | -0.0673 | 0.0191 | 4.26E-04 | 2,619 |
| BMI | cg21766592 | 19 | 51979906 | *SLC1A5* | 1stExon/5UTR/Body | -0.0104 | 0.0147 | 4.82E-01 | 2,623 | -0.0380 | 0.0294 | 1.96E-01 | 2,622 | -0.0201 | 0.0191 | 2.93E-01 | 2,623 |
| BMI | cg03218374 | 20 | 844981 | *ANGPT4* | TSS200 | 0.0172 | 0.0116 | 1.40E-01 | 2,623 | 0.0557 | 0.0232 | 1.64E-02 | 2,622 | 0.0137 | 0.0151 | 3.63E-01 | 2,623 |
| BMI, WC | cg00916899 | 20 | 35378346 | *MANBAL* | 3UTR/3UTR | 0.0236 | 0.0113 | 3.72E-02 | 2,622 | 0.0672 | 0.0226 | 2.92E-03 | 2,621 | 0.0461 | 0.0147 | 1.67E-03 | 2,622 |
| BMI | cg18217136 | 20 | 35591065 | *BLCAP* | TSS1500 | 0.0347 | 0.0268 | 1.97E-01 | 2,623 | 0.1370 | 0.0535 | 1.05E-02 | 2,622 | 0.0912 | 0.0347 | 8.66E-03 | 2,623 |
| BMI | cg24403644 | 20 | 42008038 | *TOX2* | 5UTR/Body/1stExon | 0.0594 | 0.0237 | 1.23E-02 | 2,623 | 0.1412 | 0.0474 | 2.89E-03 | 2,622 | 0.0936 | 0.0307 | 2.31E-03 | 2,623 |
| BMI, WC | cg07950000 | 21 | 30234204 | *GRIK1* | TSS200 | 0.0028 | 0.0137 | 8.40E-01 | 2,623 | 0.0386 | 0.0274 | 1.58E-01 | 2,622 | 0.0312 | 0.0178 | 7.85E-02 | 2,623 |
| BMI | cg08309687 | 21 | 34242466 | | | -0.0295 | 0.0103 | 4.35E-03 | 2,622 | -0.0876 | 0.0206 | **2.12E-05** | 2,621 | -0.0332 | 0.0134 | 1.32E-02 | 2,622 |
| BMI | cg10192877 | 21 | 42514759 | *ABCG1* | Body/5UTR | 0.1692 | 0.0386 | **1.16E-05** | 2,622 | 0.3954 | 0.0770 | **2.80E-07** | 2,621 | 0.1925 | 0.0500 | **1.19E-04** | 2,622 |
| BMI | cg27243685 | 21 | 42515435 | *ABCG1* | Body/5UTR | 0.0798 | 0.0182 | **1.13E-05** | 2,623 | 0.1503 | 0.0363 | **3.48E-05** | 2,622 | 0.0699 | 0.0236 | 3.06E-03 | 2,623 |
| BMI | cg01881899 | 21 | 42525773 | *ABCG1* | Body | 0.0490 | 0.0243 | 4.36E-02 | 2,619 | 0.2006 | 0.0483 | **3.33E-05** | 2,618 | 0.0799 | 0.0314 | 1.11E-02 | 2,619 |
| BMI | cg00222799 | 21 | 42528533 | *ABCG1* | Body | 0.0402 | 0.0141 | 4.30E-03 | 2,623 | 0.0955 | 0.0281 | 6.84E-04 | 2,622 | 0.0337 | 0.0183 | 6.54E-02 | 2,623 |
| BMI, WC | cg06500161 | 21 | 42529656 | *ABCG1* | Body | 0.0961 | 0.0159 | **1.42E-09** | 2,623 | 0.2279 | 0.0316 | **5.64E-13** | 2,622 | 0.1009 | 0.0206 | **9.67E-07** | 2,623 |
| BMI | cg23285465 | 21 | 45184872 | *C21orf67;C21orf70* | TSS1500;Body | 0.0988 | 0.0911 | 2.78E-01 | 2,617 | 0.0457 | 0.1822 | 8.02E-01 | 2,616 | 0.0907 | 0.1180 | 4.42E-01 | 2,617 |
| BMI | cg26354221 | 22 | 23152802 | *ADORA2A* | TSS1500 | 0.0117 | 0.0311 | 7.06E-01 | 2,623 | 0.0129 | 0.0622 | 8.36E-01 | 2,622 | 0.0401 | 0.0403 | 3.20E-01 | 2,623 |
| BMI, Childhood/Young Adult Obesity | cg08548559 | 22 | 30016097 | *PIK3IP1* | Body | -0.0315 | 0.0094 | 7.56E-04 | 2,615 | -0.0865 | 0.0187 | **3.56E-06** | 2,614 | -0.0470 | 0.0121 | **1.04E-04** | 2,615 |

| | | | | | | | | | | | | | | | | | |
|---|---|---|---|---|---|---|---|---|---|---|---|---|---|---|---|---|---|
| BMI | cg27115863 | 22 | 36251586 | | | -0.0232 | 0.0117 | 4.82E-02 | 2,623 | -0.0570 | 0.0234 | 1.48E-02 | 2,622 | -0.0398 | 0.0152 | 8.76E-03 | 2,623 |
| BMI | cg20496314 | 22 | 38089810 | *SYNGR1* | TSS1500/Body | 0.0087 | 0.0073 | 2.31E-01 | 2,619 | 0.0305 | 0.0146 | 3.60E-02 | 2,618 | 0.0020 | 0.0095 | 8.32E-01 | 2,619 |
| BMI | cg06397161 | 22 | 38090005 | *SYNGR1* | Body/TSS200 | 0.0043 | 0.0056 | 4.40E-01 | 2,623 | 0.0180 | 0.0112 | 1.09E-01 | 2,622 | -0.0047 | 0.0073 | 5.23E-01 | 2,623 |
| BMI | cg22650271 | 22 | 38090111 | *SYNGR1* | Body/TSS200 | 0.0171 | 0.0133 | 2.00E-01 | 2,623 | 0.0584 | 0.0266 | 2.79E-02 | 2,622 | -0.0002 | 0.0173 | 9.89E-01 | 2,623 |
| BMI | cg03318904 | 22 | 38131468 | *MAP3K7IP1* | Body | 0.0272 | 0.0190 | 1.53E-01 | 2,618 | 0.1242 | 0.0380 | 1.08E-03 | 2,617 | 0.0643 | 0.0246 | 9.03E-03 | 2,618 |
| BMI, Childhood/Young Adult Obesity | cg09349128 | 22 | 48713990 | | | 0.0085 | 0.0135 | 5.31E-01 | 2,623 | 0.0138 | 0.0270 | 6.10E-01 | 2,622 | -0.0079 | 0.0175 | 6.51E-01 | 2,623 |
| BMI | cg09182678 | 22 | 48714715 | | | -0.0356 | 0.0253 | 1.59E-01 | 2,623 | -0.0979 | 0.0505 | 5.24E-02 | 2,622 | -0.0766 | 0.0327 | 1.92E-02 | 2,623 |
